# Classic and introgressed selective sweeps shape mimicry loci across a butterfly adaptive radiation

**DOI:** 10.1101/685685

**Authors:** Markus Moest, Steven M. Van Belleghem, Jennifer E. James, Camilo Salazar, Simon H. Martin, Sarah L. Barker, Gilson R. P. Moreira, Claire Mérot, Mathieu Joron, Nicola J. Nadeau, Florian M. Steiner, Chris D. Jiggins

## Abstract

Natural selection leaves distinct signatures in the genome that can reveal the targets and history of adaptive evolution. By analysing high-coverage genome sequence data from four major colour pattern loci sampled from nearly 600 individuals in 53 populations, we show pervasive selection on wing patterns across the *Heliconius* adaptive radiation. The strongest signatures correspond to loci with the greatest phenotypic effects, consistent with visual selection by predators, and are found in colour patterns with geographically restricted distributions. These recent sweeps are similar between co-mimics and indicate colour pattern turn-over events despite strong stabilizing selection. Using simulations we compare sweep signatures expected under classic hard sweeps with those resulting from adaptive introgression, an important aspect of mimicry evolution in *Heliconius*. Simulated recipient populations show a distinct ‘volcano’ pattern with peaks of increased genetic diversity around the selected target, consistent with patterns found in some populations. Our genomic data provide unprecedented insights into the recent history of selection across the *Heliconius* adaptive radiation.

## Introduction

Identifying targets of selection and reconstructing their evolutionary history is central to understanding how populations adapt ^1–3^. In particular, genome sequences contain a rich source of information about past events in natural populations. The action of recent positive selection can leave a distinct signature known as a ‘selective sweep’, which provides information on the genomic location of targets of positive selection and the timing and strength of selection ^4,5^. While many classic examples of selective sweeps have been found in domesticated populations, such as maize ^6^, chicken ^7^, and cattle ^8^, or in humans ^9^, increasingly natural populations are also studied. Using genomic data, these latter studies can reveal the genetic architecture and evolutionary history of ecologically relevant traits ^10–13^ and provide insights into the action of natural selection by complementing field and experimental studies ^14–16^. However, to date few molecular studies of natural populations have used broad sampling across adaptive radiations with varying selection pressures and sources of adaptive variation for the same trait. Such studies will allow the investigation of both complexity and general mechanisms of natural selection in the wild at the genotypic level, especially where there is *a priori* information on the agents and targets of selection.

Positive selection can rapidly change allele frequencies leaving detectable signatures in a genome. These signals can be traced over ecological and evolutionary time scales, during which they are gradually eroded by new mutations and recombination ^1^. However, the observed patterns will depend on the sources and frequency of genetic variation upon which selection acts ^5^. For example, a classic ‘hard sweep’ due to selection on a single, novel beneficial mutation ^4^ or a very rare allele from standing variation ^17^, is distinct from a ‘soft sweep’ due to selection on standing variation already present at an appreciable frequency ^17–20^ or recurrent mutations ^21,22^. Less well studied in the context of selective sweeps is the possibility that a new variant is introduced by gene flow from a related population or distinct species. Accumulating evidence suggests that this re-use of ancient variants is far more common than was previously envisioned ^23–26^. However, the sweep signatures created by selection on one or several introgressed and therefore divergent haplotypes and the effect of migration rate on these signatures are largely unexplored (but see ^27^).

Mimicry systems provide some of the best examples of natural selection and adaptation and, thus, exceptional opportunities to study selective sweeps. In the unpalatable *Heliconius* butterflies, mimicry of wing patterns is advantageous as resemblance to a common, well-protected pattern confers protection from predator attacks on individuals. The majority of pattern diversity seen in this group is controlled by a surprisingly simple genetic system, involving allelic variation at just four major effect loci. These loci comprise the transcription factors, *optix* ^28^ and *aristaless*, which comes in two tandem copies *al1* and *al2* ^29^, a signalling ligand, *WntA* ^30^, and a gene in a family of cell cycle regulators whose exact function remains unclear, *cortex* ^31^. A complex series of regulatory variants at each of these loci is found in different combinations across populations and species, leading to great diversity of wing patterns. Novelty is generated both by mutation, and through introgression and shuffling of existing *cis*-regulatory elements (CREs) which generate new pattern combinations ^32–34^. Moreover, adaptive sharing of mimicry colour patterns has been demonstrated across many species and populations within the *H. melpomene* and *H. erato* clade ^32–39^.

*Heliconius* colour patterns are known to be subject to remarkably strong natural selection in wild populations, which has been demonstrated through pattern manipulations ^40^, reciprocal transplants across a hybrid zone ^41^, reciprocal transfers between different co-mimic communities ^42^, and artificial models. In all cases, estimates of selection strength were high with *s* = 0.52-0.64 (Table 1). Indirect estimates of selection strength from hybrid zones generated similarly high values with *s* = 0.23 for each of three colour pattern loci, *optix, cortex*, and *WntA*, in *H. erato* and *s* = 0.25 for *optix* and *cortex* in *H. melpomene* ^43–47^ (see Table 1 for details).

**Table 1:** Direct and indirect estimates of selection on colour pattern loci. Combined estimates are integrating the effect of all loci involved in warning colouration. Regions/modules associated with *optix*: D, B; with *cortex*: Cr, Yb, N; with *WntA*: Sd, Ac; with *aristaless*: K.

| Species | Targets of selection under consideration | Estimated selection coefficient ($s$) | Method | Source |
| --- | --- | --- | --- | --- |
| *H. erato* | *optix* (red band) | $s_D = 0.22$ | Pattern manipulation, survival and bird attack rate | Benson<sup>40</sup> ($s$ estimate calculated in Mallet *et al.*<sup>119</sup>) |
| *H. erato* | *optix/cortex/WntA* | combined $s = 0.52$<br>avg. per locus $s = 0.17$ | Reciprocal transplants, survival | Mallet and Barton<sup>41</sup> |
| *H. erato* | *optix/cortex/WntA* | $s_D = 0.33$<br>$s_{Cr} = 0.15$<br>$s_{Sd} = 0.15$ | Reciprocal transplants, survival | Mallet *et al.*<sup>119</sup> |
| *H. erato*<br>*H. melpomene* | *optix/cortex/WntA*<br>*optix/cortex* | avg. per locus $s = 0.23$<br>avg. per locus $s = 0.25$ | Cline and LD analysis in a hybrid zone | Mallet *et al.*<sup>45</sup> |
| *H. cydno* (polymorphic mimic)<br>*H. sapho* (model)<br>*H. eleuchia* (model) | *aristaless* | $s = 0.64$ | Reciprocal transplant of polymorphic *H. cydno* | Kapan<sup>42</sup> |
| *H. erato* | *optix/cortex/WntA* | avg. per locus $s = 0.22$<br>$s_D = 0.38$<br>$s_{Cr} = 0.17$<br>$s_{Sd} = 0.15$ | Cline and LD analysis in a hybrid zone | Rosser *et al.*<sup>46</sup> |
| *H. melpomene* | *optix/cortex* | avg. per locus $s = 0.31$<br>$s_D = s_{Yb} = s_N = 0.31$<br>$s_B = 0.19/0.15$ | | |
| *H. erato*<br>*H. melpomene* | *optix/WntA*<br>*optix/WntA* | $s_D = 0.15$<br>$s_{Sd} = 0.04$<br>$s_D = 0.27$<br>$s_{Ac} = 0.04$ | Cline analysis in a hybrid zone | Salazar<sup>47</sup> |

Although colour pattern loci in *Heliconius* are well studied, and their adaptive significance is apparent, the impact of selection at the molecular level has never been estimated in natural populations. Genetic studies have shown that populations often cluster by phenotype rather than geography at colour pattern loci ^33,48,49^, but these approaches may not detect recent adaptive changes. For example, closely related populations show peaks of high differentiation at colour pattern loci ^50,51^, but previous studies did not reveal strong sweep signatures ^52–54^, and more recent genomic analysis showed only weak evidence for reduced heterozygosity and enhanced linkage disequilibrium ^49^. However, these studies have used either few amplicons or genomic data with small sample sizes, and therefore potentially had little power to detect selective sweep signatures.

Here, we obtain a large genomic data set across the *H. melpomene* radiation, featuring both high coverage and large sample size, and combine simulations with population genomic analysis to investigate natural selection at four main colour pattern loci. We also use simulations to delimit the age of the sweeps and to derive expected patterns for introgressed sweeps in *Heliconius*. Our empirical dataset covers almost the entire biogeographic range of an adaptive radiation and demonstrates clear signatures of selective sweeps across many populations. However, many widespread colour patterns show only modest signals of selective sweeps, with the strongest sweeps found in populations with geographically restricted patterns, suggesting recent and strong selection. For adaptive introgression, our simulations demonstrate that the signals have distinct shapes, are strongly affected by effective migration rates, and are more challenging to detect. Nevertheless, we identify sweep signatures among populations with known colour pattern introgression. Moreover, we identify new putative targets of selection around colour pattern genes in some populations. Finally, we also analyse genomic data from *H. erato* populations, representing a distinct radiation of similar wing pattern forms, and find evidence for parallel evolution between co-mimetic butterfly species.

## Results

### Phylogeography and demography of the *Heliconius melpomene* clade

We obtained *ca*. 5.2 Mb of sequence distributed across 8 chromosomes from 473 individuals from 39 populations representing 10 species from the *H. melpomene* clade (Supplementary Table 1). Phylogenetic reconstructions confirmed that *Heliconius cydno* populations, with the sole exception of *H. c. cordula* found east of the Andes and in the Magdalena Valley, and *H. timareta* populations from east of the Andes cluster as separate lineages from the *H. melpomene* clade (Fig. 1B & D). Phylogenetic inferences including all sequenced regions agreed with previous multi-locus phylogenies, in which *H. cydno* and *H. timareta* form a sister clade to *H. melpomene* (Fig. 1D, Supplementary Fig. 1) ^38,55^. The tree built using only neutral background data (i.e. regions *a priori* not suspected to be under mimicry selection, see Methods) largely clustered populations according to geography, i.e. *H. cydno* with western *H. melpomene* and *H. timareta* with eastern *H. melpomene* subspecies (Fig. 1B & D). The neutral topology is consistent with ongoing gene flow between sympatric populations resulting in highly heterogeneous relatedness patterns along the genome ^56,57^. Six out of nine individuals with the dennis-ray pattern, sampled from the *H. melpomene vicina* population in the Colombian Amazon (Fig. 1A & C), consistently clustered within *H. timareta*. This suggests the presence of a lowland population of *H. timareta* considerably further from the Andes than has been detected previously, hereafter referred to as *H. timareta ssp. nov.* (Colombia). To assess demographic events, which may affect selection tests, we estimated effective population size across time for all populations with whole-genome data (Supplementary Table 1). In line with previous studies ^54,58^ we found that bottlenecks were rare across those populations with the exception of a recent decline in population size in *H. heurippa* and older, moderate dips in *H. besckei* and *H. m. nanna* (Supplementary Fig. 2).

**Figure 1.**
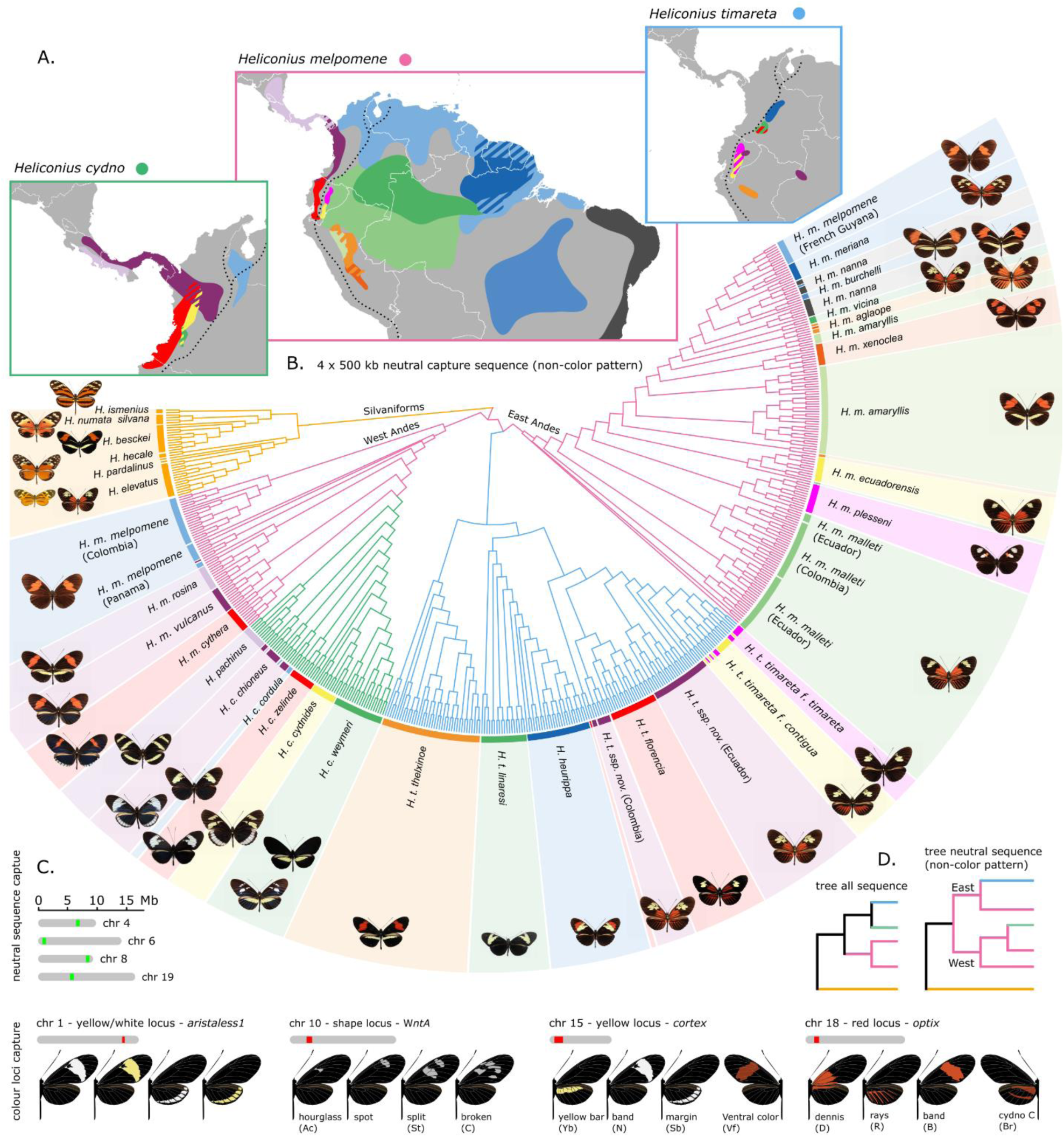
Distribution, phylogenetic relations, major colour pattern loci and sequence capture targets of the *Heliconius melpomene - cydno - timareta* clade species. (**A.**) Broad distributions of the *H. melpomene, H. cydno* and *H. timareta* colour pattern races and species (based on all known sampling localities). Distribution colours match the shadings around the phylogeny and butterfly images in panel B. The dashed line indicates the Andes. Note the distinct clusters formed by individuals sampled from the *H. m. vicina* population. The cluster grouping with *H. timareta* is referred to as *H. timareta ssp. nov.* (Colombia) (**B.**) FastTree cladogram inferred using capture sequence from putatively neutral loci. Colours in the tree indicate the *H. melpomene* (pink), *H. cydno* (green) and *H. timareta* (blue) clades and match the boxes of the distribution maps in panel A. (**C.**) Sequence information was obtained for four putatively neutral regions (green) and four regions to which functional variation has been mapped to a yellow/white colour switch (chr 1), forewing band shape (chr 10), yellow/white fore- and hindwing bars, band margins and ventral colour (chr 15) and red colour pattern elements (chr 18). The various phenotypes controlled by the respective colour pattern loci are depicted. Note that while most phenotypes have descriptive names the red blotch at the base of the forewing was termed ‘dennis’. (**D.**) Phylogenetic relations obtained when building a tree from all captured regions compared to the neutral regions.

### Signatures and limits of detection of classic sweeps assessed by simulations

We used extensive simulations to evaluate selection patterns and our power to detect sweeps in *Heliconius*. In our analysis, we primarily use SweepFinder2 (SF2), which is appropriate for our genomic data as it is robust to demography and able to identify the sweep site (for more details, see Methods). However, to more qualitatively explore patterns of diversity at sites undergoing selection, we also present results observed in Tajima’s *D* at selected sites. The time over which we can expect to detect sweep signals is determined by the time to coalescence, and is thus determined by *N*, the (effective) population size. We therefore here report time since the sweep in generations, scaled by 4*N* ^59^. Sweep signals are expected to decay rapidly due to the joint effects of mutation, recombination, and drift. Indeed, SweepFinder2, which uses the predicted effect of a selective sweep on the local site-frequency spectrum (SFS) to infer the probability and location of sweeps ^60–62^, has low power to detect even hard selective sweeps that occurred over 0.25 (scaled) generations ago and cannot localize sweeps older than 0.4 (scaled) generations ^61^. Consequently, any detected sweep signals in *Heliconius melpomene* are likely under 0.8 million years old, assuming an effective population size of 2 million ^54,63^ and a generation time of 3 months ^64^. As these estimates vary with *N*, the time limit for sweep detection varies among species, from only 0.2 Mya for *H. besckei* (*N* ∼ 0.5 million) to 1.4 Mya for *H. erato* (*N* ∼ 3.5 million). We used simulations to further interpret the empirical signatures of selection and explore the limits of detection (Fig. 2). We initially simulated the case of a hard sweep, such that *s* = 0.5, which is appropriate to the very strong selection pressure experienced by the colour pattern loci in *Heliconius* (Table 1). We found that after a sweep event, Tajima’s *D* values were reduced compared with neutral background levels for a considerable time (even after 0.5 generations, Welch’s t-test, p < 0.01), and the obvious dip around the selected site remained very pronounced for 0.1 generations post-sweep (Fig. 2). SweepFinder2 signals broke down more rapidly (Fig. 2). The magnitude of the CLR peak decreased by an order of magnitude after just 0.1 generations, corresponding to 0.2 Mya for *H. melpomene*, and was not distinguishable from background values after 0.2 generations, i.e. 0.4 Mya in *H. melpomene* (Welch t-test, p = 0.065). Similarly, the estimated strength of selection calculated with SweepFinder2 from our simulations declined rapidly with time.

**Figure 2:**
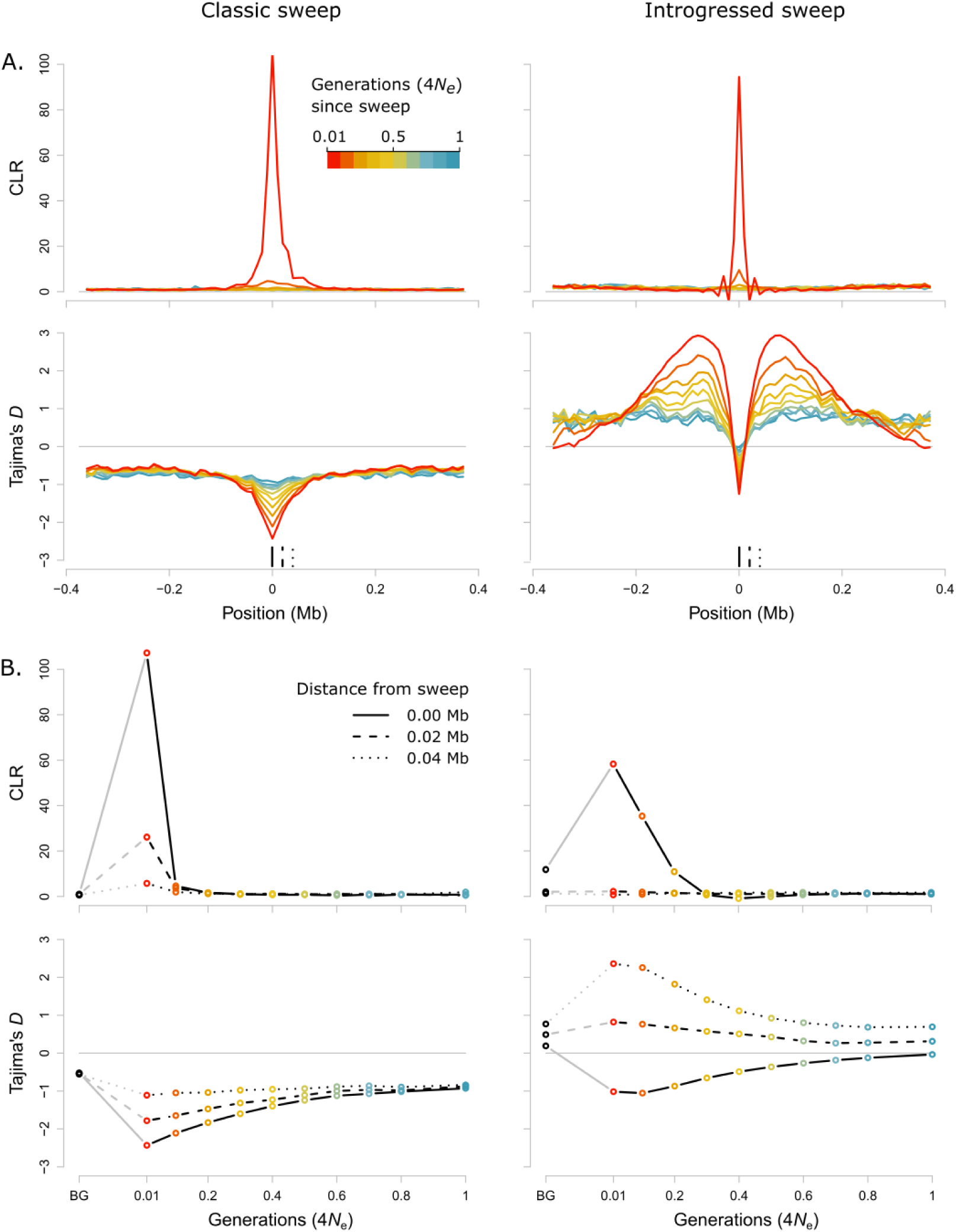
SFS signatures of selection for simulated classic hard sweeps (left) and introgressed sweeps (right). (**A**.) Composite likelihood ratio statistics (CLR, upper panel, ^60,61^) and Tajima’s *D* (lower panel) across a simulated chromosome for different time points (0.01, 0.1, 0.2, 0.3, 0.4, 0.5, 0.6, 0.7, 0.8, and 1 in units of scaled generations, i.e. 4*N* generations) after a classic hard (left) or introgressed (right) sweep (effective migration rate *M* = 0.2). The sweep occurs in the centre of the simulated chromosome. Different colours indicate time since sweep. Full, dashed and dotted vertical black lines in the lower panel indicate positions at different distances from the sweep centre for which time series of CLR and Tajima’s D statistics are depicted (B.) in the same style. (**B.**) CLR (upper panel) and Tajima’s *D* (lower panel) statistics over time at three positions relative to the sweep centre as shown in (A.). Also shown are neutral background values, BG, calculated over neutral simulations, either without migration (left hand panels, for classic sweeps) or with migration at *M* = 0.2 (right hand panels, for introgressed sweeps). Time is given in units of scaled generations.

### Signatures and limits of detection of introgressed sweeps assessed by simulations

We extended our simulations to explore the expected SFS signature left by an allele undergoing adaptive introgression. We did this by simulating a second population which exchanged migrants with the first, leading to an introgressed sweep in the second population. Estimates of effective migration rates (*M*) between hybridising *Heliconius* species vary from 0.08 to 10 migrants per generation ^65–67^. Here, we report in detail our results for *M* = 0.2, but we also investigated values of *M* from 0.002 to 200 (Supplementary Fig. 3). Adaptive introgression produces a highly distinctive SFS signature. At and very close to the selected site itself there was a reduction in diversity and an excess of rare alleles, similar to the pattern observed for a classic sweep. However, this reduction was narrow, and flanked by broad genomic regions with high diversity and an excess of intermediate frequency variants. This is due to variants that have hitch-hiked into the recipient population along with the beneficial variant, and subsequently recombined before reaching fixation ^20,27^. The overall SFS signature covered a considerably wider genomic area than that of a classic sweep (Fig. 2).

In simulations where values of *M* are under 0.2, the signature we observe at the sweep site itself was very similar to that for a classical sweep, and we could detect it for a similar length of time: the distribution of Tajima’s *D* values was not significantly different from those calculated over neutral regions after 1 (scaled) generation (Welch’s t-test p = 0.08). Therefore, SweepFinder2 could detect introgressed sweeps, however, it detected only the central region of lowered diversity, producing a high but very narrow CLR peak at the sweep site itself; this contrasts with the peaks for classic selective sweeps, which extended over a wider genomic area (Fig. 2). The distribution of CLR values at the sweep site was significantly different from values calculated over neutral regions for up to 0.1 generations after the sweep (p = 0.0041). However, as for a classical sweep, the magnitude of the peak decreased rapidly. The magnitude of the reduction at the sweep site itself was also strongly affected by migration rate. As *M* increases, the central reduction in diversity becomes less pronounced, representing an increasingly ‘soft’ introgressed sweep (Supplementary Fig. 3) ^21,68^. Therefore, detecting introgressed sweeps from this central region will be increasingly difficult with increasing *M*. However, for values of *M* below 2, varying *M* had little effect on the regions of increased diversity and excess of intermediate frequency variants that flank the sweep locus (Supplementary Fig. 3).

### Strong signatures of selection across *Heliconius* colour pattern regions

In our empirical data, SweepFinder2 found strong support for positive selection acting across multiple populations and species for all four colour pattern loci (Fig. 3). In contrast, our background regions as well as regions flanking the colour pattern associated loci showed little evidence of sweeps, apart from a few isolated examples (Supplementary Fig. 4). These results therefore provide perhaps the first unbiased evidence to support the long-standing assertion that wing patterning loci are among the most strongly selected loci in the genome and have a distinctive evolutionary history ^69^.

**Figure 3.**
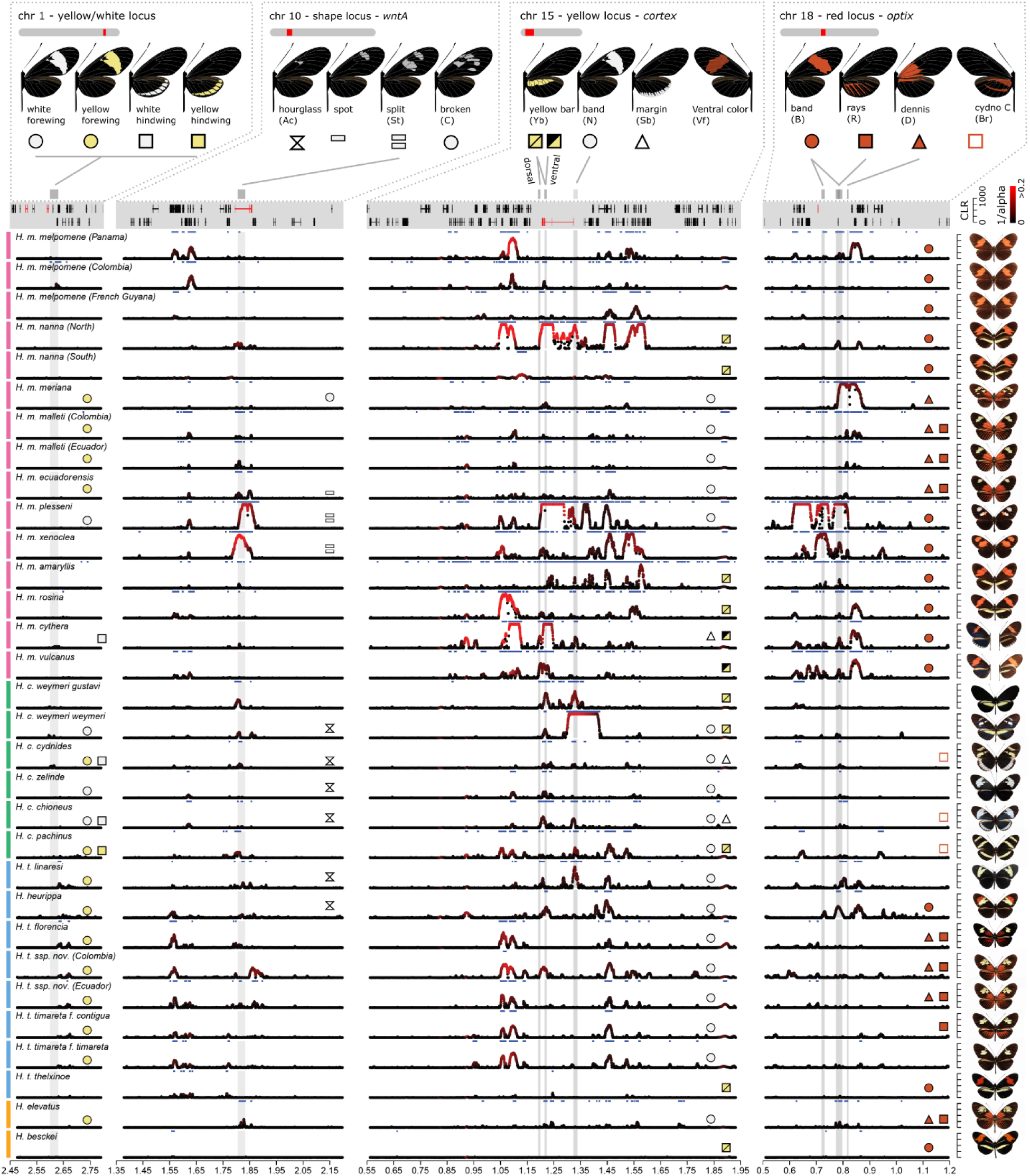
Signature of selection across colour pattern regions in the *H. melpomene*-clade. The regions containing the tandem copies of *aristaless, al1* and *al2, WntA, cortex*, and *optix* (left to right) are depicted. Colour pattern genes are annotated in red in the gene annotation panel. On the y-axis Sweepfinder2’s composite likelihood ratio statistics (CLR) is shown (peaks are capped at CLR = 1,000). The colour gradient indicates the estimated intensity of selection (black…high *α* values, weak selection; red…low *α* values, strong selection). Grey shadings indicate annotated colour pattern regulatory elements (CREs) ^31,32,34,72^ (Supplementary Fig. 7, 12-14). Blue horizontal bars indicate regions with CLR values above threshold. Top panel shows colour pattern phenotypes and symbols indicate distinct colour pattern elements and their presence is annotated in population panels. Note that the yellow hindwing bar controlled by the *cortex* region can be expressed on the dorsal and ventral side (yellow/yellow square symbol) or on the ventral side only (black/yellow square symbol) ^32^. Moreover, the actual shape of the forewing band can depend on the allelic state of *WntA*. Full, gray lines connect colour pattern elements with annotated CREs. Phenotypes are depicted on the right. *H. m. vicina, H. m. aglaope, H. m. burchelli* and *H. c. cordula* are not shown due to low sample size.

Broadly, signals of selection were stronger and more widespread near *cortex* and *optix*, and weaker near *WntA* and *aristaless*. For example, all 31 populations showed sweep signals above threshold near *cortex*, 26 near *optix*, 24 near *WntA*, albeit less pronounced in most cases, and only 7 near *aristaless* (Fig. 3; Supplementary Tables 2-5). A similar pattern was reflected in our estimates for strength of selection (*s*) calculated from *α* estimates (Table 2) with the highest selection strength in colour pattern regions being *s* = 0.141 for *cortex* (*H. m. nanna*), *s* = 0.036 for *optix* (*H. m. plesseni), s* = 0.049 for *WntA* (*H. m. xenoclea*) and *s* = 0.01 (*H. t. florencia*) for *aristaless* (*H. t. florencia*). These patterns are broadly concordant with the expected phenotypic effects of these loci. For example, in *H. cydno* which has primarily yellow and/or white patterns controlled by *cortex*, significant peaks were mostly found at this locus, while in *H. melpomene* which has red, yellow and white patterns, strong signals were seen at both *cortex* and *optix*. Consistently, a lower strength of selection was found for *aristaless* (*s* < 0.01), which controls a putatively less salient ^70^ modification of pale patterns from yellow to white.

**Table 2:**
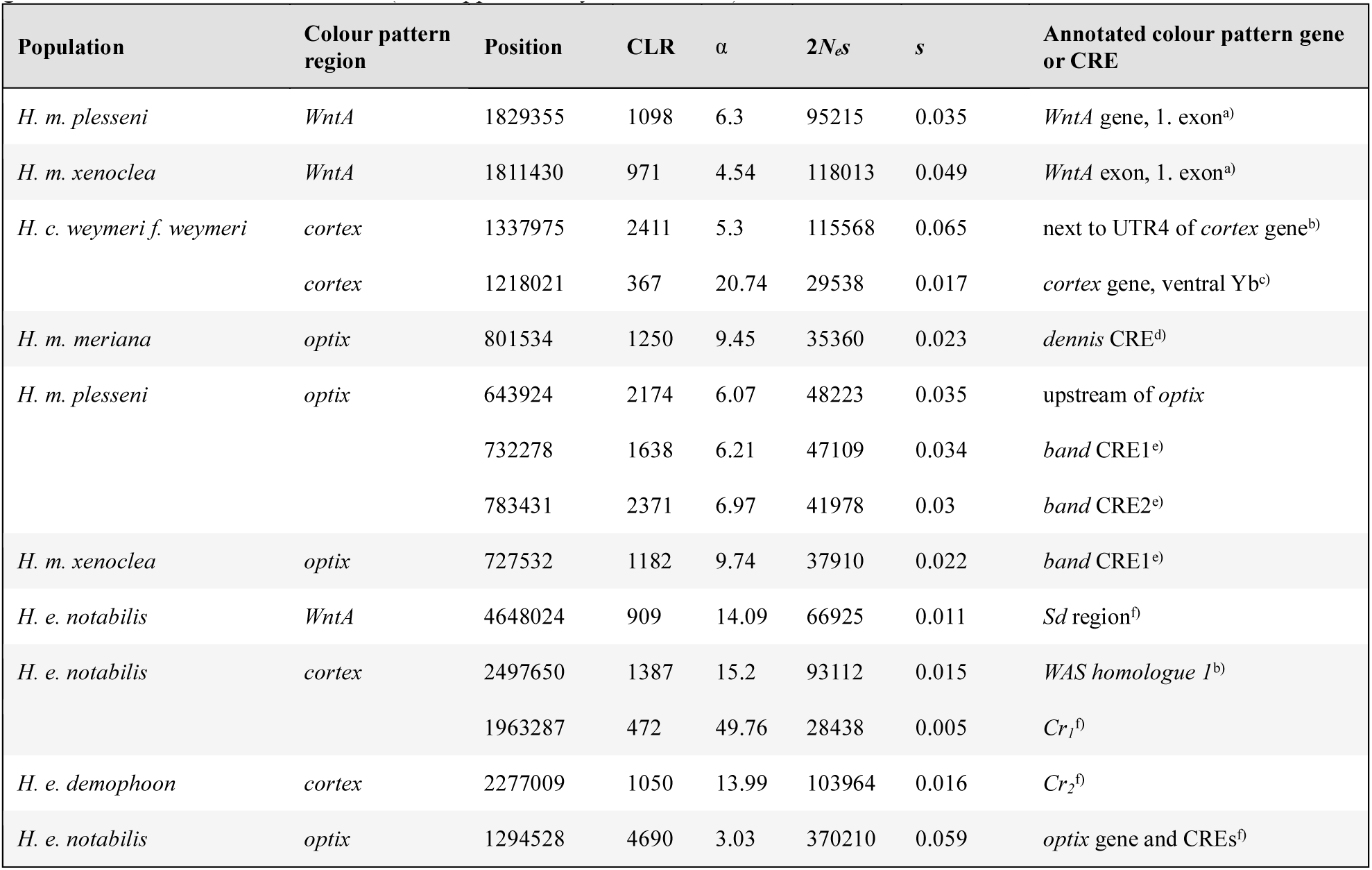
Position, composite likelihood-ratio statistics (CLR), and estimates for strength of selection (*α*, 2*Nes*, and *s*) for populations and sweeps discussed in detail. Annotated colour pattern genes and CREs that overlap with peaks are given. ^a)^Mazo-Vargas *et al*. ^120, b)^Nadeau *et al*. ^31, c)^Enciso-Romero *et al.* ^32, d)^Wallbank *et al*. ^34, e)^Hanly ^72, f)^Van Belleghem *et al*. ^33^. Positions are given in Hmel2 scaffold coordinates (see Supplementary Tables 2 & 4).

There were also differences seen across the sampled populations. Widely distributed colour patterns (e.g. *H. m. melpomene* and *H. m. malleti*) tended to show only modest evidence for selective sweeps (Fig. 3, Supplementary Fig. 5). Comparisons with our simulated data nonetheless suggest selective events that occurred no more than 400,000 years ago. Geographically localised patterns showed much stronger signatures, likely reflecting sweeps within the last 100,000 years (Fig. 4, Table 2). For example, *H. m. plesseni* is exclusively found in the upper Pastaza valley in Ecuador and shows a unique split red-white forewing band (Fig. 1 and 4). This population showed strong selection at three colour pattern regions, *optix, cortex*, and *WntA*, suggesting recent selection acting on the entire pattern (*s*_*cortex*_ = 0.074, *s*_*WntA*_ = 0.035, and *s*_*optix*_ = 0.035), and patterns of both nucleotide diversity and Tajima’s *D* are consistent with strong classic sweeps (Fig. 3 & 4, Supplementary Fig. 5, Supplementary Table 2). *Heliconius m. xenoclea*, also found on the Eastern slopes of the Andes but further south in Peru, shows the same split forewing band controlled by *WntA* and again a very strong selection signal at this locus (*s*_*WntA*_ = 0.049), as well as weaker signatures at *cortex* (*s*_*cortex*_ = 0.04) and *optix* (*s*_*optix*_ = 0.022) (Fig. 3, Supplementary Fig. 5, Supplementary Table 2). The clear signatures of recent and strong selection pressure perhaps indicate that the split forewing band is a novel and highly salient signal. Additionally, *H. m. meriana* from the Guiana shield revealed a striking signature of selection at *optix* (*s*_*optix*_=0.023). Its dennis-only pattern (see Fig. 4) has previously been shown to have arisen through recombination between adjacent dennis and ray regulatory modules at *optix*, and the signature of selection at this locus, which encompasses both of these regulatory modules, implies a recent sweep of this recombinant allele ^34^ (Fig. 3 & 4, Supplementary Fig. 5, Supplementary Table 2).

**Figure 4.**
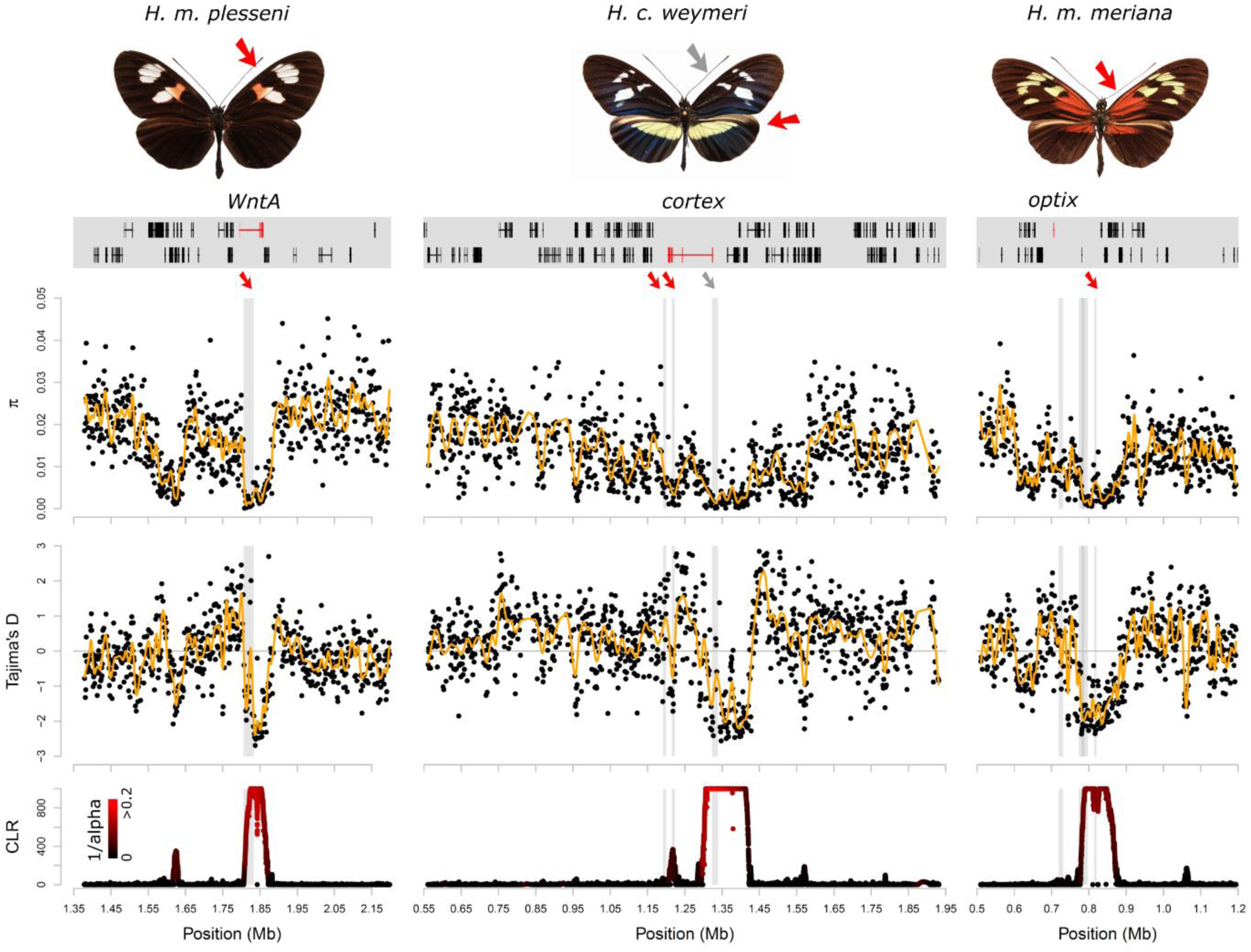
Selected examples of sweeps. The three examples show the split forewing band (*WntA*) in *H. m. plessini*, the yellow and white patterns (*cortex*) in *H. cydno weymeri f. weymeri* and the red dennis patch (*optix*) in *H. m. meriana* (left to right). The respective colour pattern elements are indicated with red and grey arrows. Colour patterns and gene annotations in the colour pattern regions are depicted in the top panel. Colour pattern genes are annotated in red. Nucleotide diversity *π*, Tajima’s *D* and SweepFinder2’s composite likelihood ratio statistics (CLR, peaks are capped at CLR = 1,000) show the signatures of a selective sweep (bottom panels). Loess smoother lines are depicted in yellow. The colour gradient in the CLR panel indicates the estimated intensity of selection (black…high *α* values, weak selection; red…low *α* values, strong selection). Grey shadings indicate annotated CREs and red and grey arrows depict associations with the respective colour pattern elements in the in the *H melpomene*-clade.

In light of our simulations of introgressed sweeps, there were cases in our data where previously documented adaptive introgression events showed signatures characteristic of introgressed sweeps. The hindwing yellow bar pattern was suggested to have introgressed from *H. melpomene* into *H. c. weymeri*, and then back again into the races *H. m. vulcanus* and *H. m. cythera* ^32^. Accordingly, we found narrow SweepFinder2 peaks and an increase in Tajima’s *D* at surrounding sites at these modules in the *cortex* region in *H. m. cythera, H. m. vulcanus* and *H. c. weymeri*, consistent with introgressed sweeps (Fig. 3, Supplementary Fig. 5). *Heliconius c. weymeri f. weymeri* also had a second, striking signature further upstream more typical of a classic sweep (Fig. 3 & 4), at a region associated with the yellow forewing band in *H. melpomene* and *H. timareta* ^31^. This is consistent with evidence for a role of *cortex* in controlling the white forewing band in *H. cydno* ^71^ and the presence of this band in the *weymeri* morph, which could therefore represent a recent evolutionary innovation. Other loci previously implicated as having introgressed include *optix* in *H. heurippa* and *H. elevatus*, which both showed signals coinciding with regions previously associated with the respective phenotypes ^34,72^. In contrast, there was a lack of clear introgressed sweep signals in dennis-ray *H. timareta*, which is one of the best documented examples of introgression. This could be explained by the age of the sweeps and/or high rates of migration, which our simulations show can reduce the sweep signal in the recipient population (Supplementary Fig. 3).

### Novel targets of selection in colour pattern regions

Many of the signals of selection we detected overlap with previously identified regulatory regions associated with colour pattern variation. However, our analysis also found additional nearby regions showing consistent signals of selection that may also be involved in colour pattern evolution (Fig. 3, Supplementary Fig. 6). For example, in the first intron of the *WntA* gene, we found a consistent signal across several *H. melpomene, H. timareta* and *H. cydno* populations (Supplementary Fig. 6B). Within this region (Hmel210004:1806000-1833000), phylogenetic clustering of the two split forewing band races *H. m. plesseni* and *H. m. xenoclea*, indicates a common origin of the split band in these currently disjunct populations (Supplementary Fig. 7). Additionally, two strong selection signatures are frequently found in a region ca. 200 kb upstream of *WntA* (Supplementary Fig. 6B; Hmel210004:1550000-1650000), which suggests additional loci involved in colour pattern regulation.

Near *cortex*, selection signatures at closely linked genes support findings from previous studies. Several populations show distinct peaks up- and downstream of *cortex* and broadly coincide with a wider region, possibly containing several genes involved in colour pattern regulation ^31^ (Supplementary Fig. 6C). Multiple peaks are located upstream of *cortex* within an array of genes that all showed significant associations with yellow colour pattern variation ^31^ (Supplementary Table 6). A particular concentration of signals fell near *LMTK1* (HMEL000033; Hmel215006:1,418,342-1,464,802) and close to *washout*, which previously showed a strong association with the yellow forewing band ^31^. Likewise, selection signals clustered downstream of *cortex* in a region containing additional candidate genes identified previously (Supplementary Table 6). In the *optix* region, consistent signals across several populations indicated that several as yet uncharacterized elements may be under mimicry selection. Intriguingly, the *kinesin* gene, which shows an association of expression with the red forewing band ^73,74^, was among these (Supplementary Fig. 6D).

### Parallel selective sweep signatures between mimetic species

There has been considerable interest in whether the *H. erato* and *H. melpomene* co-mimics have co-diverged and simultaneously converged onto the same colour pattern ^75–77^ or whether one species evolved towards diverse phenotypes of the other, i.e. advergence ^48,78–80^. Homologous genes control corresponding phenotypes ^28,31,81,82^ but there is no allele sharing between the *melpomene*- and *erato*-clade ^48,49^. We used published genomic data for *H. erato* (Van Belleghem *et al.* 2017) (Supplementary Table 7) to obtain 8.9 Mb of sequence homologous to the regions studied in the *H. melpomene*-clade for 103 individuals from 13 populations and 3 species in the *H. erato* radiation, and scanned for selective sweeps. Generally, a comparison of the location of selection peaks between *H. melpomene* and *H. erato* across several co-mimetic races suggests a rather simple and concordant regulatory architecture in the two species at the *WntA* locus. However, in the *cortex* and *optix* regions, this architecture appears to be more complex and differs more strongly between the two clades (Fig. 5, Supplementary Fig. 6 & 11).

**Figure 5.**
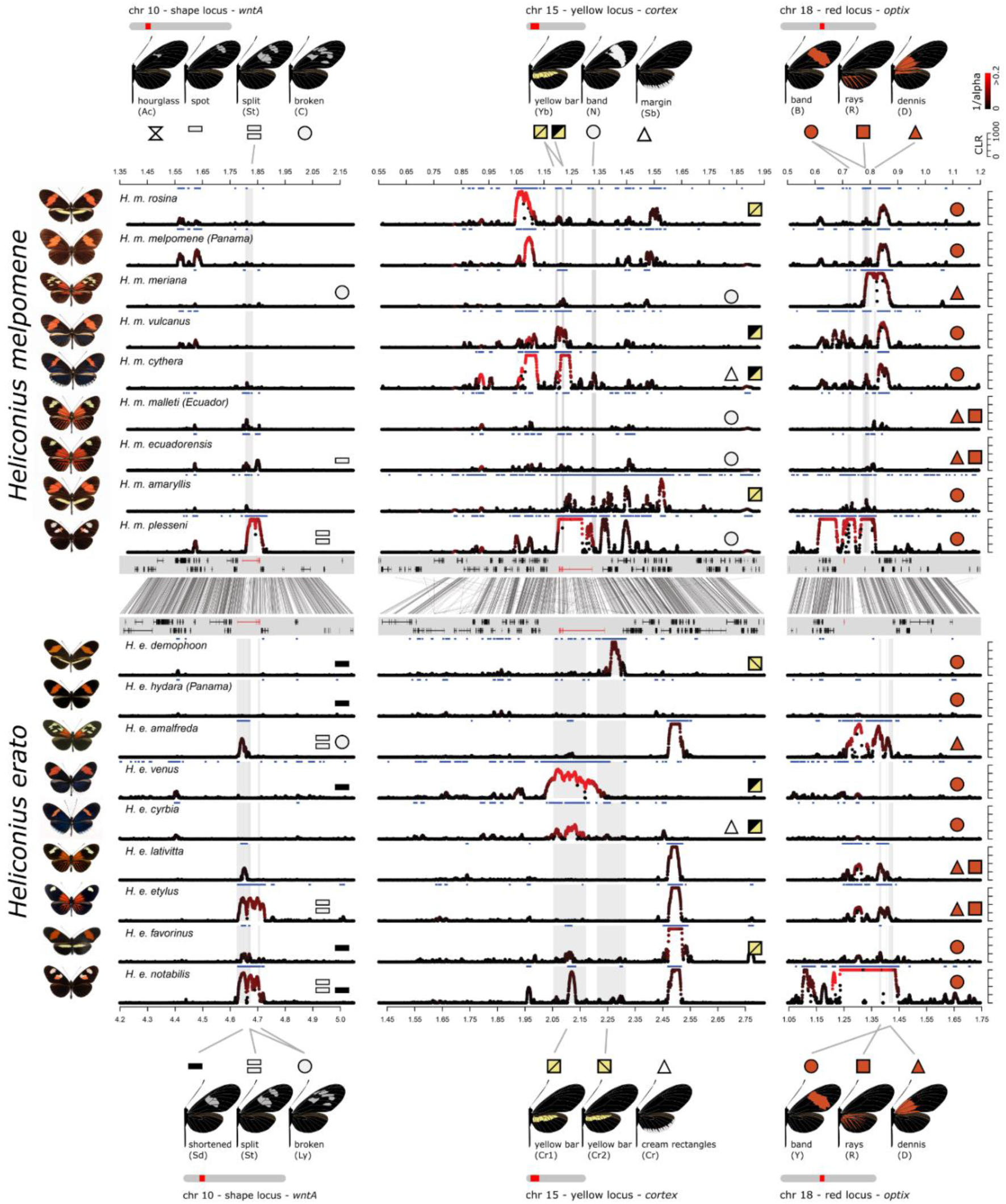
Signatures of selection in the co-mimic populations of *H. melpomene* (upper panels) and *H. erato* (lower panels). The regions containing *WntA, cortex*, and *optix* are shown (left to right). Co-mimics in *H. melpomene* and *H. erato* are depicted in the same order with phenotypes on the left. The y-axis indicates composite likelihood ratio statistics (CLR) across each region (peaks are capped at CLR = 1,000). The colour gradient indicates the estimated intensity of selection (black…high *α* values, weak selection; red…low *α* values, strong selection). Grey shadings indicate annotated colour pattern regulatory elements (CREs ^31,32,34,72^ (Supplementary Fig. 12-14) and blue horizontal bars indicate regions with CLR statistics above threshold. The central panel shows an alignment of the respective regions in *H. melpomene* and *H. erato* and gene annotations with colour pattern genes in red. Top and bottom panel show colour pattern phenotypes and symbols indicate distinct colour pattern elements and their presence in each population panel. Note that the yellow hindwing bar controlled by the *cortex* region can be expressed on the dorsal and ventral side (yellow/yellow square symbol) or on the ventral side only (black/yellow square symbol) ^32^. Full, grey lines connect colour pattern elements with annotated CREs. Note that the genetics of the yellow forewing band differs between *H. erato*, in which it involves the *WntA* and *optix* locus, and *H. melpomene*, in which the band is controlled by *cortex* and its shape by *WntA.*

Similar to the *melpomene*-clade radiation, we found strong signatures of selection across the *optix, cortex*, and *WntA* regions (Fig. 5, Supplementary Fig. 8, 9 & 10, Table 2, Supplementary Tables 8-11). Most notably, *H. e. notabilis* from Ecuador showed strong signals of selection at three colour pattern loci (*s*_*optix*_=0.06, *s*_*cortex*_=0.015, *s*_*WntA*_=0.015) similar to its co-mimic *H. m. plesseni* (Table 2). In both cases, selection across the three major loci represented some of the strongest signals in both species. Additionally, *H. e. amalfreda*, co-mimic with the red dennis-only race *H. m. meriana*, showed one of the strongest selection signals at *optix*. This suggests that these phenotypes are recent innovations in both species, consistent with co-divergence. Other geographically localised variants controlled by *WntA* also showed strong signals of selection, indicating a recent origin. For example, *H. e. etylus*, like *H. m. ecuadoriensis*, has a restricted forewing band shape that corresponds to the more distal element of the *notabilis* forewing band (*s*_*WntA*_=0.015). Clear, narrow, and very similar selection signals were found near *WntA* in *H. e. amalfreda* and *H. e. erato* (*s*_*WntA*_=0.006 in each), both with a broken forewing band, as well as *H. e. emma* (*s*_*WntA*_=0.003) and *H. e. lativitta* (*s*_*WntA*_=0.004), both with a narrow forewing band (Supplementary Table 8).

More broadly across the *H. erato* populations, there was a clear difference between the Amazonian dennis-ray races (i.e. *H. e. amalfreda, H. e. erato, H. e. emma, H. e. etylus* and *H. e. lativitta*), all exhibiting a similar selection pattern at *optix*, and red forewing band races (*H. e. favorinus, H. e. venus, H. e. cyrbia* and *H. e. hydara* in Panama, and *H. e. demophoon*) which showed little or no signature of selection. This is in agreement with the hypothesis that the widespread dennis-ray phenotype at *optix* has a more recent origin as compared with the red band phenotype ^48^. One notable exception to this pattern was *H. e. hydara* in French Guiana, the only red banded *H. erato* form with a strong signal at *optix* (*s*_*optix*_=0.09). There are slight variations across the range in the band phenotype, and perhaps a recent modification of the band phenotype swept in this population. The pattern in *H. melpomene* is less clear, possibly due to age of the alleles and the considerably lower effective population size in *H. melpomene*.

At the *cortex* locus, there was a consistent peak centred on *lethal (2)* just next to *domeless* and *washout* (annotated in Supplementary Fig. 11). However, surprisingly the signal is almost identical across populations with a variety of different yellow colour pattern phenotypes (*H. e. amalfreda, H. e. erato, H. e. hydara* in French Guiana, *H. e. emma, H. e. etylus, H. e. lativitta, H.e. notabilis, H. e. favorinus, H. himera*), and completely absent in North-Western populations (*H. e. cyrbia, H. e. venus, H. e. hydara* in Panama, *H. e. demophoon*) (Supplementary Fig. 8). The sweep signal therefore shows little obvious association with any particular wing pattern phenotype but may still indicate a locus involved in the colour pattern pathway. In addition, we detected very distinct signals between *H. e. favorinus* (*Cr1*) and *H. e. demophoon (Cr2)* consistent with previous studies ^31,33,83^ that found evidence for independent evolution of the yellow hindwing bar on either side of the Andes. While *H. e. favorinus* lacks any signature at *Cr2* and shows a weak signal at *Cr1*, a clear peak was found for *H. e. demophoon* at *Cr2* indicating that this allele may be more recent (Fig. 5, Supplementary Fig. 8 and 11).

## Discussion

Elucidating the evolutionary history and spread of advantageous variants in natural populations lies at the heart of evolutionary research, ever since Wallace ^84^ and Darwin ^85^ established the theory of evolution by natural selection. However, detecting and quantifying selection has been a challenge, particularly in wild populations ^3^. We have combined a large dataset of high coverage genomic data with novel theoretical analyses to identify molecular signatures of recent selection at genes known to control adaptive wing patterning traits in *Heliconius* butterflies. We demonstrate that these strongly selected loci have been subject to recent bouts of natural selection even within the last 100,000 years, with geography and phenotype standing out as strong predictors of selection (Fig. 6).

**Figure 6.**
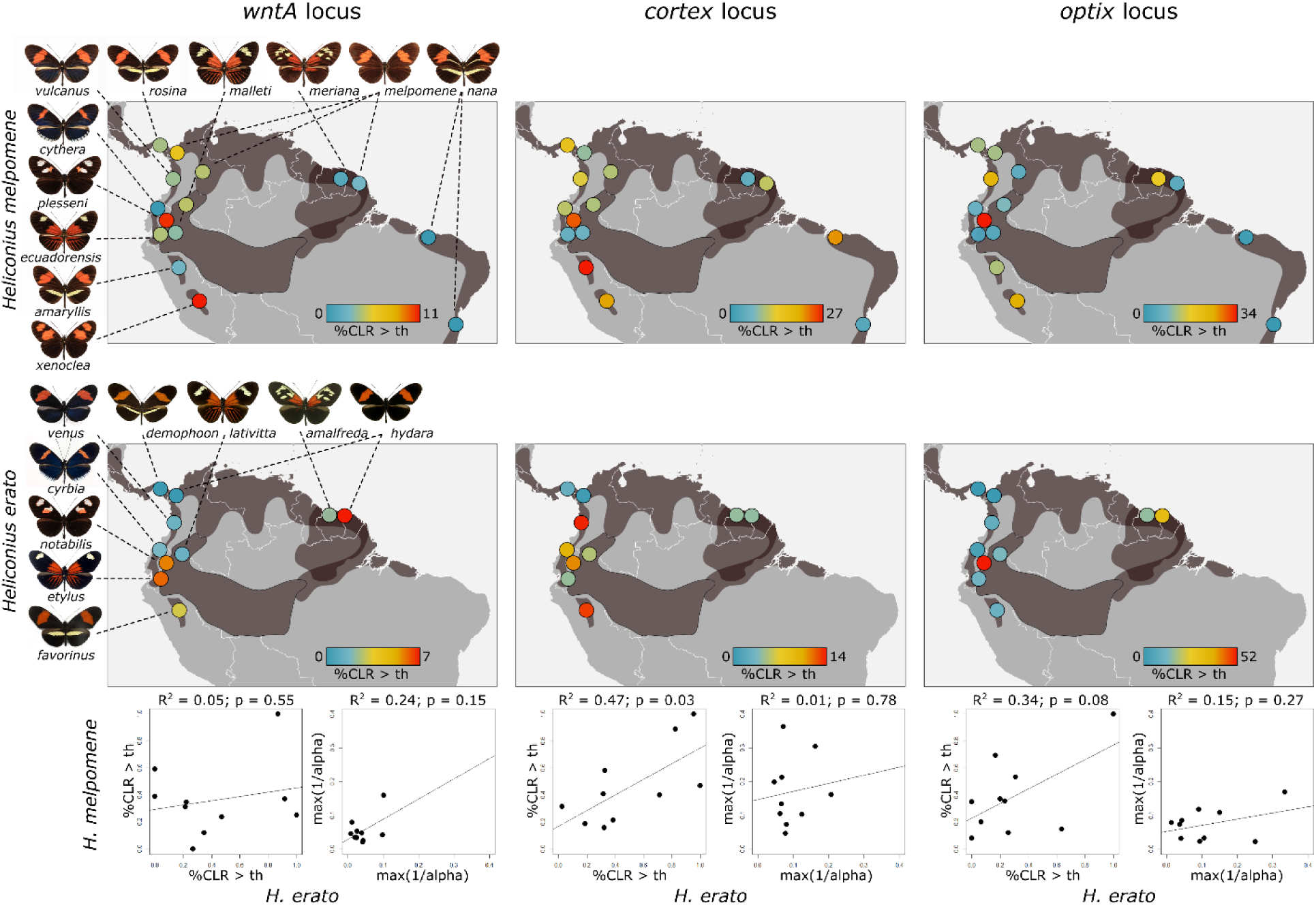
Geographic mapping of colour pattern selection in *H. melpomene* (top) and *H. erato* (middle). Dark-grey shadings indicate distributional ranges of the depicted colour patterns. Coloured circles indicate the colour pattern selection summarized as percentage of CLR values across the colour pattern region which are above the CLR threshold [%CLR>th] scaled by the maximum value for *WntA, cortex* and *optix* regions (left to right) in *H. melpomene* (top) and *H. erato* (middle). The bottom panel shows correlations for percentage CLR values above threshold [%CLR>th] and maximum intensity of selection [max(1/*α*)] between *H. melpomene* and *H. erato*.

Many studies have used naive genome scans to identify selection in natural populations, but such an approach can lead to false positives ^86^. More integrative approaches, which combine selection scans with information on phenotypic selection in the wild and genetic trait mapping, can give a more complete picture of how selection shapes specific loci and phenotypes ^10,12,14,16,87^. Such studies are increasingly common, but with few exceptions focus on a single locus, or a limited set of populations or phenotypes, often because of the high sampling and sequencing effort required. We take advantage of 150 years of *Heliconius* research, including field selection experiments, hybrid zone studies and detailed dissection of the genetics of colour pattern elements, to survey genomic signatures of selective sweeps across many populations and loci. With our study design, we reconcile large geographic sampling and high-coverage sequence data by targeting well-defined regions in the genome. This combination of ‘top-down’ and ‘bottom-up’ approaches, as defined by Linnen and Hoekstra ^1^, reveals pervasive evidence for the action of natural selection on mimicry loci across an entire adaptive radiation associated with a great diversity of phenotypes.

We have shown a pervasive pattern of strong selection acting on mimicry colour patterns, which contrasts strongly with the regions flanking the selected loci and neutral background genome regions. This supports the assertion of ‘*contrasted modes of evolution in the genome’*, first formulated by John R. G. Turner 40 years ago ^69^, who concluded that mimicry genes and neutral parts of the genome were subject to different modes of evolution. Of course, our data do not preclude the existence of other strongly selected loci not associated with mimicry in the genome. The frequency of evidence for selection is consistent with the large effective population sizes in *Heliconius* that preserve the signature of selective sweeps over a relatively long period of time. Our estimates of selection strength indicate strong selection acting on mimicry genotypes, which is in line with field and hybrid zone studies on the colour pattern phenotypes (Table 1, Supplementary Tables 3 & 8) and strong selection on colour polymorphisms in other species ^1,10,88^. *Heliconius* butterflies therefore join a small group of systems for which strong natural selection on ecologically important traits has been documented in detail at both the phenotypic and molecular level ^1,2^. Other examples include Darwin’s finches, where climate-driven changes in seed size and hardness imposed strong selection on beak size and body weight ^15,89,90^, industrial melanism in the peppered moth *Biston betularia* ^88,91^, the body armour locus *Eda* in sticklebacks ^92^ and crypsis in *Peromyscus maniculatus* deer mice controlled by the *agouti* pigment locus ^16^.

However, both strength and direction of selection can vary substantially in time and space, and a snapshot of a single population may be misleading about the action of selection in the wild ^90,92–94^. One way to account for this variation is by studying patterns of selection across geographically widespread adaptive radiations, comprising ecological replicates. This approach allows us to describe general patterns in the action of selection on a continental scale. For example, there is consistently stronger selection on the *optix* and *cortex* loci across the range of these species, consistent with the greater phenotypic effect of alleles at these loci. In addition, we also identify what seem to be more recent phenotypes showing a stronger signature of selection, such as the split band phenotype in the Andes and the dennis-only phenotype on the Guiana shield (Fig. 6).

One of the defining characteristics of the *Heliconius* radiation has been the importance of adaptive introgression and recombination of pre-existing variants in generating novelty ^32,34,38^. We used simulations to explore the expected patterns resulting from both new mutations and introgressed selective sweeps. These demonstrated a distinct signature of selection on introgressed variation, consistent with recent theory ^27^ and revealed that depending on the frequency of the acquired variant, introgressed sweeps show a range of characteristics reminiscent of classic sweeps. Consistently, we found that tests designed for detecting classic sweeps can also detect introgressed sweeps, but the signal becomes narrower, and the time window for detection decreases. In addition, the power to detect selection decreases with increasing effective migration rate between hybridising species. These conclusions may explain the scarcity of selection signatures in the *Heliconius timareta* populations that represent well documented recipients of adaptive introgression but also show strong genome-wide admixture, suggesting relatively high migration rates with *H. melpomene* ^34,38,57^. Nonetheless, we detected putative introgressed sweeps in *H. c. weymeri, H. m. cythera, H. m. vulcanus* and *H. heurippa*, for which acquisition of colour pattern phenotypes via adaptive introgression has been demonstrated ^32,74,95^. Combining tests for introgression with scans for selection provides a powerful means to study adaptive introgression more generally ^e.g. see 96,97^. Improved methods specifically designed to detect the molecular signature of an introgressed sweep and investigate its strength and timing, will help to improve the accuracy of this approach ^27,97^.

Our results imply a complex history in which multiple bouts of selection have occurred at the same loci. Although recurrent sweeps can alter or even eradicate previous signatures ^5^, there is nonetheless evidence for sweeps, both at previously characterised genomic regions and in novel locations. Previously, regulatory loci have been identified based on association studies across divergent populations ^34,32,33^, and many of these regions indeed show strong signatures of selection providing further support for their functional roles. However, consistent signatures of selection are also found at nearby loci, suggesting additional targets of selection some of which had not previously been identified using top-down approaches. Some caution is required, as the signatures of selective sweeps are notoriously stochastic and can be misleading in their precise localisation due to linkage. Nonetheless, there are consistent patterns across multiple populations suggesting additional targets of selection that may represent regulatory elements affecting already characterised genes ^32,34^, similar to multiple mutations under selection at the *Agouti* gene in deer mice (*Peromyscus maniculatus*) ^10^. In addition, however, some of these signals may represent selection at linked genes, and the architecture colour pattern in *Heliconius* may be comparable to the situation in *Antirrhinum* snapdragons in which loci encoding flower pattern differences, i.e. *ROSEA* and *ELUTA*, are in tight linkage.^12^. Further functional studies will be required to unravel the roles of these loci, but theory suggests that physical linkage between genes contributing to the same adaptive trait can be favoured ^98,12^. Intriguingly, *Heliconius* butterflies show both unlinked colour pattern loci, as well as tightly linked CREs and genes within loci, putatively preserving locally adaptive allelic combinations. It is conceivable that this architecture provides a high degree of flexibility that has facilitated the radiation of colour patterns in *Heliconius*.

Müllerian mimics can exert mutual selection pressures, offering the rare opportunity to study replicated selection in a co-evolutionary context. The diversity of mimicry alleles between *H. melpomene* and *H. erato* evolved independently ^48,49^, but several co-mimics between the two radiations show signatures of selection in homologous colour pattern regions, demonstrating repeated action of natural selection between co-mimics over recent time. Our findings also contribute to long-standing arguments on the origin and spread of the colour patterns ^48,75–80^. Signatures of selection at *optix*, particularly in *H. erato*, are consistent with the hypothesis that the red forewing band is ancestral and dennis-ray is a younger innovation that spread through the Amazon. However, in contrast to this ‘recent Amazon’ hypothesis, we find the strongest signatures of selection in some of the unique and geographically restricted phenotypes found in Andean populations suggesting novel colour patterns have experienced strong recent selection in both species, consistent with co-divergence and ongoing co-evolution (Fig. 6). The most striking example are *H. e. notabilis* and *H. m. plesseni*, which show imperfect mimicry (see Fig. 5) and are possibly still evolving towards an adaptive optimum. In summary, our results provide evidence for co-divergence and the potential for co-evolution in the sense of mutual evolutionary convergence ^79^ but do not rule out advergence in other cases.

To conclude, understanding the adaptive process that creates biodiversity requires knowledge of the phenotypes under selection, of their underlying genetic basis, and estimates of phenotypic and genotypic strength and timing of selection ^1^. While decades of *Heliconius* research have resulted in a detailed understanding of most of these levels, our study fills a gap by providing estimates of the distribution and strength of genotypic selection across two radiations and dozens of populations. However, our results not only highlight the complexity of mimicry selection across the *Heliconius* radiation but also reveal a surprisingly dynamic turn-over in colour pattern evolution, in particular in geographically peripheral patterns (Fig. 6). This is in stark contrast to the predicted evolutionary inertia of mimicry patterns due to strong stabilizing selection pressure exerted by mimicry selection ^99^. We provide evidence that colour patterns are actively evolving under both classic and introgressed sweeps. Many of the detected sweep signatures are considerably younger than estimates of the age of colour pattern alleles based on phylogenetic patterns ^32,34^ suggesting ongoing improvement, innovation and local switching between combinations of pattern elements. This is also consistent with observations of phenotypically distinct colour patterns restricted to the only 5,000 year-old islands Ilha de Marajó in the South of Brazil and a few documented cases of rapid, local colour pattern turn-over ^100^. Therefore, our study offers a new perspective to the long-standing discussion of the paradox: ‘How and why do new colour patterns arise’. More generally, we here demonstrate that by considering selection across populations and species of an entire radiation, comparative information can capture spatial and temporal variability of genotypic selection and help to gain a more comprehensive understanding of the dynamics of adaptation in the wild.

## Methods

### Sampling and DNA extraction

Our sampling covers most of the distribution and colour pattern variation of the *Heliconius* radiation in South and Central America. Specimens were sampled or provided by collaborators with the respective sampling permissions and stored in salt saturated DMSO or ethanol at −20°C until further processing. For DNA extractions, thorax muscle tissue was dissected, disrupted, digested, and DNA was extracted using a TissueLyser II bead mill together with the DNeasy Blood and Tissue Kit (Qiagen, Hilden, Germany) following supplier recommendations.

### Targeted capture and sequencing

For hybridization-based target enrichment a NimbleGen SeqCap EZ Library SR capture probes library was designed and synthesized by the provider (Roche NimbleGen Inc, United States). The templates for designing probes for four colour pattern regions (∼ 3.2 Mb) and four genomic background regions (∼ 2 Mb) were assembled and curated using the *H. melpomene* genome assembly Hmel1 ^38^, available BAC walks ^52,101^, fosmid data ^50^, and alignments from Wallbank *et al.* ^34^. The neutral background regions were chosen to represent the average genome. We therefore excluded regions with extended stretches of extreme values for diversity and/or divergence and we only considered regions located on a single, well-assembled scaffold.

Sample DNA was sheared with an ultrasonicator (Covaris Inc, Massachusetts, United States) and adapter-ligated libraries with insert sizes of 200-250 bp were generated using the Custom NEXTflex-96 Pre-Capture Combo Kit (Bioo Scientific Corporation, United States). For sequence capture, 24 libraries each were pooled into a capture library, hybridized with blocking oligos and the biotinylated capture library probes, and subsequently captured with streptavidin-coated magnetic capture beads using the NimbleGen SeqCap EZ Kits (Roche NimbleGen Inc, Wisconsin, United States). After capture and clean-up, three capture library pools were combined, each. For the resulting sequencing pools of 72 samples, Illumina 100 or 150 bp paired-end short read data were generated on Illumina’s HiSeq 2000 (BGI, China) and HiSeq 4000 (Novogene Co. Ltd, China), respectively (Supplementary Table 1).

### Whole genome data

Whole genome resequencing data available for the *melpomene*-clade from previously published work were also included ^31,32,36,38,39,54,56–58^. For a few additional samples, 100-150 bp paired-end whole genome resequencing data were generated on an Illumina X Ten platform (Novogene Co. Ltd, China) (Supplementary Table 1). For the *erato*-clade already published whole genome-resequencing data were used ^33^ (Supplementary Table 7).

### Genotyping

For *melpomene-*clade data, sequenced reads were aligned to the *H. melpomene* v2 reference genome (Hmel2, Davey et al. 2016), using BWA-mem v0.7 ^102^. PCR duplicated reads were removed using Picard v2.2.4 (http://picard.sourceforge.net) and reads were sorted using SAMtools v1.3.1 ^103^. Genotypes for variant and invariant sites were called using the Genome Analysis Tool Kit’s (GATK) Haplotypecaller v3.5 ^104^. Individual genomic VCF records (gVCF) were jointly genotyped per population using GATK’s genotypeGVCFs v3.5 ^104^. Genotype calls were only considered in downstream analyses if they had a minimum depth (DP) ≥ 10, and for variant calls, a minimum genotype quality (GQ) ≥ 30, and indels were removed. Filtering was done with bcftools v.1.4 ^103^, and for downstream calculations of summary statistics and creating SweepFinder2 input, vcf files were parsed into tab delimited genotype files (scripts available at https://github.com/simonhmartin). For the *erato-*clade, read data were mapped to the *H. erato demophoon* v1 genome reference ^33^ and further processed as described above.

### Phasing

SHAPEIT2 ^105^ was used to phase haplotypes using both population information and paired read information. First, monomorphic and biallelic sites were filtered with GQ ≥ 30 and DP ≥ 10 and sites with less than 20% of sample genotypes were removed.

Next, phase informative reads (PIRs) with a minimum base-quality and read quality of 20 were extracted from individual BAM files using the extractPIRs tool. These BAM files were obtained from BWA-mem ^102^ mappings to the *H. melpomene* v2 genome, with duplicates removed. Finally, SHAPEIT2 was run with PIR information and default parameters on each scaffold using samples from single populations, which resulted in a haplotype file that was transformed into VCF format. Sites with no genotype information were imputed.

### Phylogenetic reconstruction

FastTree2 ^106^ was run using default parameters to infer approximate maximum likelihood phylogenies. Separate phylogenies for a concatenated SNP dataset comprising neutral background regions only and for the full dataset including the colour pattern regions for a phylogeny to account for the effect of including regions putatively under strong selection were produced.

### Population historical demography

Changes in the historical population size were inferred from individual consensus genome sequences using Pairwise Sequentially Markovian Coalescent (PSMC’) analyses as implemented in MSMC ^107^. This method fits a model of changing population size by estimating the distribution of times to the most recent common ancestor along diploid genomes. When used on single diploid genomes, this method is similar to pairwise sequentially Markovian coalescent (PSMC) analyses ^108^. Genotypes were inferred from BWA v0.7 ^102^ mapped reads separately from previous genotyping analysis using SAMtools v0.1.19 ^103^ according to authors’ suggestions. This involved a minimum mapping (-q) and base (-Q) quality of 20 and adjustment of mapping quality (-C) 50. A mask file was generated for regions of the genome with a minimum coverage depth of 30 × and was provided together with heterozygosity calls to the MSMC tool. MSMC was run on heterozygosity calls from all contiguous scaffolds longer than 500 kb, excluding scaffolds on the Z chromosome. We scaled the PSMC’ estimates using a generation time of 0.25 years and a mutation rate of 2 × 10^−9^ estimated for *H. melpomene* ^63,67^.

### SLiM Simulations

Simulations were conducted to compare the genomic signatures of classical selective sweeps and sweeps that occur via adaptive introgression using SLiM (version 2) forward in time population simulation software ^109,110^. Two populations of *N* = 1000 were simulated with a neutral mutation rate *µ* of 6 × 10^−7^ such that the expected level of neutral diversity in the population was 0.0024, which is within an order of magnitude of that observed in our *Heliconius* populations. Each individual in our simulated populations was represented by a single diploid recombining chromosome (recombination rate was also scaled such that *NR* is within the values of those observed in *Heliconius*, 4 × 10^−7^, or 40 cM/Mb), of length 750,000 bp.

Our simulations were first allowed to equilibrate for a burn-in phase of 10*N* generations, after which we introduced a single strongly advantageous mutation of *s* = 0.5 in the centre of the chromosome, in order to simulate a ‘classical’ hard selective sweep in the population (which we will refer to as p1). Only those simulations in which the mutation went to fixation were kept: if the beneficial mutation was lost during the course of a simulation, the simulation was reset to a point just after the burn-in phase and the mutation was reintroduced. The simulations were then allowed to run for a further 5*N* generations. During this time, p1 does not experience any migration or population size change. In order to simulate an introgressed sweep, we simulated an additional neutrally-evolving population, p2, which exchanges migrants with population p1 at a constant rate of 0.0001 migrants per generation, which allowed the beneficial mutation fixed in p1 to introgress into p2. The simulations were then allowed to run for a further 10*N* generations with a constant migration rate. For each set of parameters, we ran our simulations 100 times. For both populations, a complete sample of the segregating neutral mutations was taken every 100 generations after the burn-in phase and prior to the introduction of the beneficial mutation, and every 50 generations after the introduction of the beneficial mutation. We also tracked the change in frequency over time of the beneficial mutation during the simulations. From these results we calculated two summary statistics, Tajima’s *D* and *π*, in windows of 10,000 bp across our simulated chromosomes for a range of time-points. Time-points are as follows, in 4*N* generations post sweep: 0.01, 0.1, 0.2, 0.3, 0.4, 0.5, 0.6, 0.7, 0.8, 1, and two background rates: one post burn-in, during which populations are not experiencing any migration, and one post-sweep, during which the populations are exchanging migrants. Values were then averaged across simulations. Additionally, to model the effect of changing effective migration rates on the introgression sweep signal we ran simulations with different levels of migration, using the following 4 values of *M*: 200, 2, 0.2 and 0.02, with recombination rate = 4cM/Mb and *s* = 0.1. The simulations were otherwise set up as before, with 30 simulation runs generated for each set of parameters.

We also used these results to generate SweepFinder2 ^62^ input files, after first subsampling the number of mutations down, such that our simulated SweepFinder2 files for each population represent a sample of 500 simulated individuals. This step is necessary because SweepFinder has an upper limit on the number of sequences that can be included per sample ^111^. We then ran SweepFinder2 using mode –lg 100 for each simulation for each of the time-points, using one of two pre-computed site frequency spectra as appropriate: one calculated across multiple neutral simulations without migration, and one calculated across multiple neutral simulations with migration (these neutral simulations correspond to the two background rates described above). Further details of SweepFinder2 and its various run modes are included in the ‘SweepFinder2’ section.

### Phylogenetic weighting

A phylogenetic weighting approach was used to evaluate the support for alternative phylogenetic hypotheses across colour pattern loci using *Twisst* ^112^. Given a tree and a set of pre-defined groups, in this case *Heliconius* populations sharing specific colour pattern elements, *Twisst* determines a weighting for each possible topology describing the relationship of the groups. The weightings thus represent to what extent loci cluster according to phenotype, rather than geographic relatedness of populations. Topology weightings are determined by sampling a single member of each group and identifying the topology matched by the resulting subtree. This process is iterated over a large number of subtrees and weightings are calculated as the frequency of occurrence of each topology. Weightings were estimated from 1,000 sampling iterations over trees produced by RAxML v8.0.2681 ^113^ for 50 SNP windows with a stepping size of 20 SNPs. For phylogenetic weighting along the *WntA* interval, weightings of topologies that grouped populations with the split forewing band phenotype or, alternatively, the hourglass shape were assessed (Supplementary Fig. 7). For the region containing the *aristaless* genes, we focused on topologies that clustered populations with white or yellow colour phenotypes (Supplementary Fig. 12). For the *cortex* region we focused on topologies grouping populations showing the ventral and dorsal yellow hindwing bar, respectively (Supplementary Fig. 13). Finally, for the *optix* interval we assessed topologies grouping populations according to the absence or presence of the red dennis patch, the red hindwing rays or the red forewing band and repeated the analysis for different geographic settings (Supplementary Fig. 14). To obtain weightings for hypothesized phylogenetic groupings of specific colour pattern forms, we summed the counts of all topologies that were consistent with the hypothesized grouping.

### Inference of selection and summary statistics in sliding windows

Summary statistics informative on diversity and selection patterns were calculated. From the unphased data, nucleotide diversity, Tajima’s *D*, and number of sites genotyped for each population were calculated in 1 kb non-overlapping sliding windows with at least 100 sites genotyped for at least 75% of all individuals within that population using custom python scripts and the EggLib library v3^114^. Scans for selection using signals of extended haploptype homozygosity and calculation of the pooled integrated haplotype homozygosity score (iHH12) ^11,115^ were performed using the program selscan1.2 ^116^ and our phased dataset.

### SweepFinder2

To detect local distortions of the site-frequency spectrum that are indicative of selective sweeps, SweepFinder2, an extension of Nielsen et al.’s ^60^ SweepFinder program, with increased sensitivity and robustness ^61,62^ was used. The SweepFinder framework builds on a composite likelihood ratio test using the site frequency spectrum to compare the likelihood for a model with a selective sweep *versus* the likelihood for a model without a sweep. Huber *et al.* ^61^ showed that including substitutions, i.e. fixed differences relative to an outgroup, increases power while maintaining robustness to variation in mutation rate. SweepFinder2 also permits the use of recombination maps. The use of polarised sites increases power and we therefore polarized sites when possible.

We filtered our dataset for biallelic sites only and initially tested different input datasets and parameter settings and created two types of datasets for this purpose; one using polymorphic sites only with both polarised and unpolarised sites, and one with polymorphic sites and substitutions that contained only polarised sites. As an outgroup, *Heliconius numata* was used for the *melpomene*-clade and *H. hermathena* for the *erato*-clade. We used biallelic sites only that were present in ≥75% of the focal populations and polarized sites by randomly drawing an outgroup allele from sites with a minimum number of outgroup samples with genotype data of either one (-OM1) or three (-OM3) of four for the *melpomene*-clade and one (-OM1) or two (-OM2) of three for the *erato*-clade.

SweepFinder2 was then run in two modes for each dataset; with flag -s, calculating the likelihoods from the site-frequency spectrum of the respective region and with flag -l, using a site-frequency spectrum pre-calculated either from the background regions only or from background regions and colour pattern regions combined. For the *melpomene*-clade, recombination rate information from a fine scale recombination map was included (flag -r) ^117^. To create a recombination file, recombination map coordinates were transferred to Hmel2 coordinates and between sites recombination rates were calculated.

SweepFinder2 test runs for different grid spaces (flag –g; tested values: -g1, -g5, -g50, -g100, -g1000) were performed to find a setting allowing for reasonable runtimes without loss of accuracy and based on these test CLR and α were calculated for every 50th site (-g50) across all populations and regions.

Generally, the results were largely consistent among the different runs and datasets. As expected power to detect sweeps was higher when including substitutions ^61^ and the minimum number of outgroup samples had only marginal effects. We therefore focussed on the results for datasets with outgroup minimum 1 (-OM1) and background SFS calculated from background regions and background regions and colour pattern regions combined, respectively. Including the colour pattern regions inflates the estimated background SFS with regions affected by selective sweeps which results in slightly lower CLR and higher α estimates. Since selective sweeps across the genome have been found to be rare in *H. melpomene* ^54^, these estimates represent a lower bound and the estimates derived with background SFS from the background regions only are most likely a better approximation. Only CLR peaks exceeding a threshold defined as the 99.9th percentile of the distribution of CLR values across all background regions were considered as evidence for selection.

To obtain estimates for strength of selection (s) we used the formula from Nielsen et al. ^60^ and calculated *s* as *s* = *r* × ln(2*N*_*e*_) /*α* with region- and population-specific estimates of *N*_*e*_ estimated from the data using the mutation rate given in Keightley et al. ^63^ and per chromosome recombination rate estimates from Davey et al. ^117^ and Van Belleghem et al. ^33^. All custom scripts used for filtering and creating SweepFinder2 input files are available from the authors upon request.

## Data availability

GenBank accession numbers for all whole-genome and capture samples are available in Supplementary Table 1.

## Acknowledgements

We thank John Davey and Richard Wallbank for assistance with the capture design and Joe Hanly and Erica Westerman for providing *aristaless* CRE coordinates. We are also grateful to Chris Kozak, Patricio Salazar, Gislene L. Gonçalves, and Jake Morris for their help with samples. We thank Ian Warren for help with database management. Furthermore, we thank Christian Huber, Derek Setter, and Joachim Hermisson for valuable discussions and their help with selection analyses. We acknowledge the lepbase.org database ^118^ for providing valuable resources for analysis and the Welcome Trust - Medical Research Council Stem Cell Institute for technical support. This work was performed using resources provided by the Cambridge Service for Data Driven Discovery (CSD3) operated by the University of Cambridge Research Computing Service (http://www.csd3.cam.ac.uk/), provided by Dell EMC and Intel using Tier-2 funding from the Engineering and Physical Sciences Research Council (capital grant EP/P020259/1), and DiRAC funding from the Science and Technology Facilities Council (www.dirac.ac.uk). Colombian specimens were collected under the permit 530 granted to Universidad del Rosario by the Autoridad Nacional de Liencias Ambientales (ANLA-Colombia).

## Funding

The project was funded by an ERC Advanced Grant to CJ (FP7-IDEAS-ERC 339873). M.M. was also supported by an Early Postdoc Mobility fellowship (P2EZP3_148773) of the Swiss National Science Foundation (SNF) and an Erwin Schrödinger fellowship (J 3774) of the Austrian Science Fund (FWF). S.M.V.B. was supported by an Isaac Newton Trust Grant of Trinity College Cambridge. C.S. was supported by the Universidad del Rosario ‘Big Grant’ IV-FGD-001 and COLCIENCIAS (Grant FP44842-5-2017). N.J.N. was supported by the UK Natural Environment Research Council (NERC) through an Independent Research Fellowship (NE/K008498/1). M.J. was funded by Agence Nationale de la Recherche through grants ANR-12-JSV7-0005 and ANR-18-CE02-0019-02. G.R.P.M. was supported by a CNPq fellowship (Grant 309676/2011-8).

## Author contributions

M.M., S.M.V.B., J.E.J. and C.D.J. designed the study and wrote the paper. M.M., S.M.V.B. and J.E.J. analysed the data. S.H.M contributed scripts and data for population genomics analyses. C.S., G.R.P.M., C.M., M. J., and N.J.N contributed samples for sequencing. S.L.B. extracted DNA and helped with library preparation and targeted capture and database management. All authors read and commented on the manuscript.

## Competing interests

The authors declare no competing interests.

## Materials & Correspondence

Correspondence and material requests should be addressed to Markus Moest

