## Supplementary Material for "Classic and introgressed selective sweeps shape mimicry loci across a butterfly adaptive radiation"

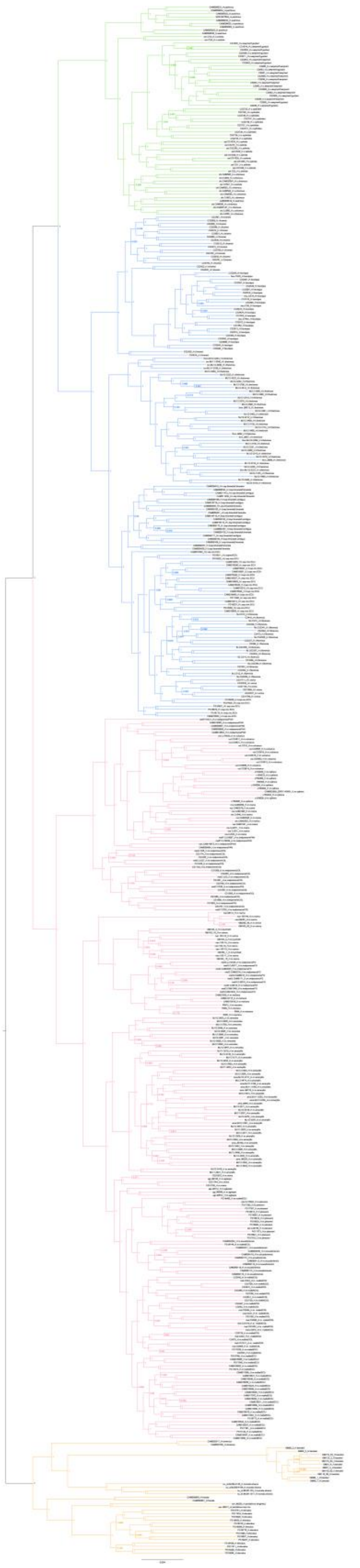

**Supplementary Figure 1. Phylogenetic reconstruction of the *H. melpomene*-clade.** Phylogenetic reconstruction for *H. melpomene*-clade samples used in this study including all sequenced region, i.e. colour pattern regions and neutral background regions. *Heliconius cydno* (green) and *H. timareta* (blue) cluster together and form a sister clade to *H. melpomene* (red). The ‘silvaniforms’ outgroup is shown in orange.

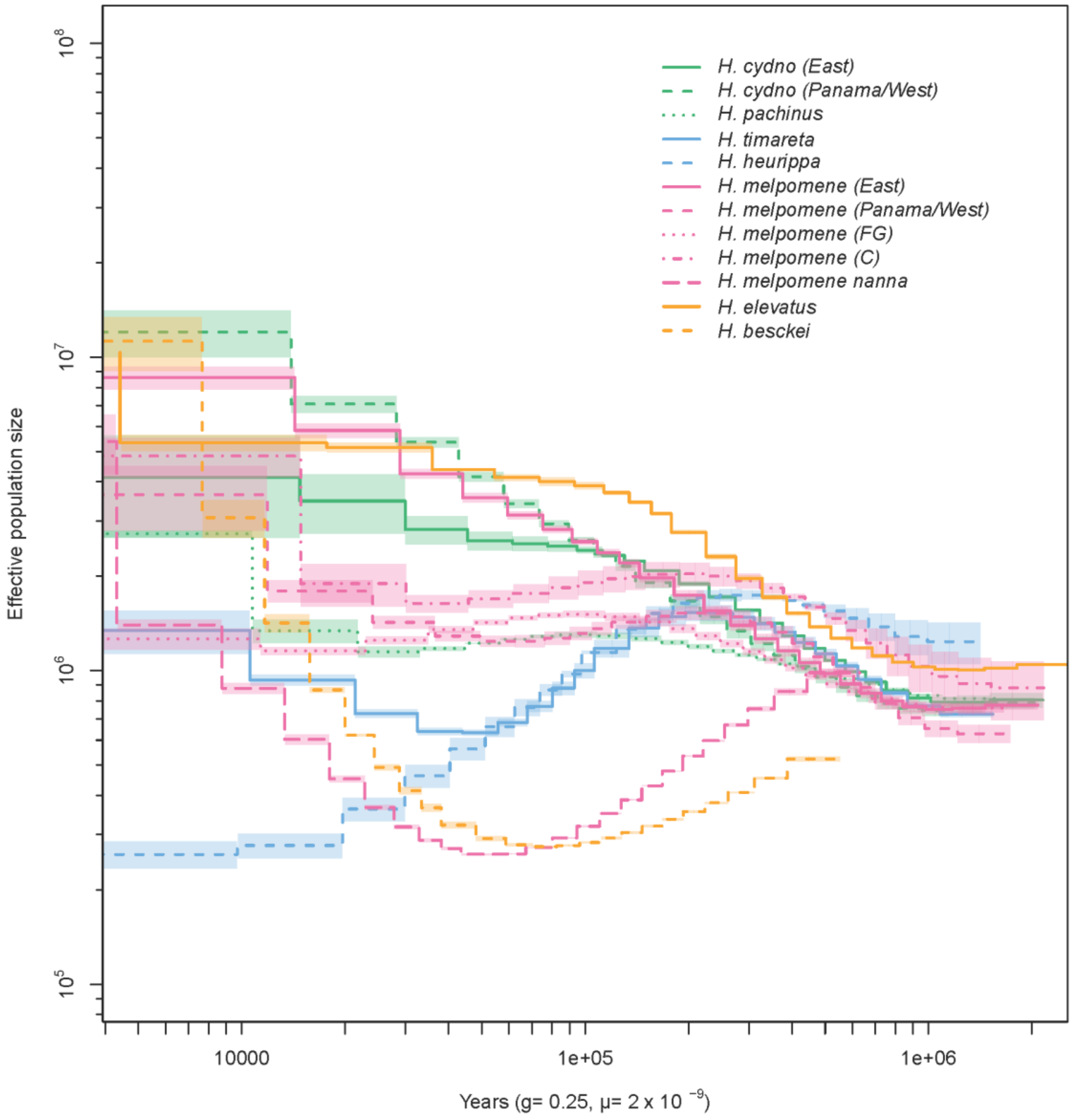

**Supplementary Figure 2. Demographic history of *Heliconius melpomene*-clade populations.** Demographic histories for populations in the *Heliconius melpomene*-clade for which whole genome data were available reconstructed with PSMC'[1]. Additional demographic histories for *Heliconius* species considered in this study are already published[2].

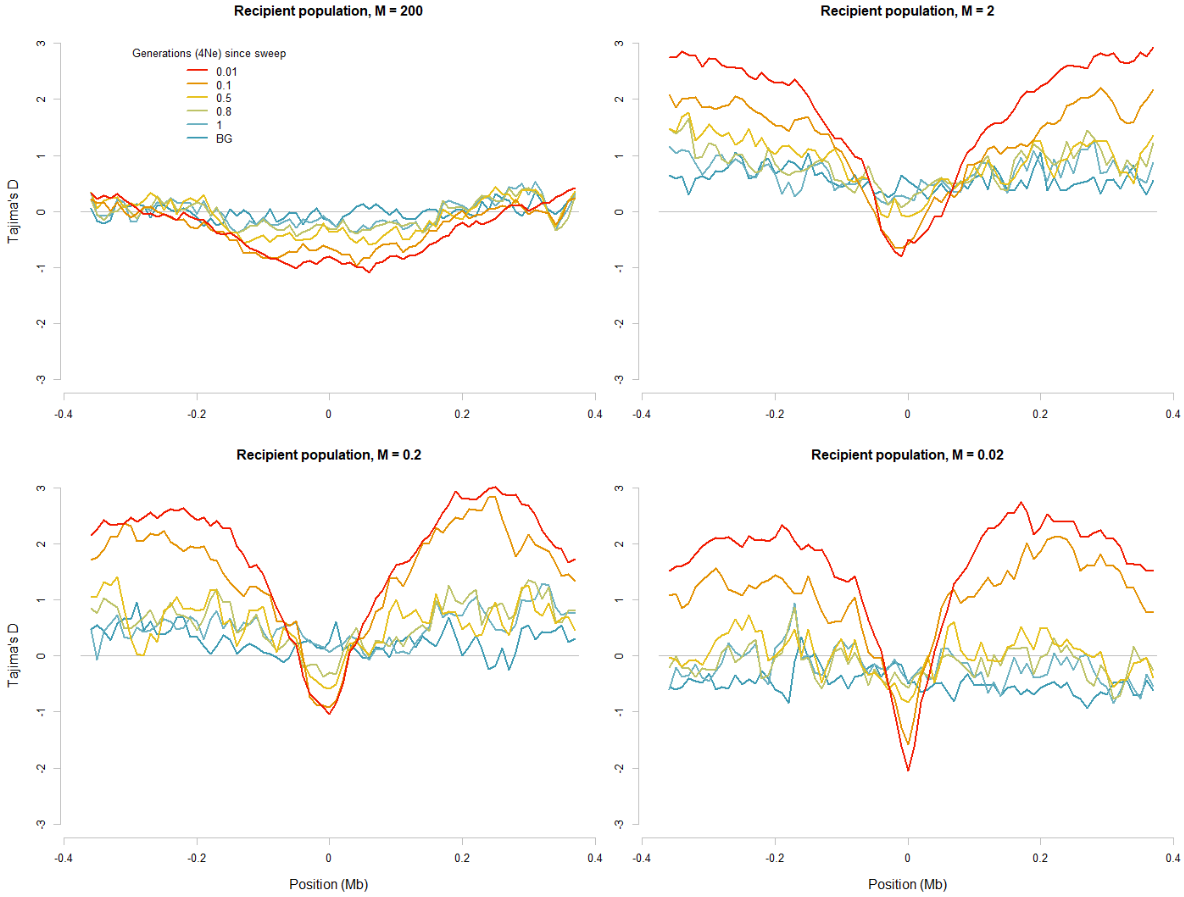

**Supplementary Figure 3. Effect of effective migration rate on introgressed sweep signatures.** Site frequency spectrum (SFS) signatures of simulated introgressed sweeps across a chromosome for different time points summarised as Tajima's *D* statistics. The sweep occurs in the centre of the simulated chromosome. Different colours indicate patterns at different time points since sweep (0.01, 0.1, 0.5, 0.8, and 1 scaled generations, *i.e.* 4*N* generations). Simulated data for four different effective migration rates are shown (*M* = 200, 2, 0.2, and 0.002).

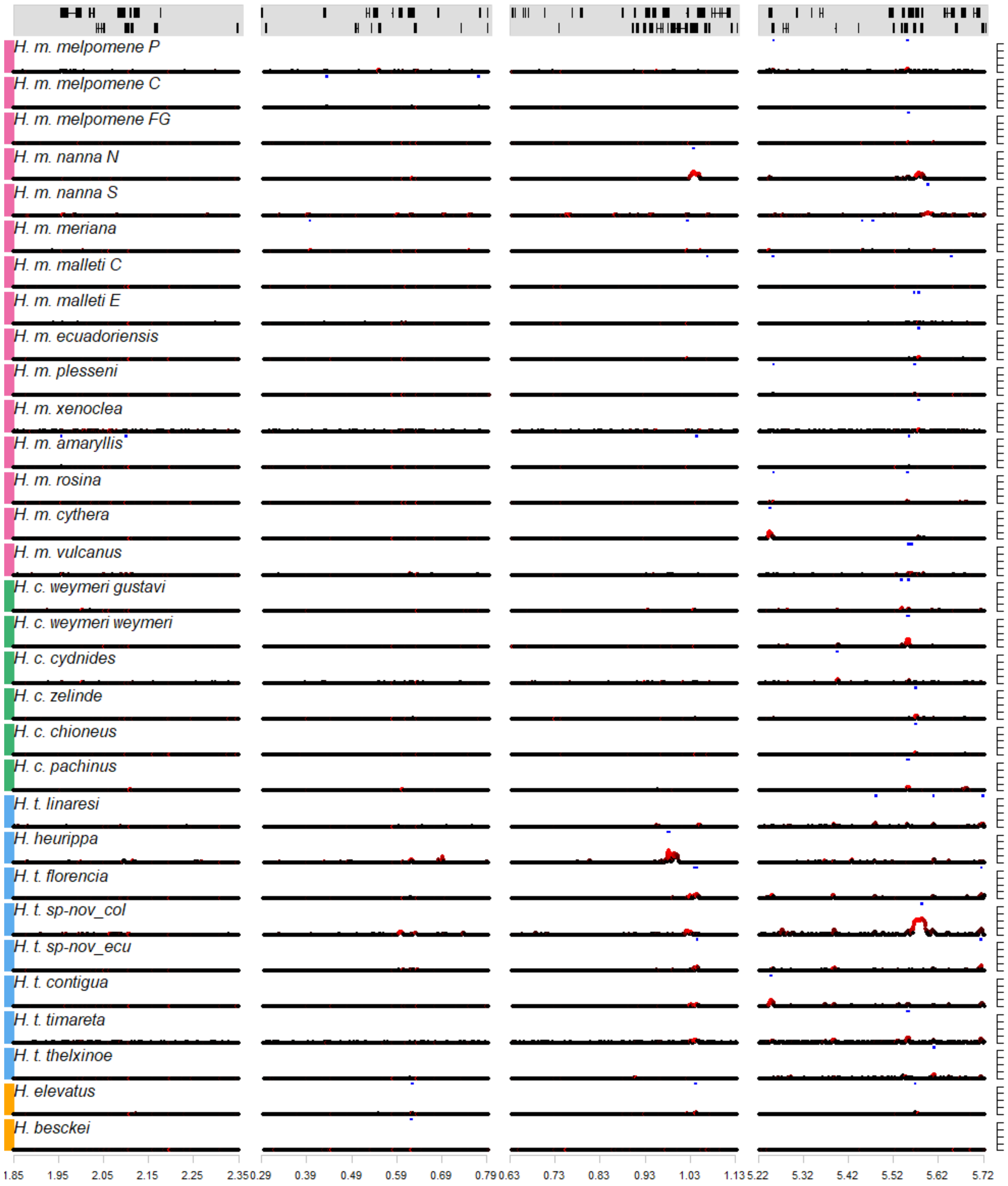

**Supplementary Figure 4. Signatures of selection across neutral background regions in the *H. melpomene*-clade.** Genes are annotated in the top gene annotation panel. On the y-axis Sweepfinder2's[3,4] composite likelihood ratio statistics (CLR) is shown (peaks are capped at CLR = 1,000). The colour gradient indicates estimated intensity of selection (black…high $\alpha$ values, weak selection; red…low $\alpha$ values, strong selection). Blue horizontal bars indicate regions with CLR values above threshold. *H. m. vicina, H. m. aglaope*, *H. m. burchelli* and *H. c. cordula* are not shown due to low sample size.

**Supplementary Figure 5. Summary and selection statistics across colour pattern regions for all populations analysed in the *Heliconius melpomene*-clade.** For each population genotyping coverage (calculated as proportion of retained genotypes after quality filtering in 500 bp windows), nucleotide diversity, Tajima's *D*, pooled integrated haplotype homozygosity score, and SweepFinder2's[3,4] composite likelihood ratio statistics across each colour pattern region are shown (top to bottom). File names contain population and colour pattern region identifiers (Hmel201011…*aristaless*, Hmel210004…*WntA*, Hmel215006…*cortex*, Hmel218003…*optix*). The 120 single pictures have been uploaded to github:

https://github.com/markusmoest/SelectionHeliconius/tree/master/H_melpomene

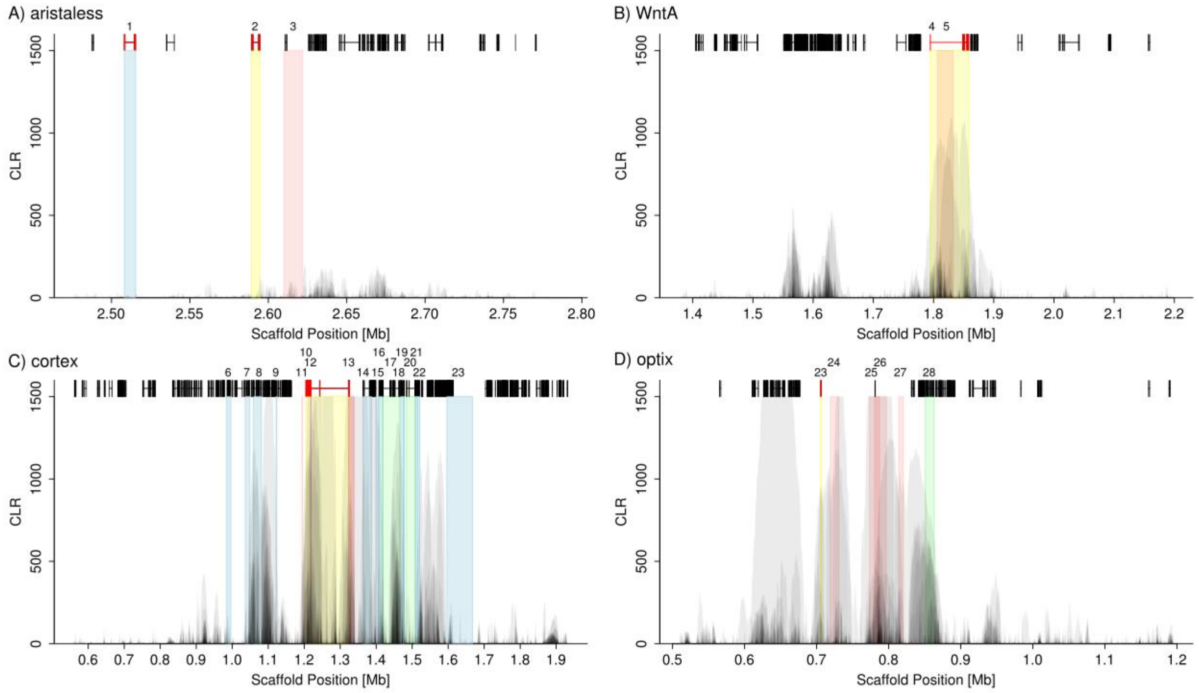

**Supplementary Figure 6. Superposition of SweepFinder2's[3,4] composite likelihood ratio peaks of all *H. melpomene*-clade populations for each of the four colour pattern regions.** Colour pattern genes (yellow), known CREs (red), and additional genes with evidence for a putative role in colour patterning (blue and green for genes discussed in the main text) are highlighted and assigned a number in the top row. The scale on the x-axes differs and the y-axis is capped at CLR = 1,500. **(A)** *aristaless1* (yellow, 2), *aristaless1* CRE (red, 3)[5], *aristaless2* (blue, 1); **(B)** *wntA* (yellow, 4), CRE associated with split forewing band indentified in this study (red, 5); **(C)** *cortex* (yellow, 10), CREs for dorsal (11) and ventral (12) hindwing topology[6], a region containing SNPs with strongest association with forewing band[7] (13) (red), additional genes with evidence for wing patterning control[7] (blue: 7,8,9,14,15,16,18,19,21,22,23; green: 17 (*LMTK1* /HM00033), 20 (*washout/WAS homologue 1*/HM00036); also see Supplementary Table 6); **(D)** *optix* (yellow, 23), CREs for 'band1'(24), 'band2'(26), 'rays'(25) and 'dennis'(27) (red)[8,9], *kinesin* (green, 28)[10,11].

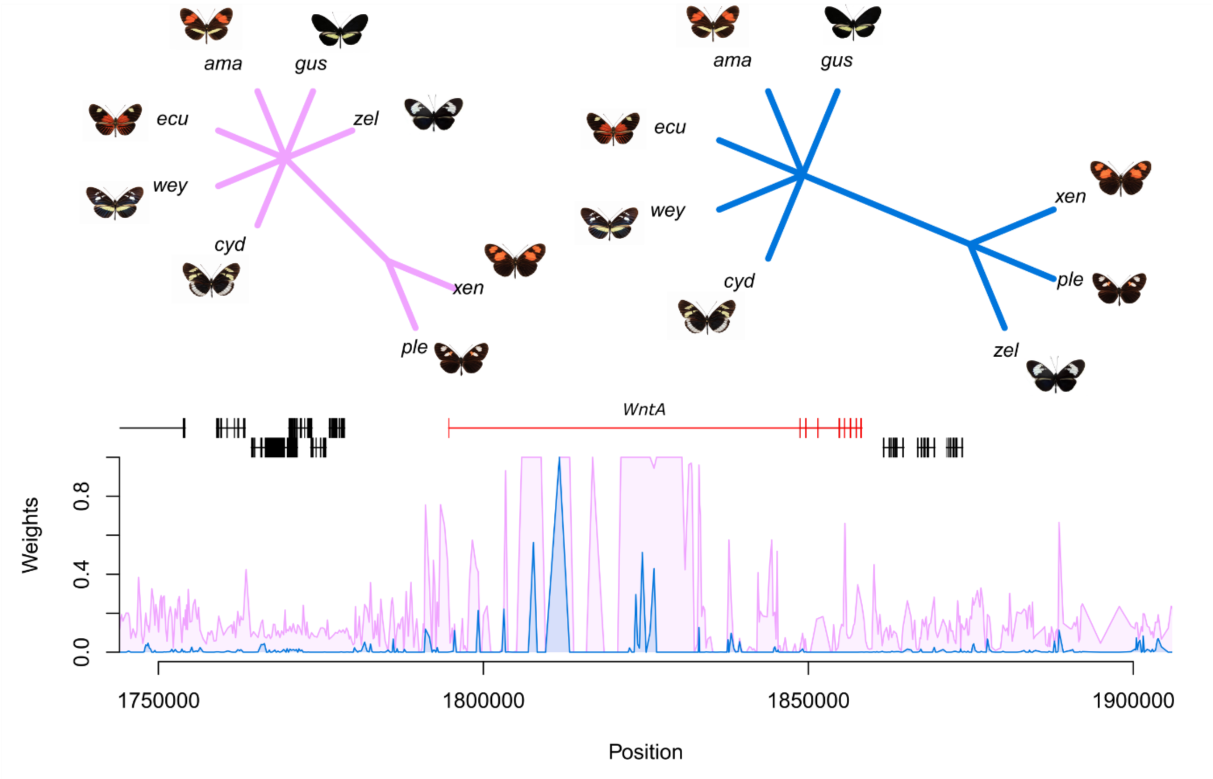

**Supplementary Figure 7. Tree weighting (Twisst[12]) analysis of the *WntA* gene region.** Topology weightings for topologies clustering the split-forewing band phenotype (magenta) and the hourglass shape phenotype (blue) are shown. (ama…*H. m. amaryllis*, ecu…*H. m. ecuadoriensis*, ple…*H. m. plesseni*, xen…*H. m. xenoclea*, cyd…*H. cydnides*, wey…*H. c. weymeri f. weymeri*, gus…*H. c. weymeri f. gustavi*, zel… *H. c. zelinde*)

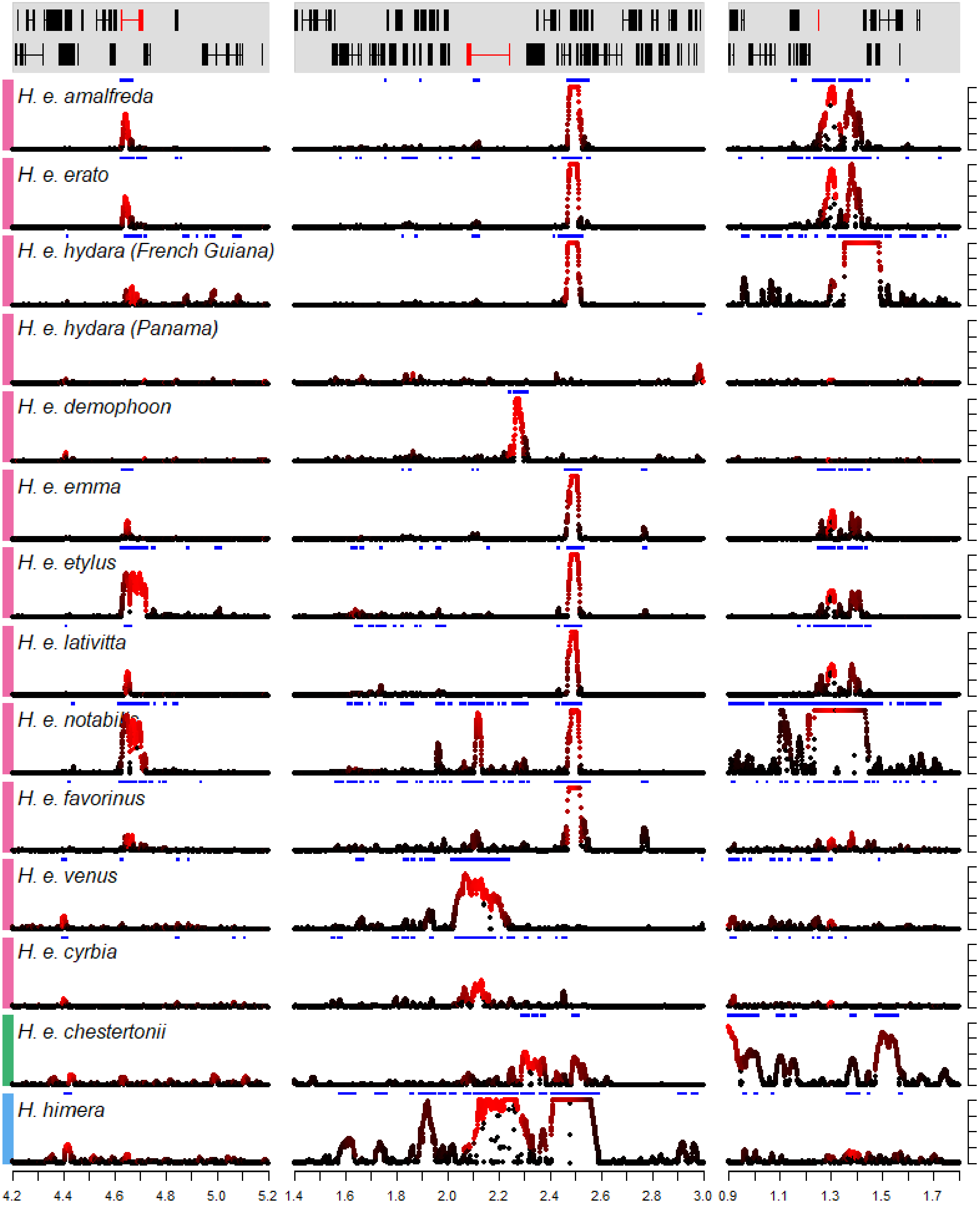

**Supplementary Figure 8. Signature of selection across colour pattern regions in the *H. erato*-clade.** The regions containing *WntA*, *cortex*, and *optix* (left to right) are depicted. Colour pattern genes are annotated in red in the gene annotation panel. On the y-axis Sweepfinder2's[3,4] composite likelihood ratio statistics (CLR) is shown (peaks are capped at CLR = 1,000). The colour gradient indicates the estimated intensity of selection (black…high $\alpha$ values, weak selection; red…low $\alpha$ values, strong selection). Blue horizontal bars indicate regions above the CLR threshold value.

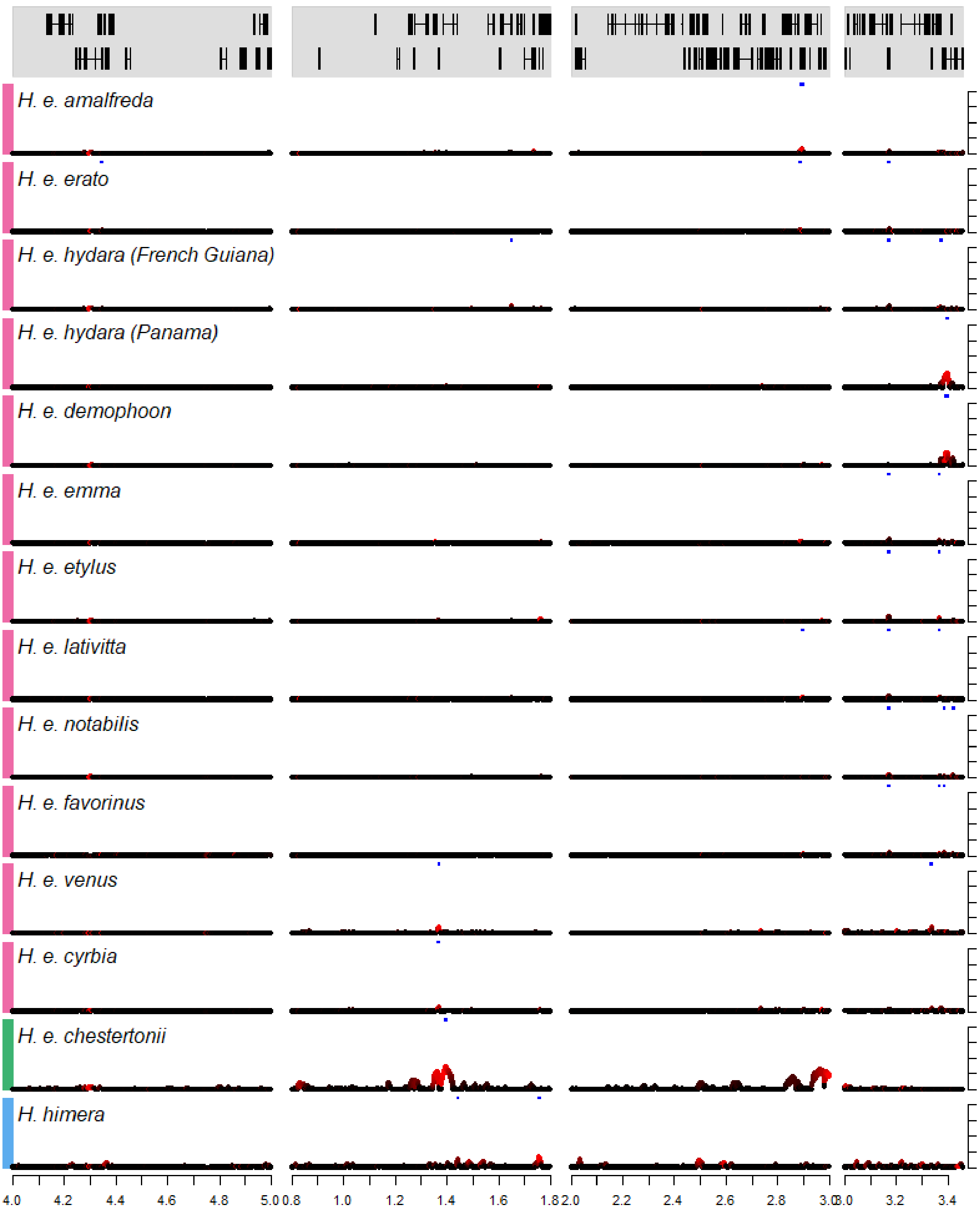

**Supplementary Figure 9. Signature of selection across neutral background regions in the *H. erato*-clade.** Genes are annotated in red in the gene annotation panel. On the y-axis Sweepfinder2's[3,4] composite likelihood ratio statistics (CLR) is shown (peaks are capped at 1,000). The colour gradient indicates the estimated intensity of selection (black…high $\alpha$ values, weak selection; red…low $\alpha$ values, strong selection). Blue horizontal bars indicate regions above the CLR threshold value.

**Supplementary Figure 10. Summary and selection statistics across colour pattern regions for all populations analysed in the *Heliconius erato*-clade.** For each population genotyping coverage (calculated as proportion of retained genotypes after quality filtering in 500 bp windows), nucleotide diversity, Tajima's *D*, pooled integrated haplotype homozygosity score, and SweepFinder2's[3,4] composite likelihood ratio statistics across each colour pattern region are shown (top to bottom). File names contain population and colour pattern region identifiers (Herato1001…*WntA*, Herato1505…*cortex*, Herato1801…*optix*). The 18 single pictures have been uploaded to github:

https://github.com/markusmoest/SelectionHeliconius/tree/master/H_erato

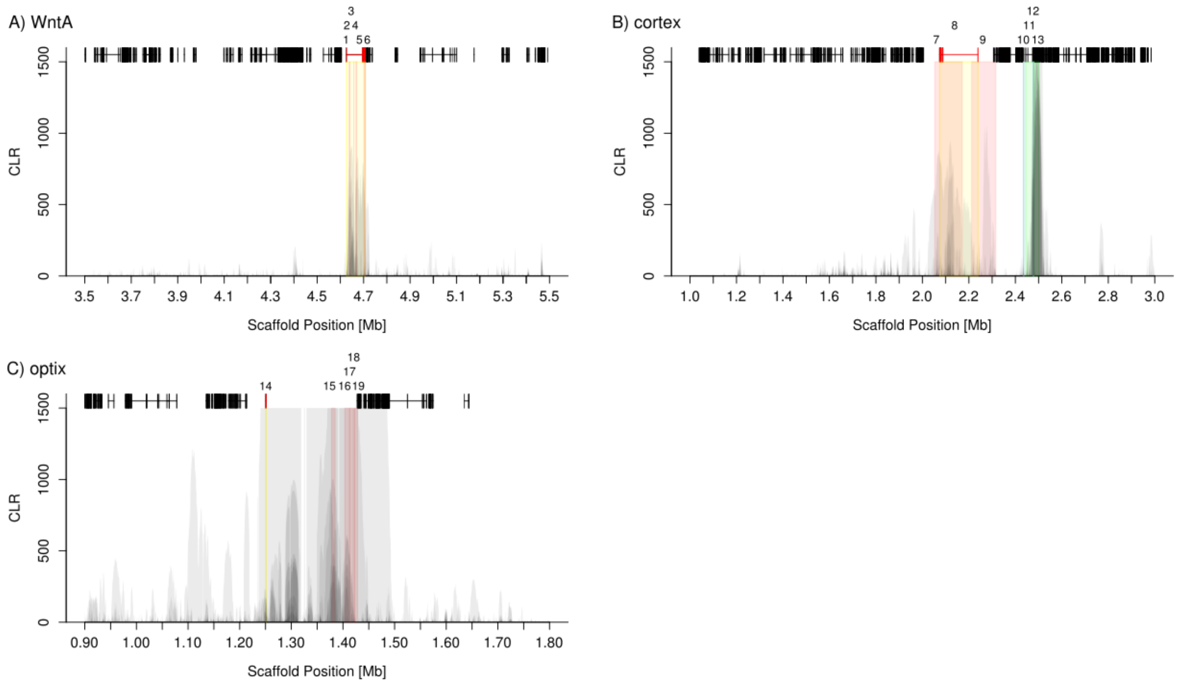

**Supplementary Figure 11. Superposition of SF2 composite likelihood ratio peaks of all *H. erato*-clade populations for each of the four colour pattern regions.** Colour pattern genes (yellow), known CREs (red), and additional genes with evidence for a putative role in colour patterning (blue and green for genes discussed in the main text) are highlighted and assigned a number in the top row. The scale on the x-axes differs and the y-axis is capped at CLR=1,500. **(A)** *wntA* (yellow,1), CREs associated with 'Sd1'(2), 'Sd2'(3), 'St'(4), 'Ly1'(5) and 'Ly2'(6) elements (red)[2]; **(B)** *cortex* (yellow, 8), 'Cr1'(7) and 'Cr2'(9) regions (red )[2], and additional genes with evidence for wing patterning control[7] (blue: 10,12; green; 11 (*washout*/*WAS homologue 1*/HERA000061), 13 (*lethal (2)*/HERA000062); also see Supplementary Table 6; **(C)** *optix* (yellow,14), CREs for 'rays'(15), 'band' Y1(16)/ Y2(18), and 'dennis' D1(17),/ D2(19) elements (red)[2].

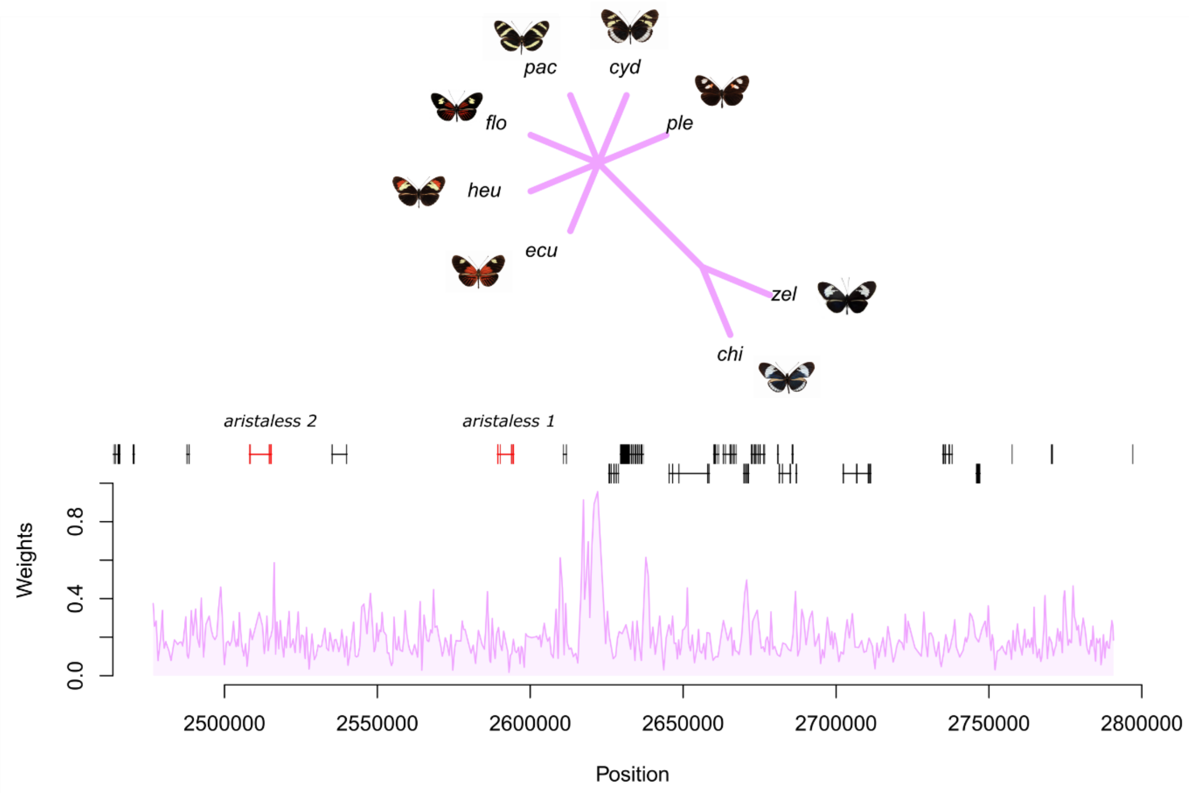

**Supplementary Figure 12. Tree weighting (Twisst[12]) analysis of the *aristaless* genes region.** Topology weightings for topologies clustering the white (chi…*H. c. chioneus*, zel…*H. c. zelinde*) and yellow (ecu…*H. m. ecuadoriensis*, ple…*H. m. plesseni*, heu…*H. heurippa*, flo…*H. t. florencia*, cyd…*H. cydnides*, pac…*H. pachinus*) colour phenotypes (magenta) are shown.

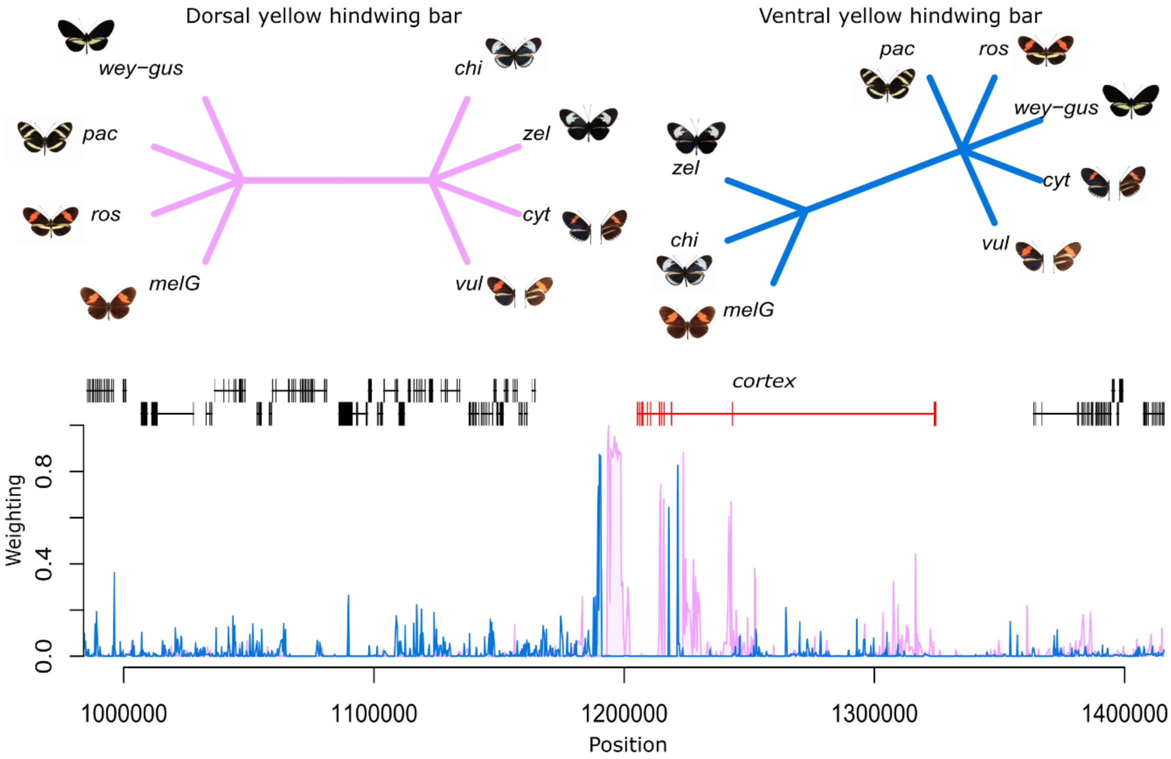

**Supplementary Figure 13. Tree weighting (Twisst[12]) analysis of the *cortex* gene regions.** Topology weightings for topologies clustering the dorsal yellow hindwing bar (magenta) and ventral yellow hindwing bar (blue) phenotypes are shown (cyt…*H. m. cythera*, bur…*H. m burchelli*, nan…*H. m. nanna*, ros…*H. m. rosina*, vul…*H. m.* vulcanus, chi…*H. c. chioneus*, wey…*H. c. weymeri f. weymeri*, gus…*H. c. weymeri f. gustavi*, zel…*H. c. zelinde*, pac…*H. pachinus*).

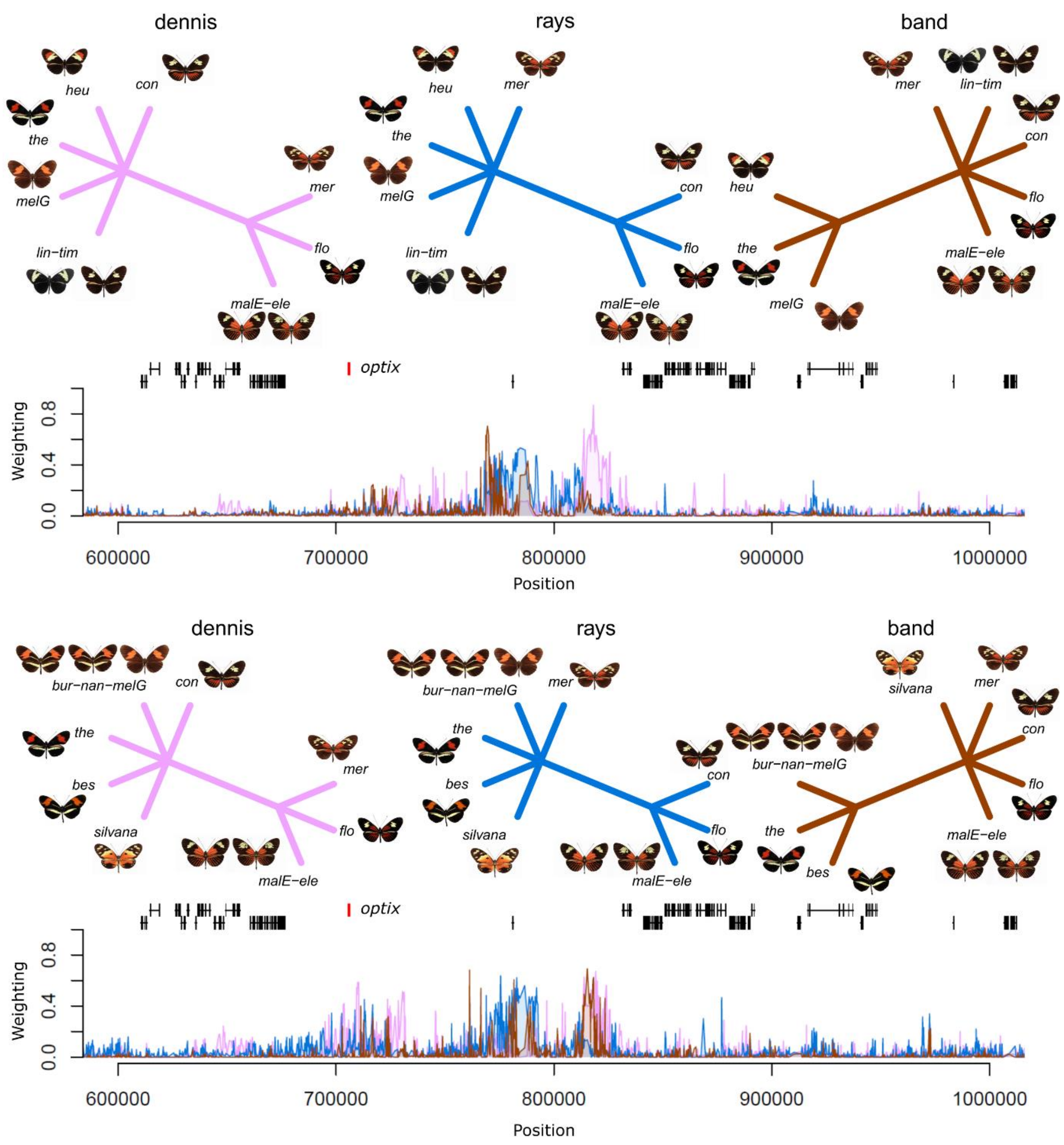

**Supplementary Figure 14. Tree weighting (Twisst[12]) analysis of the *optix* gene regions.** Topology weightings for topologies clustering the dennis (magenta), rays (blue) and band (brown) phenotypes for different geographic regions are shown (bur…*H. m burchelli*, malE…*H. m. malleti* (ECU), melG….*H. m, melpomene* (FG), mer…*H. m. meriana*, nan…*H. m. nanna*, ros…*H. m. rosina*, vul…*H. m.* vulcanus, heu…*H. heurippa*, flo…*H. t. florencia*, lin…*H. t. linaresi*, the…*H. t. thelxinoe*, tim…*H. t. timareta f. timareta*, con…*H. t. timareta f. contigua*, ele…*H. elevatus*, bes…*H. besckei*, silvana…*H. numata silvana*).

**Supplementary Table 1:** Sample information and genotyping statistics for all samples from the *Heliconius melpomene*-clade.

| SequenceID | EarthCapeID | Taxon Name | Country | Latitude | Longitude | Accession | Publication | Called sites | Genotyped sites after filters (%) | Proportion of hetero-zygous genotypes (%) | Data Type (WGS…whole genome re-sequencing, CS…capture sequencing) |
|---|---|---|---|---|---|---|---|---|---|---|---|
| GM110_86 | CAM019819 | *Heliconius besckei* | Brazil | -26.2500 | -49.3836 | pending | | 3,140,439 | 60.08 | 0.80 | CS |
| GM110_83 | CAM019818 | *Heliconius besckei* | Brazil | -26.2500 | -49.3836 | pending | | 3,246,473 | 62.07 | 0.85 | CS |
| GM110_82 | CAM019817 | *Heliconius besckei* | Brazil | -26.2500 | -49.3836 | pending | | 3,220,010 | 61.59 | 0.82 | CS |
| GM123_2 | CAM019816 | *Heliconius besckei* | Brazil | -29.4423 | -50.5798 | pending | | 3,164,769 | 60.55 | 0.79 | CS |
| GM81_3 | CAM019815 | *Heliconius besckei* | Brazil | -25.0462 | -51.5395 | pending | | 3,120,441 | 59.71 | 0.77 | CS |
| GM90_3 | CAM019814 | *Heliconius besckei* | Brazil | -20.1515 | -44.2011 | pending | | 3,189,219 | 60.99 | 0.82 | CS |
| GM89_2 | CAM019813 | *Heliconius besckei* | Brazil | -19.8226 | -43.6762 | pending | | 3,200,508 | 61.26 | 0.74 | CS |
| GM110_84 | CAM019812 | *Heliconius besckei* | Brazil | -26.2500 | -49.3836 | pending | | 3,189,985 | 61.05 | 0.76 | CS |
| GM81_4 | CAM019811 | *Heliconius besckei* | Brazil | -25.0462 | -51.5395 | pending | | 3,319,062 | 63.48 | 0.82 | CS |
| GM88_7 | CAM019810 | *Heliconius besckei* | Brazil | -19.8853 | -43.6644 | pending | | 3,224,622 | 61.69 | 0.79 | CS |
| GM88_1 | CAM019809 | *Heliconius besckei* | Brazil | -19.8853 | -43.6644 | pending | | 3,245,428 | 62.10 | 0.78 | CS |
| chi.CAM584 | CAM000584 | *Heliconius cydno ssp. chioneus* | Panamá | 9.1366 | -79.7236 | SAMEA104585046 | | 3,277,148 | 61.75 | 2.29 | WGS |
| chi.CAM580 | CAM000580 | *Heliconius cydno ssp. chioneus* | Panamá | 9.1366 | -79.7236 | SAMEA104585044 | Van Belleghem *et al.*[13] | 3,534,220 | 66.61 | 2.26 | WGS |
| chi.CAM582 | CAM000582 | *Heliconius cydno ssp. chioneus* | Panamá | 9.1366 | -79.7236 | SAMEA104585045 | Van Belleghem *et al.*[13] | 3,654,172 | 68.82 | 2.33 | WGS |
| chi.CAM585 | CAM000585 | *Heliconius cydno ssp. chioneus* | Panamá | 9.1366 | -79.7236 | SAMEA104585047 | Van Belleghem *et al.*[13] | 3,469,652 | 65.38 | 2.29 | WGS |
| chi.CAM586 | CAM000586 | *Heliconius cydno ssp. chioneus* | Panamá | 9.1366 | -79.7236 | SAMEA104585048 | Van Belleghem *et al.*[13] | 3,538,969 | 66.67 | 2.31 | WGS |
| chi.CAM25091 | CAM025091 | *Heliconius cydno ssp. chioneus* | Panamá | 9.1200 | -79.7020 | SAMEA104585050 | Van Belleghem *et al.*[13] | 3,449,544 | 64.97 | 2.33 | WGS |
| chi.CAM25137 | CAM025137 | *Heliconius cydno ssp. chioneus* | Panamá | 9.1200 | -79.7020 | SAMEA104585051 | Van Belleghem *et al.*[13] | 3,548,034 | 66.82 | 2.35 | WGS |
| chi.CJ553 | CAM000553 | *Heliconius cydno ssp. chioneus* | Panamá | 9.1366 | -79.7236 | SAMEA1919256 | Martin *et al.*[14] | 3,666,518 | 69.06 | 2.33 | WGS |
| chi.CJ565 | CAM000565 | *Heliconius cydno ssp. chioneus* | Panamá | 9.1714 | -79.7573 | SAMEA1919262 | Martin *et al.*[14] | 3,799,061 | 71.53 | 2.36 | WGS |
| chi.CJ560 | CAM000560 | *Heliconius cydno ssp. chioneus* | Panamá | 9.1366 | -79.7236 | SAMEA1919265 | Martin *et al.*[14] | 3,694,123 | 69.60 | 2.30 | WGS |
| chi.CJ564 | CAM000564 | *Heliconius cydno ssp. chioneus* | Panamá | 9.1366 | -79.7236 | SAMEA1919278 | Martin *et al.*[14] | 3,704,117 | 69.75 | 2.36 | WGS |
| cor.CS3 | CS002167 | *Heliconius cydno ssp. cordula* | Venezuela | 7.9406 | -72.3406 | SAMEA104585052 | Van Belleghem *et al.*[13] | 3,783,632 | 71.33 | 2.24 | WGS |
| cor.CS4 | CS002258 | *Heliconius cydno ssp. cordula* | Venezuela | 7.9406 | -72.3406 | SAMEA104585053 | Van Belleghem *et al.*[13] | 3,775,592 | 71.22 | 2.18 | WGS |
| cor.STRI7 | STRI_007 | *Heliconius cydno ssp. cordula* | Venezuela | 7.7980 | -72.2047 | SAMEA3670511 | Wallbank *et al.*[8] | 3,780,111 | 71.17 | 2.37 | WGS |
| CS2737 | CS002737 | *Heliconius cydno ssp. cydnides* | Colombia | 3.8665 | -76.3834 | pending | | 3,818,575 | 72.17 | 1.99 | CS |
| CS2729 | CS002729 | *Heliconius cydno ssp. cydnides* | Colombia | 3.8525 | -76.4269 | pending | | 3,822,998 | 72.30 | 1.93 | CS |
| CS2727 | CS002727 | *Heliconius cydno ssp. cydnides* | Colombia | 3.8525 | -76.4269 | pending | | 3,822,358 | 72.29 | 1.93 | CS |
| CS2748 | CS002748 | *Heliconius cydno ssp. cydnides* | Colombia | 3.8665 | -76.3834 | pending | | 3,812,649 | 72.08 | 1.96 | CS |
| CS2740 | CS002740 | *Heliconius cydno ssp. cydnides* | Colombia | 3.8665 | -76.3834 | pending | | 3,857,734 | 72.86 | 2.06 | CS |
| CS2738 | CS002738 | *Heliconius cydno ssp. cydnides* | Colombia | 3.8665 | -76.3834 | pending | | 3,856,673 | 72.84 | 2.06 | CS |
| CS2724 | CS002724 | *Heliconius cydno ssp. cydnides* | Colombia | 3.8525 | -76.4269 | pending | | 3,841,436 | 72.58 | 2.03 | CS |
| CS2731 | CS002731 | *Heliconius cydno ssp. cydnides* | Colombia | 3.8525 | -76.4269 | pending | | 3,846,873 | 72.68 | 2.02 | CS |
| CS2739 | CS002739 | *Heliconius cydno ssp. cydnides* | Colombia | 3.8665 | -76.3834 | pending | | 3,825,523 | 72.33 | 1.96 | CS |
| CS2732 | CS002732 | *Heliconius cydno ssp. cydnides* | Colombia | 3.8525 | -76.4269 | pending | | 3,840,359 | 72.72 | 1.81 | CS |

| SequenceID | EarthCapeID | Taxon Name | Country | Latitude | Longitude | Accession | Publication | Called sites | Genotyped sites after filters (%) | Proportion of hetero-zygous genotypes (%) | Data Type (WGS…whole genome re-sequencing, CS…capture sequencing) |
|---|---|---|---|---|---|---|---|---|---|---|---|
| CS936 | CS000936 | *Heliconius cydno ssp. weymeri f. gustavi* | Colombia | 3.4834 | -76.6158 | pending | | 3,840,229 | 72.86 | 1.62 | CS |
| CS2492 | CS002492 | *Heliconius cydno ssp. weymeri f. gustavi* | Colombia | 3.3242 | -76.6364 | pending | | 3,873,417 | 73.27 | 1.91 | CS |
| CS2511 | CS002511 | *Heliconius cydno ssp. weymeri f. gustavi* | Colombia | 3.6106 | -76.5986 | pending | | 3,866,621 | 73.24 | 1.7 | CS |
| CS2506 | CS002506 | *Heliconius cydno ssp. weymeri f. gustavi* | Colombia | 3.6106 | -76.5986 | pending | | 3,802,771 | 71.98 | 1.85 | CS |
| CS1914 | CS001914 | *Heliconius cydno ssp. weymeri f. gustavi* | Colombia | 2.5133 | -76.5942 | pending | | 3,821,081 | 72.38 | 1.78 | CS |
| CS953 | CS000953 | *Heliconius cydno ssp. weymeri f. gustavi* | Colombia | 3.4834 | -76.6158 | pending | | 3,838,035 | 72.73 | 1.74 | CS |
| CS950 | CS000950 | *Heliconius cydno ssp. weymeri f. gustavi* | Colombia | 3.4834 | -76.6158 | pending | | 3,823,525 | 72.35 | 1.88 | CS |
| CS1926 | CS001926 | *Heliconius cydno ssp. weymeri f. gustavi* | Colombia | 2.5133 | -76.5942 | pending | | 3,862,132 | 73.14 | 1.80 | CS |
| CS2503 | CS002503 | *Heliconius cydno ssp. weymeri f. gustavi* | Colombia | 2.5133 | -76.5942 | pending | | 3,845,743 | 72.86 | 1.76 | CS |
| CS2504 | CS002504 | *Heliconius cydno ssp. weymeri f. gustavi* | Colombia | 3.6106 | -76.5986 | pending | | 3,835,440 | 72.40 | 2.11 | CS |
| CS982 | CS000982 | *Heliconius cydno ssp. weymeri f. weymeri* | Colombia | 3.4834 | -76.6158 | pending | | 3,820,791 | 72.47 | 1.65 | CS |
| CS942 | CS000942 | *Heliconius cydno ssp. weymeri f. weymeri* | Colombia | 3.4834 | -76.6158 | pending | | 3,880,634 | 73.44 | 1.87 | CS |
| CS2996 | CS002996 | *Heliconius cydno ssp. weymeri f. weymeri* | Colombia | 3.4834 | -76.6158 | pending | | 3,827,233 | 72.48 | 1.80 | CS |
| CS2998 | CS002998 | *Heliconius cydno ssp. weymeri f. weymeri* | Colombia | 3.4834 | -76.6158 | pending | | 3,859,920 | 72.97 | 1.97 | CS |
| CS943 | CS000943 | *Heliconius cydno ssp. weymeri f. weymeri* | Colombia | 3.4834 | -76.6158 | pending | | 3,852,736 | 72.88 | 1.90 | CS |
| CS951 | CS000951 | *Heliconius cydno ssp. weymeri f. weymeri* | Colombia | 3.4834 | -76.6158 | pending | | 3,857,073 | 73.09 | 1.73 | CS |
| CS960 | CS000960 | *Heliconius cydno ssp. weymeri f. weymeri* | Colombia | 3.4834 | -76.6158 | pending | | 3,827,645 | 72.52 | 1.75 | CS |
| CS945 | CS000945 | *Heliconius cydno ssp. weymeri f. weymeri* | Colombia | 3.4834 | -76.6158 | pending | | 3,879,725 | 73.38 | 1.93 | CS |
| CS2542 | CS002542 | *Heliconius cydno ssp. weymeri f. weymeri* | Colombia | 3.3242 | -76.6364 | pending | | 3,844,853 | 72.78 | 1.84 | CS |
| CS939 | CS000939 | *Heliconius cydno ssp. weymeri f. weymeri* | Colombia | 3.4834 | -76.6158 | pending | | 3,826,346 | 72.55 | 1.68 | CS |
| zel.CS1 | CS002242 | *Heliconius cydno ssp. zelinde* | Colombia | 3.9394 | -77.3689 | SAMEA104106540 | Enciso-Romero *et al.*[6] | 3,921,772 | 74.40 | 1.62 | WGS |
| zel.CS2 | CS002261 | *Heliconius cydno ssp. zelinde* | Colombia | 3.9394 | -77.3689 | SAMEA104106542 | Enciso-Romero *et al.*[6] | 3,932,964 | 74.03 | 2.40 | WGS |
| zel.CS30 | CS002260 | *Heliconius cydno ssp. zelinde* | Colombia | 3.9394 | -77.3689 | SAMEA104106543 | Enciso-Romero *et al.*[6] | 3,958,948 | 74.78 | 2.05 | WGS |
| zel.CS1028 | CS001028 | *Heliconius cydno ssp. zelinde* | Colombia | 3.9394 | -77.3689 | SAMEA104585054 | Van Belleghem *et al.*[13] | 3,901,819 | 73.44 | 2.39 | WGS |
| zel.CS1029 | CS001029 | *Heliconius cydno ssp. zelinde* | Colombia | 3.9583 | -77.3733 | SAMEA104585055 | Van Belleghem *et al.*[13] | 3,571,047 | 67.27 | 2.32 | WGS |
| zel.CS1030 | CS001030 | *Heliconius cydno ssp. zelinde* | Colombia | 3.9583 | -77.3733 | SAMEA104585056 | Van Belleghem *et al.*[13] | 3,559,106 | 67.05 | 2.30 | WGS |
| zel.CS1033 | CS001033 | *Heliconius cydno ssp. zelinde* | Colombia | 3.9394 | -77.3689 | SAMEA104585057 | Van Belleghem *et al.*[13] | 3,644,672 | 68.61 | 2.38 | WGS |

| SequenceID | EarthCapeID | Taxon Name | Country | Latitude | Longitude | Accession | Publication | Called sites | Genotyped sites after filters (%) | Proportion of hetero-zygous genotypes (%) | Data Type (WGS…whole genome re-sequencing, CS…capture sequencing) |
|---|---|---|---|---|---|---|---|---|---|---|---|
| zel.CS1035 | CS001035 | *Heliconius cydno ssp. zelinde* | Colombia | 3.9394 | -77.3689 | SAMEA104585058 | Van Belleghem *et al.*[13] | 3,830,391 | 72.15 | 2.33 | WGS |
| zel.CS273 | CS000273 | *Heliconius cydno ssp. zelinde* | Colombia | 3.9394 | -77.3689 | SAMEA104585059 | Van Belleghem *et al.*[13] | 3,841,140 | 72.49 | 2.14 | WGS |
| zel.CS2262 | CS002262 | *Heliconius cydno ssp. zelinde* | Colombia | 3.9394 | -77.3689 | SAMEA3670517 | Enciso-Romero *et al.*[6] | 3,249,828 | 61.23 | 2.29 | WGS |
| PS17434 | CAM017434 | *Heliconius elevatus* | Ecuador | -1.2908 | -77.8419 | pending | | 3,420,534 | 64.04 | 2.92 | CS |
| PS17411 | CAM017411 | *Heliconius elevatus* | Ecuador | -1.0157 | -77.5975 | pending | | 3,442,881 | 64.69 | 2.57 | CS |
| PS17098 | CAM017098 | *Heliconius elevatus* | Ecuador | -1.0614 | -77.6684 | pending | | 3,403,379 | 63.73 | 2.89 | CS |
| PS16984 | CAM016984 | *Heliconius elevatus* | Ecuador | -1.1878 | -77.8311 | pending | | 3,418,205 | 64.08 | 2.78 | CS |
| PS16933 | CAM016933 | *Heliconius elevatus* | Ecuador | -1.1156 | -77.7783 | pending | | 3,409,748 | 63.99 | 2.68 | CS |
| PS16781 | CAM016781 | *Heliconius elevatus* | Ecuador | -1.3382 | -77.8354 | pending | | 3,403,394 | 63.81 | 2.78 | CS |
| PS16776 | CAM016776 | *Heliconius elevatus* | Ecuador | -1.2519 | -77.8196 | pending | | 3,413,732 | 63.89 | 2.95 | CS |
| PS16626 | CAM016626 | *Heliconius elevatus* | Ecuador | -1.1156 | -77.7783 | pending | | 3,400,423 | 63.68 | 2.89 | CS |
| PS16607 | CAM016607 | *Heliconius elevatus* | Ecuador | -1.1156 | -77.7783 | pending | | 3,423,138 | 64.13 | 2.85 | CS |
| PS16596 | CAM016596 | *Heliconius elevatus* | Ecuador | -1.1156 | -77.7783 | pending | | 3,411,721 | 63.91 | 2.86 | CS |
| PS16439 | CAM016439 | *Heliconius elevatus* | Ecuador | -1.1878 | -77.8311 | pending | | 3,375,448 | 63.24 | 2.84 | CS |
| PS16216 | CAM016216 | *Heliconius elevatus* | Ecuador | -1.1156 | -77.7783 | pending | | 3,409,726 | 63.83 | 2.92 | CS |
| PS16209 | CAM016209 | *Heliconius elevatus* | Ecuador | -1.1156 | -77.7783 | pending | | 3,393,480 | 63.77 | 2.55 | CS |
| PS16207 | CAM016207 | *Heliconius elevatus* | Ecuador | -1.1156 | -77.7783 | pending | | 3,417,159 | 63.96 | 2.93 | CS |
| CAM008861 | CAM008861 | *Heliconius hecale* | Panamá | 7.7568 | -77.6841 | pending | | 3,467,332 | 65.32 | 2.30 | CS |
| CAM008683 | CAM008683 | *Heliconius hecale* | Peru | -6.2897 | -76.2289 | pending | | 3,431,006 | 64.70 | 2.22 | CS |
| heu.CS18 | CS00CH18 | *Heliconius heurippa* | Colombia | 4.1750 | -73.6781 | on rcs; pending | | 3,999,662 | 76.08 | 1.36 | WGS |
| CS2014 | CS002014 | *Heliconius heurippa* | Colombia | 4.1750 | -73.6781 | pending | | 3,876,614 | 73.70 | 1.42 | CS |
| CS2420 | CS002420 | *Heliconius heurippa* | Colombia | 4.1750 | -73.6781 | pending | | 3,852,612 | 73.37 | 1.24 | CS |
| CS3394 | CS003394 | *Heliconius heurippa* | Colombia | 4.1750 | -73.6781 | pending | | 3,843,551 | 73.12 | 1.35 | CS |
| CS3424 | CS003424 | *Heliconius heurippa* | Colombia | 4.1750 | -73.6781 | pending | | 3,837,343 | 73.07 | 1.26 | CS |
| CS1510 | CS001510 | *Heliconius heurippa* | Colombia | 4.1750 | -73.6781 | pending | | 3,838,220 | 72.95 | 1.45 | CS |
| CS3392 | CS003392 | *Heliconius heurippa* | Colombia | 4.1750 | -73.6781 | pending | | 3,846,603 | 73.24 | 1.26 | CS |
| CS3843 | CS003843 | *Heliconius heurippa* | Colombia | 3.5667 | -74.0761 | pending | | 3,849,937 | 73.16 | 1.45 | CS |
| CS3391 | CS003391 | *Heliconius heurippa* | Colombia | 4.1750 | -73.6781 | pending | | 3,880,909 | 73.93 | 1.22 | CS |
| CS3696 | CS003696 | *Heliconius heurippa* | Colombia | 4.1750 | -73.6781 | pending | | 3,812,208 | 72.53 | 1.35 | CS |
| CS3393 | CS003393 | *Heliconius heurippa* | Colombia | 4.1750 | -73.6781 | pending | | 3,864,380 | 73.45 | 1.44 | CS |
| CS3423 | CS003423 | *Heliconius heurippa* | Colombia | 4.1750 | -73.6781 | pending | | 3,887,467 | 73.92 | 1.40 | CS |
| CS1353 | CS001353 | *Heliconius heurippa* | Colombia | 4.1750 | -73.6781 | pending | | 3,877,700 | 73.79 | 1.32 | CS |
| CS2012 | CS002012 | *Heliconius heurippa* | Colombia | 4.1750 | -73.6781 | pending | | 3,880,389 | 74.07 | 1.01 | CS |
| CS2618 | CS002618 | *Heliconius heurippa* | Colombia | 4.1750 | -73.6781 | pending | | 3,837,340 | 73.02 | 1.33 | CS |
| CS3007 | CS003007 | *Heliconius heurippa* | Colombia | 4.1750 | -73.6781 | pending | | 3,858,645 | 73.33 | 1.45 | CS |
| CS1352 | CS001352 | *Heliconius heurippa* | Colombia | 4.1750 | -73.6781 | pending | | 3,858,710 | 73.53 | 1.18 | CS |
| CS3844 | CS003844 | *Heliconius heurippa* | Colombia | 3.5667 | -74.0761 | pending | | 3,848,242 | 73.03 | 1.60 | CS |
| CS3397 | CS003397 | *Heliconius heurippa* | Colombia | 4.1750 | -73.6781 | pending | | 3,881,108 | 73.43 | 1.89 | CS |
| CS2204 | CS002204 | *Heliconius heurippa* | Colombia | 4.1750 | -73.6781 | pending | | 3,824,158 | 72.79 | 1.30 | CS |
| heu.CS20 | CS00CH21 | *Heliconius heurippa* | Colombia | 4.1750 | -73.6781 | SAMEA104585060 | Van Belleghem *et al.*[13] | 3,926,742 | 74.58 | 1.51 | WGS |

| SequenceID | EarthCapeID | Taxon Name | Country | Latitude | Longitude | Accession | Publication | Called sites | Genotyped sites after filters (%) | Proportion of hetero-zygous genotypes (%) | Data Type (WGS…whole genome re-sequencing, CS…capture sequencing) |
|---|---|---|---|---|---|---|---|---|---|---|---|
| CS000CH9 | CS000CH9 | *Heliconius heurippa* | Colombia | 4.1750 | -73.6781 | SAMEA1322902 | The Heliconius Consortium[15]; RAD | 4,080,230 | 77.72 | 1.23 | WGS |
| heu.STRI2 | STRI_002 | *Heliconius heurippa* | Colombia | 4.1750 | -73.6781 | SAMEA3670535 | Wallbank *et al.*[8] | 3,754,542 | 71.42 | 1.36 | WGS |
| CS2013 | CS002013 | *Heliconius heurippa* | Colombia | 4.1750 | -73.6781 | pending | | 3,883,113 | 73.32 | 2.09 | CS |
| CAM002700 | CAM002700 | *Heliconius ismenius* | Panamá | 9.1366 | -79.7236 | pending | | 3,268,849 | 62.18 | 1.36 | CS |
| CAM002517 | CAM002517 | *Heliconius ismenius* | Panamá | 9.1366 | -79.7236 | pending | | 3,396,394 | 64.28 | 1.86 | CS |
| melP.HGC1 | gen_ref | *Heliconius melpomene reference genome* | - | - | - | SAMN00794386 | The Heliconius Consortium[15] | 4,418,717 | 84.76 | 0.53 | WGS |
| agl.JM108 | JM-09-108 | *Heliconius melpomene ssp. aglaope* | Peru | -5.9103 | -76.2256 | SAMEA1919251 | Martin *et al.*[14] | 3,705,462 | 69.73 | 2.42 | WGS |
| agl.JM572 | JM-11-572 | *Heliconius melpomene ssp. aglaope* | Peru | -5.9717 | -76.2319 | SAMEA1919259 | Martin *et al.*[14] | 3,777,573 | 71.08 | 2.43 | WGS |
| agl.JM112 | JM-09-112 | *Heliconius melpomene ssp. aglaope* | Peru | -5.9103 | -76.2256 | SAMEA1919264 | Martin *et al.*[14] | 3,785,847 | 71.25 | 2.41 | WGS |
| agl.JM569 | JM-11-569 | *Heliconius melpomene ssp. aglaope* | Peru | -5.9717 | -76.2319 | SAMEA1919274 | Martin *et al.*[14] | 3,823,969 | 71.97 | 2.40 | WGS |
| MJ12-3414 | MJ12-3414 | *Heliconius melpomene ssp. amaryllis* | Peru | -6.4540 | -76.3002 | pending | | 3,537,813 | 66.92 | 1.91 | CS |
| MJ12-3397 | MJ12-3397 | *Heliconius melpomene ssp. amaryllis* | Peru | -6.4528 | -76.2862 | pending | | 3,511,652 | 66.44 | 1.89 | CS |
| MJ12-3302 | MJ12-3302 | *Heliconius melpomene ssp. amaryllis* | Peru | -6.4528 | -76.2862 | pending | | 3,586,055 | 67.84 | 1.90 | CS |
| MJ12-3074 | MJ12-3074 | *Heliconius melpomene ssp. amaryllis* | Peru | -6.4528 | -76.2862 | pending | | 3,356,271 | 63.55 | 1.81 | CS |
| MJ12-3371 | MJ12-3371 | *Heliconius melpomene ssp. amaryllis* | Peru | -6.4528 | -76.2862 | pending | | 3,513,745 | 66.47 | 1.90 | CS |
| MJ12-3442 | MJ12-3442 | *Heliconius melpomene ssp. amaryllis* | Peru | -6.4547 | -76.2994 | pending | | 3,554,687 | 67.20 | 1.97 | CS |
| MJ12-3416 | MJ12-3416 | *Heliconius melpomene ssp. amaryllis* | Peru | -6.4537 | -76.2981 | pending | | 3,491,823 | 65.99 | 2.00 | CS |
| MJ12-3393 | MJ12-3393 | *Heliconius melpomene ssp. amaryllis* | Peru | -6.4528 | -76.2862 | pending | | 3,584,960 | 67.85 | 1.86 | CS |
| MJ12-3417 | MJ12-3417 | *Heliconius melpomene ssp. amaryllis* | Peru | -6.4537 | -76.2981 | pending | | 3,547,133 | 67.04 | 1.99 | CS |
| MJ12-3267 | MJ12-3267 | *Heliconius melpomene ssp. amaryllis* | Peru | -6.4528 | -76.2862 | pending | | 3,360,799 | 63.64 | 1.80 | CS |
| MJ12-3474 | MJ12-3474 | *Heliconius melpomene ssp. amaryllis* | Peru | -6.4537 | -76.2981 | pending | | 3,575,687 | 67.61 | 1.94 | CS |
| MJ12-3467 | MJ12-3467 | *Heliconius melpomene ssp. amaryllis* | Peru | -6.4528 | -76.2862 | pending | | 3,555,664 | 67.22 | 1.97 | CS |
| MJ12-3482 | MJ12-3482 | *Heliconius melpomene ssp. amaryllis* | Peru | -6.4524 | -76.2869 | pending | | 3,568,507 | 67.46 | 1.97 | CS |
| MJ12-3306 | MJ12-3306 | *Heliconius melpomene ssp. amaryllis* | Peru | -6.4572 | -76.2986 | pending | | 3,570,395 | 67.53 | 1.91 | CS |
| MJ11-3018 | MJ11-3018 | *Heliconius melpomene ssp. amaryllis* | Peru | -6.4567 | -76.2845 | pending | | 3,409,538 | 64.54 | 1.85 | CS |
| MJ12-3418 | MJ12-3418 | *Heliconius melpomene ssp. amaryllis* | Peru | -6.4537 | -76.2981 | pending | | 3,577,439 | 67.60 | 2.01 | CS |
| MJ12-3585 | MJ12-3585 | *Heliconius melpomene ssp. amaryllis* | Peru | -6.4528 | -76.2862 | pending | | 3,428,919 | 64.91 | 1.83 | CS |
| MJ12-3466 | MJ12-3466 | *Heliconius melpomene ssp. amaryllis* | Peru | -6.4530 | -76.2876 | pending | | 3,580,238 | 67.72 | 1.92 | CS |

| SequenceID | EarthCapeID | Taxon Name | Country | Latitude | Longitude | Accession | Publication | Called sites | Genotyped sites after filters (%) | Proportion of hetero-zygous genotypes (%) | Data Type (WGS…whole genome re-sequencing, CS…capture sequencing) |
|---|---|---|---|---|---|---|---|---|---|---|---|
| MJ12-3137 | MJ12-3137 | *Heliconius melpomene ssp. amaryllis* | Peru | -6.4547 | -76.2994 | pending | | 3,195,835 | 60.21 | 2.30 | CS |
| MJ12-3394 | MJ12-3394 | *Heliconius melpomene ssp. amaryllis* | Peru | -6.4528 | -76.2862 | pending | | 3,426,831 | 64.84 | 1.89 | CS |
| MJ12-3453 | MJ12-3453 | *Heliconius melpomene ssp. amaryllis* | Peru | -6.4537 | -76.2981 | pending | | 3,521,043 | 66.65 | 1.84 | CS |
| MJ12-3392 | MJ12-3392 | *Heliconius melpomene ssp. amaryllis* | Peru | -6.4528 | -76.2862 | pending | | 3,585,104 | 67.79 | 1.94 | CS |
| MJ11-3021 | MJ11-3021 | *Heliconius melpomene ssp. amaryllis* | Peru | -6.4555 | -76.2843 | pending | | 3,354,395 | 63.45 | 1.92 | CS |
| MJ11-3024 | MJ11-3024 | *Heliconius melpomene ssp. amaryllis* | Peru | -6.4555 | -76.2843 | pending | | 3,399,058 | 64.35 | 1.83 | CS |
| MJ11-3023 | MJ11-3023 | *Heliconius melpomene ssp. amaryllis* | Peru | -6.4555 | -76.2843 | pending | | 3,388,398 | 64.12 | 1.86 | CS |
| MJ12-3483 | MJ12-3483 | *Heliconius melpomene ssp. amaryllis* | Peru | -6.4528 | -76.2862 | pending | | 3,398,807 | 64.32 | 1.87 | CS |
| MJ12-3396 | MJ12-3396 | *Heliconius melpomene ssp. amaryllis* | Peru | -6.4528 | -76.2862 | pending | | 3,575,104 | 67.60 | 1.94 | CS |
| MJ12-3130 | MJ12-3130 | *Heliconius melpomene ssp. amaryllis* | Peru | -6.4572 | -76.2986 | pending | | 3,475,149 | 65.79 | 1.83 | CS |
| ama.MJ11-3188 | MJ11-3188 | *Heliconius melpomene ssp. amaryllis* | Peru | -5.6728 | -77.7195 | SAMEA104585061 | Van Belleghem *et al.*[13] | 3,432,288 | 64.83 | 2.05 | WGS |
| ama.MJ11-3189 | MJ11-3189 | *Heliconius melpomene ssp. amaryllis* | Peru | -5.6728 | -77.7195 | SAMEA104585062 | Van Belleghem *et al.*[13] | 3,502,389 | 66.25 | 1.92 | WGS |
| ama.MJ11-3202 | MJ11-3202 | *Heliconius melpomene ssp. amaryllis* | Peru | -5.6745 | -77.6711 | SAMEA104585063 | Van Belleghem *et al.*[13] | 3,365,693 | 63.43 | 2.28 | WGS |
| ama.MJ12-3217 | MJ12-3217 | *Heliconius melpomene ssp. amaryllis* | Peru | -6.4547 | -76.2994 | SAMEA104585064 | Van Belleghem *et al.*[13] | 3,742,548 | 70.64 | 2.13 | WGS |
| ama.MJ12-3258 | MJ12-3258 | *Heliconius melpomene ssp. amaryllis* | Peru | -6.4530 | -76.2876 | SAMEA104585065 | Van Belleghem *et al.*[13] | 3,648,380 | 68.72 | 2.32 | WGS |
| ama.MJ12-3301 | MJ12-3301 | *Heliconius melpomene ssp. amaryllis* | Peru | -6.4528 | -76.2862 | SAMEA104585066 | Van Belleghem *et al.*[13] | 3,655,176 | 69.06 | 2.02 | WGS |
| ama.JM160 | JM-11-160 | *Heliconius melpomene ssp. amaryllis* | Peru | -6.4645 | -76.3514 | SAMEA1919261 | Martin *et al.*[14] | 3,911,236 | 73.79 | 2.17 | WGS |
| ama.JM216 | JM-09-216 | *Heliconius melpomene ssp. amaryllis* | Peru | -5.6745 | -77.6748 | SAMEA1919261 | Martin *et al.*[14] | 3,823,200 | 72.18 | 2.10 | WGS |
| ama.JM48 | JM-11-48 | *Heliconius melpomene ssp. amaryllis* | Peru | -6.0960 | -76.9774 | SAMEA1919269 | Martin *et al.*[14] | 3,945,310 | 74.40 | 2.21 | WGS |
| ama.JM293 | JM-11-293 | *Heliconius melpomene ssp. amaryllis* | Peru | -6.4667 | -76.3347 | SAMEA1919277 | Martin *et al.*[14] | 3,935,343 | 74.27 | 2.13 | WGS |
| GM184_4 | CAM019822 | *Heliconius melpomene ssp. burchelli* | Brazil | -6.9639 | -46.6799 | pending | | 3,841,610 | 72.76 | 1.78 | CS |
| GM184_3 | CAM019821 | *Heliconius melpomene ssp. burchelli* | Brazil | -6.9639 | -46.6799 | pending | | 3,852,678 | 72.93 | 1.84 | CS |
| GM184_1 | CAM019820 | *Heliconius melpomene ssp. burchelli* | Brazil | -6.9639 | -46.6799 | pending | | 3,852,984 | 72.96 | 1.81 | CS |
| s15N005 | 15N005 | *Heliconius melpomene ssp. cythera* | Ecuador | 0.1753 | -78.9075 | pending | | 4,051,459 | 76.58 | 1.98 | CS |
| s15N006 | 15N006 | *Heliconius melpomene ssp. cythera* | Ecuador | 0.1937 | -78.8587 | pending | | 4,062,354 | 76.87 | 1.88 | CS |
| s15N009 | 15N009 | *Heliconius melpomene ssp. cythera* | Ecuador | 0.1753 | -78.9075 | pending | | 4,097,078 | 77.41 | 2.02 | CS |
| s15N022 | 15N022 | *Heliconius melpomene ssp. cythera* | Ecuador | 0.1850 | -78.8530 | pending | | 4,061,051 | 76.72 | 2.04 | CS |

| SequenceID | EarthCapeID | Taxon Name | Country | Latitude | Longitude | Accession | Publication | Called sites | Genotyped sites after filters (%) | Proportion of hetero-zygous genotypes (%) | Data Type (WGS…whole genome re-sequencing, CS…capture sequencing) |
|---|---|---|---|---|---|---|---|---|---|---|---|
| 14N594 | 14N594 | *Heliconius melpomene ssp. cythera* | Ecuador | 0.1659 | -78.8882 | pending | | 3,984,318 | 75.31 | 1.98 | CS |
| s15N008 | 15N008 | *Heliconius melpomene ssp. cythera* | Ecuador | 0.1850 | -78.8530 | pending | | 4,065,406 | 76.90 | 1.91 | CS |
| s15N023 | 15N023 | *Heliconius melpomene ssp. cythera* | Ecuador | 0.1937 | -78.8587 | pending | | 4,074,163 | 77.15 | 1.81 | CS |
| s15N020 | 15N020 | *Heliconius melpomene ssp. cythera* | Ecuador | 0.1850 | -78.8530 | pending | | 4,064,674 | 76.85 | 1.96 | CS |
| s15N004 | 15N004 | *Heliconius melpomene ssp. cythera* | Ecuador | 0.1753 | -78.9075 | pending | | 4,077,502 | 77.11 | 1.93 | CS |
| CAM002856_ERR1143591 | CAM002856 | *Heliconius melpomene ssp. cythera* | Ecuador | -0.3197 | -79.3370 | SAMEA3670542 | Wallbank *et al.*[8] | 3,223,375 | 60.67 | 2.39 | WGS |
| CAM009116 | CAM009116 | *Heliconius melpomene ssp. ecuadoriensis* | Ecuador | -4.0439 | -78.9861 | pending | | 3,878,044 | 73.89 | 1.20 | CS |
| CAM009119 | CAM009119 | *Heliconius melpomene ssp. ecuadoriensis* | Ecuador | -4.0439 | -78.9861 | pending | | 3,857,233 | 72.72 | 2.23 | CS |
| CAM009118 | CAM009118 | *Heliconius melpomene ssp. ecuadoriensis* | Ecuador | -4.0439 | -78.9861 | pending | | 3,883,754 | 73.22 | 2.23 | CS |
| CAM009115 | CAM009115 | *Heliconius melpomene ssp. ecuadoriensis* | Ecuador | -4.0439 | -78.9861 | pending | | 3,866,026 | 72.92 | 2.19 | CS |
| CAM009114 | CAM009114 | *Heliconius melpomene ssp. ecuadoriensis* | Ecuador | -4.0439 | -78.9861 | pending | | 3,887,362 | 73.28 | 2.24 | CS |
| CAM009113 | CAM009113 | *Heliconius melpomene ssp. ecuadoriensis* | Ecuador | -4.0439 | -78.9861 | pending | | 3,864,300 | 72.87 | 2.22 | CS |
| CAM002432 | CAM002432 | *Heliconius melpomene ssp. ecuadoriensis* | Ecuador | -4.0653 | -78.9587 | pending | | 3,869,756 | 73.02 | 2.15 | CS |
| CAM002430 | CAM002430 | *Heliconius melpomene ssp. ecuadoriensis* | Ecuador | -4.0653 | -78.9587 | pending | | 3,910,420 | 73.75 | 2.20 | CS |
| CAM002417 | CAM002417 | *Heliconius melpomene ssp. ecuadoriensis* | Ecuador | -4.0653 | -78.9587 | pending | | 3,874,823 | 73.06 | 2.23 | CS |
| CAM009112 | CAM009112 | *Heliconius melpomene ssp. ecuadoriensis* | Ecuador | -4.0439 | -78.9861 | SAMEA1322914; pending | The Heliconius Consortium[15]; RAD; SureSelect | 3,872,864 | 73.07 | 2.17 | CS |
| CS3447 | CS003447 | *Heliconius melpomene ssp. malleti* | Colombia | 1.8033 | -75.6553 | pending | | 3,738,051 | 70.61 | 2.05 | CS |
| CS3706 | CS003706 | *Heliconius melpomene ssp. malleti* | Colombia | 1.8033 | -75.6553 | pending | | 3,791,670 | 71.98 | 1.56 | CS |
| CS3542 | CS003542 | *Heliconius melpomene ssp. malleti* | Colombia | 1.7506 | -75.6319 | pending | | 3,790,119 | 71.49 | 2.19 | CS |
| CS3728 | CS003728 | *Heliconius melpomene ssp. malleti* | Colombia | 1.8033 | -75.6553 | pending | | 3,779,425 | 71.29 | 2.18 | CS |
| CS583 | CS000583 | *Heliconius melpomene ssp. malleti* | Colombia | 1.8033 | -75.6553 | pending | | 3,675,883 | 69.34 | 2.18 | CS |
| CS3109 | CS003109 | *Heliconius melpomene ssp. malleti* | Colombia | 1.7108 | -75.7089 | pending | | 3,823,774 | 72.05 | 2.29 | CS |
| CS3366 | CS003366 | *Heliconius melpomene ssp. malleti* | Colombia | 1.7108 | -75.7089 | pending | | 3,791,556 | 71.50 | 2.21 | CS |
| CS3541 | CS003541 | *Heliconius melpomene ssp. malleti* | Colombia | 1.7506 | -75.6319 | pending | | 3,795,645 | 71.58 | 2.21 | CS |
| CS3543 | CS003543 | *Heliconius melpomene ssp. malleti* | Colombia | 1.7506 | -75.6319 | pending | | 3,816,480 | 71.98 | 2.20 | CS |
| CS3815 | CS003815 | *Heliconius melpomene ssp. malleti* | Colombia | 1.8033 | -75.6553 | pending | | 3,725,648 | 70.29 | 2.17 | CS |

| SequenceID | EarthCapeID | Taxon Name | Country | Latitude | Longitude | Accession | Publication | Called sites | Genotyped sites after filters (%) | Proportion of hetero-zygous genotypes (%) | Data Type (WGS…whole genome re-sequencing, CS…capture sequencing) |
|---|---|---|---|---|---|---|---|---|---|---|---|
| CS3730 | CS003730 | *Heliconius melpomene ssp. malleti* | Colombia | 1.8033 | -75.6553 | pending | | 3,780,839 | 71.51 | 1.92 | CS |
| CS470 | CS000470 | *Heliconius melpomene ssp. malleti* | Colombia | 1.8033 | -75.6553 | pending | | 3,627,455 | 68.43 | 2.18 | CS |
| CS3709 | CS003709 | *Heliconius melpomene ssp. malleti* | Colombia | 1.8033 | -75.6553 | pending | | 3,718,116 | 70.24 | 2.03 | CS |
| mal.CS1002 | CS001002 | *Heliconius melpomene ssp. malleti* | Colombia | 1.8033 | -75.6553 | SAMEA104585067 | Van Belleghem *et al.*[13] | 3,648,672 | 68.78 | 2.24 | WGS |
| mal.CS1011 | CS001011 | *Heliconius melpomene ssp. malleti* | Colombia | 1.8033 | -75.6553 | SAMEA104585068 | Van Belleghem *et al.*[13] | 3,425,148 | 64.49 | 2.36 | WGS |
| mal.CS1815 | CS001815 | *Heliconius melpomene ssp. malleti* | Colombia | 1.8033 | -75.6553 | SAMEA104585069 | Van Belleghem *et al.*[13] | 3,439,461 | 64.91 | 2.14 | WGS |
| mal.CS586 | CS000586 | *Heliconius melpomene ssp. malleti* | Colombia | 1.8033 | -75.6553 | SAMEA104585071 | Van Belleghem *et al.*[13] | 3,350,546 | 63.08 | 2.37 | WGS |
| mal.CS594 | CS000594 | *Heliconius melpomene ssp. malleti* | Colombia | 1.8033 | -75.6553 | SAMEA104585072 | Van Belleghem *et al.*[13] | 3,889,131 | 73.23 | 2.36 | WGS |
| mal.CS604 | CS000604 | *Heliconius melpomene ssp. malleti* | Colombia | 1.8033 | -75.6553 | SAMEA104585073 | Van Belleghem *et al.*[13] | 3,785,826 | 71.33 | 2.30 | WGS |
| mal.CS615 | CS000615 | *Heliconius melpomene ssp. malleti* | Colombia | 1.8033 | -75.6553 | SAMEA104585074 | Van Belleghem *et al.*[13] | 3,614,972 | 68.10 | 2.31 | WGS |
| mal.CS21 | CS002311 | *Heliconius melpomene ssp. malleti* | Colombia | 1.8136 | -75.6686 | SAMEA3723397 | Enciso-Romero *et al.*[6] | 3,918,174 | 73.70 | 2.46 | WGS |
| mal.CS22 | CS001286 | *Heliconius melpomene ssp. malleti* | Colombia | 1.6097 | -75.6669 | SAMEA3723398 | Enciso-Romero *et al.*[6] | 3,899,819 | 73.41 | 2.39 | WGS |
| mal.CS24 | CS001321 | *Heliconius melpomene ssp. malleti* | Colombia | 1.7506 | -75.6319 | SAMEA3723399 | Enciso-Romero *et al.*[6] | 3,845,940 | 72.38 | 2.41 | WGS |
| PS17381 | CAM017381 | *Heliconius melpomene ssp. malleti* | Ecuador | -1.0983 | -77.5839 | pending | | 3,695,732 | 69.70 | 2.20 | CS |
| PS17120 | CAM017120 | *Heliconius melpomene ssp. malleti* | Ecuador | -1.0916 | -77.7200 | pending | | 3,726,292 | 70.25 | 2.24 | CS |
| CAM017091 | CAM017091 | *Heliconius melpomene ssp. malleti* | Ecuador | -1.0614 | -77.6684 | pending | | 3,579,661 | 67.51 | 2.20 | CS |
| CAM017089 | CAM017089 | *Heliconius melpomene ssp. malleti* | Ecuador | -1.0614 | -77.6684 | pending | | 3,593,930 | 67.74 | 2.26 | CS |
| PS17073 | CAM017073 | *Heliconius melpomene ssp. malleti* | Ecuador | -1.1684 | -77.7811 | pending | | 3,719,819 | 70.14 | 2.23 | CS |
| CAM017070 | CAM017070 | *Heliconius melpomene ssp. malleti* | Ecuador | -1.1684 | -77.7811 | pending | | 3,603,436 | 67.92 | 2.26 | CS |
| CAM017069 | CAM017069 | *Heliconius melpomene ssp. malleti* | Ecuador | -1.1684 | -77.7811 | pending | | 3,611,832 | 68.07 | 2.27 | CS |
| CAM017064 | CAM017064 | *Heliconius melpomene ssp. malleti* | Ecuador | -1.1684 | -77.7811 | pending | | 3,579,409 | 67.50 | 2.21 | CS |
| PS17049 | CAM017049 | *Heliconius melpomene ssp. malleti* | Ecuador | -1.1684 | -77.7811 | pending | | 3,701,041 | 69.80 | 2.21 | CS |
| PS17030 | CAM017030 | *Heliconius melpomene ssp. malleti* | Ecuador | -1.0916 | -77.7200 | pending | | 3,706,478 | 69.91 | 2.19 | CS |
| CAM016956 | CAM016956 | *Heliconius melpomene ssp. malleti* | Ecuador | -1.1156 | -77.7783 | pending | | 3,557,390 | 67.11 | 2.17 | CS |
| CAM016955 | CAM016955 | *Heliconius melpomene ssp. malleti* | Ecuador | -1.1156 | -77.7783 | pending | | 3,624,964 | 68.34 | 2.24 | CS |
| CAM016954 | CAM016954 | *Heliconius melpomene ssp. malleti* | Ecuador | -1.1156 | -77.7783 | pending | | 3,586,392 | 67.65 | 2.19 | CS |
| PS16772 | CAM016772 | *Heliconius melpomene ssp. malleti* | Ecuador | -1.2519 | -77.8196 | pending | | 3,697,550 | 69.70 | 2.25 | CS |

| SequenceID | EarthCapeID | Taxon Name | Country | Latitude | Longitude | Accession | Publication | Called sites | Genotyped sites after filters (%) | Proportion of hetero-zygous genotypes (%) | Data Type (WGS...whole genome re-sequencing, CS...capture sequencing) |
|---|---|---|---|---|---|---|---|---|---|---|---|
| PS16766 | CAM016766 | *Heliconius melpomene ssp. malleti* | Ecuador | -1.2519 | -77.8196 | pending | | 3,730,622 | 70.35 | 2.21 | CS |
| CAM016611 | CAM016611 | *Heliconius melpomene ssp. malleti* | Ecuador | -1.1156 | -77.7783 | pending | | 3,635,301 | 68.54 | 2.23 | CS |
| CAM016610 | CAM016610 | *Heliconius melpomene ssp. malleti* | Ecuador | -1.1156 | -77.7783 | pending | | 3,606,744 | 68.01 | 2.22 | CS |
| CAM016609 | CAM016609 | *Heliconius melpomene ssp. malleti* | Ecuador | -1.1156 | -77.7783 | pending | | 3,592,637 | 67.74 | 2.22 | CS |
| CAM016606 | CAM016606 | *Heliconius melpomene ssp. malleti* | Ecuador | -1.1156 | -77.7783 | pending | | 3,575,042 | 67.39 | 2.26 | CS |
| CAM016599 | CAM016599 | *Heliconius melpomene ssp. malleti* | Ecuador | -1.1156 | -77.7783 | pending | | 3,613,904 | 68.14 | 2.22 | CS |
| CAM016549 | CAM016549 | *Heliconius melpomene ssp. malleti* | Ecuador | -1.0614 | -77.6684 | pending | | 3,623,370 | 68.30 | 2.25 | CS |
| CAM016547 | CAM016547 | *Heliconius melpomene ssp. malleti* | Ecuador | -1.0614 | -77.6684 | pending | | 3,645,314 | 68.69 | 2.28 | CS |
| CAM016546 | CAM016546 | *Heliconius melpomene ssp. malleti* | Ecuador | -1.0614 | -77.6684 | pending | | 3,615,315 | 68.10 | 2.32 | CS |
| CAM016544 | CAM016544 | *Heliconius melpomene ssp. malleti* | Ecuador | -1.0614 | -77.6684 | pending | | 3,593,545 | 67.75 | 2.24 | CS |
| CAM016541 | CAM016541 | *Heliconius melpomene ssp. malleti* | Ecuador | -1.0614 | -77.6684 | pending | | 3,604,024 | 67.93 | 2.26 | CS |
| PS16465 | CAM016465 | *Heliconius melpomene ssp. malleti* | Ecuador | -1.1878 | -77.8311 | pending | | 3,699,981 | 69.74 | 2.26 | CS |
| CAM016267 | CAM016267 | *Heliconius melpomene ssp. malleti* | Ecuador | -1.2510 | -77.6989 | pending | | 3,627,980 | 68.39 | 2.24 | CS |
| CAM016224 | CAM016224 | *Heliconius melpomene ssp. malleti* | Ecuador | -1.1156 | -77.7783 | pending | | 3,533,002 | 66.66 | 2.16 | CS |
| PS16144 | CAM016144 | *Heliconius melpomene ssp. malleti* | Ecuador | -1.4161 | -77.7290 | pending | | 3,711,697 | 70.00 | 2.20 | CS |
| CS1456 | CS001456 | *Heliconius melpomene ssp. melpomene* | Colombia | 4.1789 | -73.6494 | pending | | 4,016,701 | 75.81 | 2.13 | CS |
| CS1055 | CS001055 | *Heliconius melpomene ssp. melpomene* | Colombia | 4.0052 | -73.7775 | pending | | 4,016,080 | 75.83 | 2.09 | CS |
| CS1669 | CS001669 | *Heliconius melpomene ssp. melpomene* | Colombia | 4.2133 | -73.8028 | pending | | 3,985,424 | 75.67 | 1.55 | CS |
| CS1120 | CS001120 | *Heliconius melpomene ssp. melpomene* | Colombia | 4.0052 | -73.7775 | pending | | 4,028,453 | 76.02 | 2.14 | CS |
| CS1057 | CS001057 | *Heliconius melpomene ssp. melpomene* | Colombia | 4.2133 | -73.8028 | pending | | 4,003,690 | 75.55 | 2.15 | CS |
| CS1058 | CS001058 | *Heliconius melpomene ssp. melpomene* | Colombia | 4.2133 | -73.8028 | pending | | 3,978,857 | 75.20 | 2.00 | CS |
| CS3255 | CS003255 | *Heliconius melpomene ssp. melpomene* | Colombia | 4.2133 | -73.8028 | pending | | 4,015,755 | 75.89 | 2.00 | CS |
| CS3758 | CS003758 | *Heliconius melpomene ssp. melpomene* | Colombia | 4.2133 | -73.8028 | pending | | 4,005,712 | 75.68 | 2.03 | CS |
| CS1059 | CS001059 | *Heliconius melpomene ssp. melpomene* | Colombia | 4.2133 | -73.8028 | pending | | 4,007,649 | 75.75 | 1.99 | CS |
| CS1060 | CS001060 | *Heliconius melpomene ssp. melpomene* | Colombia | 4.2133 | -73.8028 | pending | | 3,985,438 | 75.49 | 1.77 | CS |
| CS1061 | CS001061 | *Heliconius melpomene ssp. melpomene* | Colombia | 4.2133 | -73.8028 | pending | | 4,012,207 | 75.76 | 2.08 | CS |
| CS3757 | CS003757 | *Heliconius melpomene ssp. melpomene* | Colombia | 4.2133 | -73.8028 | pending | | 4,000,033 | 75.56 | 2.05 | CS |

| SequenceID | EarthCapeID | Taxon Name | Country | Latitude | Longitude | Accession | Publication | Called sites | Genotyped sites after filters (%) | Proportion of hetero-zygous genotypes (%) | Data Type (WGS…whole genome re-sequencing, CS…capture sequencing) |
|---|---|---|---|---|---|---|---|---|---|---|---|
| CS1062 | CS001062 | *Heliconius melpomene ssp. melpomene* | Colombia | 4.2133 | -73.8028 | pending | | 4,016,089 | 76.03 | 1.83 | CS |
| melC.CS3 | CS000CM3 | *Heliconius melpomene ssp. melpomene* | Colombia | 4.2133 | -73.8028 | SAMEA3723393 | Martin *et al.*[16] | 4,161,140 | 78.65 | 1.99 | WGS |
| melC.CS6 | CS000CM6 | *Heliconius melpomene ssp. melpomene* | Colombia | 5.6169 | -72.3000 | SAMEA3723394 | Martin *et al.*[16] | 4,210,293 | 79.39 | 2.22 | WGS |
| melC.CS25 | CS000CM4 | *Heliconius melpomene ssp. melpomene* | Colombia | 4.2133 | -73.8028 | SAMEA3723400 | Martin *et al.*[16] | 4,102,616 | 77.43 | 2.13 | WGS |
| melC.CS26 | CS000CM5 | *Heliconius melpomene ssp. melpomene* | Colombia | 4.2133 | -73.8028 | SAMEA3723401 | Martin *et al.*[16] | 4,073,969 | 77.06 | 1.91 | WGS |
| melC.CS27 | CS00CM10 | *Heliconius melpomene ssp. melpomene* | Colombia | 5.6169 | -72.3000 | SAMEA3723402 | Martin *et al.*[16] | 4,198,256 | 79.16 | 2.22 | WGS |
| melG.CAM8215 | CAM008215 | *Heliconius melpomene ssp. melpomene* | French Guiana | 4.7890 | -52.4040 | SAMEA104585079 | | 1,228,623 | 23.43 | 1.12 | WGS |
| melG.CAM1349 | CAM001349 | *Heliconius melpomene ssp. melpomene* | French Guiana | 4.9632 | -52.4200 | SAMEA104585075 | Van Belleghem *et al.*[13] | 3,507,398 | 66.40 | 1.83 | WGS |
| melG.CAM1422 | CAM001422 | *Heliconius melpomene ssp. melpomene* | French Guiana | 4.9632 | -52.4200 | SAMEA104585076 | Van Belleghem *et al.*[13] | 3,217,943 | 60.93 | 1.82 | WGS |
| melG.CAM2035 | CAM002035 | *Heliconius melpomene ssp. melpomene* | French Guiana | 4.9632 | -52.4200 | SAMEA104585077 | Van Belleghem *et al.*[13] | 3,687,629 | 69.79 | 1.86 | WGS |
| melG.CAM8171 | CAM008171 | *Heliconius melpomene ssp. melpomene* | French Guiana | 4.9632 | -52.4200 | SAMEA104585078 | Van Belleghem *et al.*[13] | 3,466,269 | 65.62 | 1.84 | WGS |
| melG.CAM8216 | CAM008216 | *Heliconius melpomene ssp. melpomene* | French Guiana | 4.7890 | -52.4040 | SAMEA104585080 | Van Belleghem *et al.*[13] | 3,654,987 | 69.19 | 1.84 | WGS |
| melG.CAM8218 | CAM008218 | *Heliconius melpomene ssp. melpomene* | French Guiana | 4.7890 | -52.4040 | SAMEA104585081 | Van Belleghem *et al.*[13] | 3,464,056 | 65.56 | 1.86 | WGS |
| melG.CJ9316 | CAM009316 | *Heliconius melpomene ssp. melpomene* | French Guiana | 4.9632 | -52.4200 | SAMEA1919252 | Martin *et al.*[14] | 3,238,215 | 61.34 | 1.77 | WGS |
| melG.CJ9317 | CAM009317 | *Heliconius melpomene ssp. melpomene* | French Guiana | 4.9632 | -52.4200 | SAMEA1919267 | Martin *et al.*[14] | 3,798,559 | 71.89 | 1.86 | WGS |
| melG.CJ9315 | CAM009315 | *Heliconius melpomene ssp. melpomene* | French Guiana | 4.9632 | -52.4200 | SAMEA1919270 | Martin *et al.*[14] | 3,349,594 | 63.43 | 1.81 | WGS |
| melG.CJ13435 | CAM013435 | *Heliconius melpomene ssp. melpomene* | French Guiana | 4.9632 | -52.4200 | SAMEA1919276 | Martin *et al.*[14] | 3,827,918 | 72.44 | 1.87 | WGS |
| nan.CAM14673 | CAM014673 | *Heliconius melpomene ssp. melpomene* | Panamá | 8.6136 | -78.1398 | SAMEA104585082 | | 4,131,231 | 77.88 | 2.25 | WGS |
| CAM014662 | CAM014662 | *Heliconius melpomene ssp. melpomene* | Panamá | 8.6136 | -78.1398 | pending | | 4,102,605 | 77.51 | 2.03 | CS |
| CAM014654 | CAM014654 | *Heliconius melpomene ssp. melpomene* | Panamá | 8.6136 | -78.1398 | pending | | 4,110,765 | 77.69 | 2.00 | CS |
| CAM008956 | CAM008956 | *Heliconius melpomene ssp. melpomene* | Panamá | 7.6362 | -78.1897 | pending | | 4,112,268 | 77.60 | 2.15 | CS |
| CAM008954 | CAM008954 | *Heliconius melpomene ssp. melpomene* | Panamá | 7.6362 | -78.1897 | pending | | 4,109,799 | 77.65 | 2.02 | CS |
| CAM008887 | CAM008887 | *Heliconius melpomene ssp. melpomene* | Panamá | 7.7568 | -77.6841 | pending | | 4,116,541 | 77.77 | 2.04 | CS |
| melP.CJ18038 | CAM018038 | *Heliconius melpomene ssp. melpomene* | Panamá | 8.6136 | -78.1398 | SAMEA1919255 | Martin *et al.*[14] | 4,127,165 | 77.92 | 2.09 | WGS |
| melP.CJ18097 | CAM018097 | *Heliconius melpomene ssp. melpomene* | Panamá | 8.2797 | -77.8098 | SAMEA1919258 | Martin *et al.*[14] | 2,597,633 | 49.02 | 2.13 | WGS |
| CS2174 | CS002174 | *Heliconius melpomene ssp. melpomene* | Venezuela | 7.5878 | -72.1275 | pending | | 4,020,251 | 75.92 | 2.08 | CS |
| CAM013819 | CAM013819 | *Heliconius melpomene ssp. meriana* | French Guiana | 3.6883 | -54.0825 | SAMEA3670554 | Martin *et al.*[16] | 3,812,465 | 72.12 | 1.91 | CS |

| SequenceID | EarthCapeID | Taxon Name | Country | Latitude | Longitude | Accession | Publication | Called sites | Genotyped sites after filters (%) | Proportion of hetero-zygous genotypes (%) | Data Type (WGS…whole genome re-sequencing, CS…capture sequencing) |
|---|---|---|---|---|---|---|---|---|---|---|---|
| CAM013715 | CAM013715 | *Heliconius melpomene ssp. meriana* | French Guiana | 3.6883 | -54.0825 | SAMEA3670555 | Martin *et al.*[16] | 3,828,840 | 72.51 | 1.79 | CS |
| SHM21055 | CAM021055 | *Heliconius melpomene ssp. meriana* | Suriname | 5.0818 | -54.9791 | pending | | 3,874,135 | 73.31 | 1.88 | CS |
| RW12 | CAM019804 | *Heliconius melpomene ssp. 'meriana'* | England | 52.1905 | -1.7037 | pending | | 3,860,439 | 73.04 | 1.88 | CS |
| RW9 | CAM019803 | *Heliconius melpomene ssp. 'meriana'* | England | 52.1905 | -1.7037 | pending | | 3,814,479 | 72.17 | 1.89 | CS |
| RW5 | CAM019801 | *Heliconius melpomene ssp. 'meriana'* | England | 52.1905 | -1.7037 | pending | | 3,760,066 | 71.32 | 1.64 | CS |
| RW4 | CAM019800 | *Heliconius melpomene ssp. 'meriana'* | England | 52.1905 | -1.7037 | pending | | 3,884,713 | 73.88 | 1.38 | CS |
| nan.183-18 | CAM019828 | *Heliconius melpomene ssp. nanna* | Brazil | -3.8692 | -41.0192 | on rcs; pending | | 3,947,974 | 74.67 | 1.92 | WGS |
| nan.183-16 | CAM019827 | *Heliconius melpomene ssp. nanna* | Brazil | -3.8692 | -41.0192 | on rcs; pending | | 3,930,012 | 74.39 | 1.85 | WGS |
| nan.183-14 | CAM019826 | *Heliconius melpomene ssp. nanna* | Brazil | -3.8692 | -41.0192 | on rcs; pending | | 3,625,799 | 68.79 | 1.61 | WGS |
| nan.183-11 | CAM019825 | *Heliconius melpomene ssp. nanna* | Brazil | -3.8692 | -41.0192 | on rcs; pending | | 3,207,112 | 60.92 | 1.49 | WGS |
| nan.183-10 | CAM019824 | *Heliconius melpomene ssp. nanna* | Brazil | -3.8692 | -41.0192 | on rcs; pending | | 3,844,584 | 72.75 | 1.88 | WGS |
| GM209_26 | CAM019808 | *Heliconius melpomene ssp. nanna* | Brazil | -15.4201 | -39.4964 | pending | | 3,860,711 | 74.03 | 0.56 | CS |
| GM183_19 | CAM019807 | *Heliconius melpomene ssp. nanna* | Brazil | -3.8692 | -41.0192 | pending | | 3,707,040 | 70.20 | 1.80 | CS |
| GM183_15 | CAM019806 | *Heliconius melpomene ssp. nanna* | Brazil | -3.8692 | -41.0192 | pending | | 3,885,848 | 73.57 | 1.83 | CS |
| MK000014 | MK000014 | *Heliconius melpomene ssp. nanna* | Brazil | -19.0984 | -40.1862 | SAMN04407968 | Zhang *et al.*[17] | 3,301,446 | 63.27 | 0.63 | WGS |
| MK000062 | MK000062 | *Heliconius melpomene ssp. nanna* | Brazil | -19.0984 | -40.1862 | SAMN04407969; pending | Zhang *et al.*[17] | 3,845,064 | 73.68 | 0.63 | CS |
| MK000063 | MK000063 | *Heliconius melpomene ssp. nanna* | Brazil | -19.0984 | -40.1862 | SAMN04407970 | Zhang *et al.*[17] | 3,710,880 | 71.16 | 0.56 | WGS |
| MK000064 | MK000064 | *Heliconius melpomene ssp. nanna* | Brazil | -19.0984 | -40.1862 | SAMN04407971 | Zhang *et al.*[17] | 3,984,097 | 76.34 | 0.64 | WGS |
| PS17613 | CAM017613 | *Heliconius melpomene ssp. plesseni* | Ecuador | -1.4224 | -78.1729 | pending | | 3,868,444 | 73.25 | 1.81 | CS |
| PS17373 | CAM017373 | *Heliconius melpomene ssp. plesseni* | Ecuador | -1.4600 | -78.0728 | pending | | 3,897,348 | 73.57 | 2.11 | CS |
| PS17363 | CAM017363 | *Heliconius melpomene ssp. plesseni* | Ecuador | -1.4600 | -78.0728 | pending | | 3,888,631 | 73.47 | 2.02 | CS |
| PS17357 | CAM017357 | *Heliconius melpomene ssp. plesseni* | Ecuador | -1.4600 | -78.0728 | pending | | 3,896,339 | 73.61 | 2.04 | CS |
| PS16913 | CAM016913 | *Heliconius melpomene ssp. plesseni* | Ecuador | -1.4371 | -78.1229 | pending | | 3,839,335 | 72.55 | 2.01 | CS |
| PS16910 | CAM016910 | *Heliconius melpomene ssp. plesseni* | Ecuador | -1.4371 | -78.1229 | pending | | 3,842,150 | 72.59 | 2.03 | CS |
| PS16691 | CAM016691 | *Heliconius melpomene ssp. plesseni* | Ecuador | -1.3980 | -78.1781 | pending | | 3,862,444 | 73.00 | 2.00 | CS |
| PS16004 | CAM016004 | *Heliconius melpomene ssp. plesseni* | Ecuador | -1.4224 | -78.1729 | pending | | 3,859,802 | 72.95 | 2.00 | CS |
| PS16003 | CAM016003 | *Heliconius melpomene ssp. plesseni* | Ecuador | -1.4224 | -78.1729 | pending | | 3,879,223 | 73.31 | 2.00 | CS |

| SequenceID | EarthCapeID | Taxon Name | Country | Latitude | Longitude | Accession | Publication | Called sites | Genotyped sites after filters (%) | Proportion of hetero-zygous genotypes (%) | Data Type (WGS…whole genome re-sequencing, CS…capture sequencing) |
|---|---|---|---|---|---|---|---|---|---|---|---|
| PS16001 | CAM016001 | *Heliconius melpomene ssp. plesseni* | Ecuador | -1.4224 | -78.1729 | pending | | 3,902,878 | 73.92 | 1.79 | CS |
| ple.CJ9156 | CAM009156 | *Heliconius melpomene ssp. plesseni* | Ecuador | -1.3980 | -78.1781 | SAMEA3670556 | Martin *et al.*[16] | 2,789,820 | 52.74 | 1.96 | WGS |
| ple.CJ16293 | CAM016293 | *Heliconius melpomene ssp. plesseni* | Ecuador | -1.4600 | -78.0728 | SAMEA3670557 | Martin *et al.*[16] | 2,610,868 | 49.33 | 2.02 | WGS |
| ros.CAM1841 | CAM001841 | *Heliconius melpomene ssp. rosina* | Panamá | 9.0760 | -79.6590 | SAMEA104585083 | Van Belleghem *et al.*[13] | 3,937,058 | 74.32 | 2.11 | WGS |
| ros.CAM1880 | CAM001880 | *Heliconius melpomene ssp. rosina* | Panamá | 9.0760 | -79.6590 | SAMEA104585084 | Van Belleghem *et al.*[13] | 3,970,697 | 74.99 | 2.07 | WGS |
| ros.CAM2045 | CAM002045 | *Heliconius melpomene ssp. rosina* | Panamá | 9.1103 | -79.6907 | SAMEA104585085 | Van Belleghem *et al.*[13] | 4,105,191 | 77.53 | 2.06 | WGS |
| ros.CAM2059 | CAM002059 | *Heliconius melpomene ssp. rosina* | Panamá | 9.1103 | -79.6907 | SAMEA104585086 | Van Belleghem *et al.*[13] | 4,154,613 | 78.51 | 2.01 | WGS |
| ros.CAM2519 | CAM002519 | *Heliconius melpomene ssp. rosina* | Panamá | 9.0109 | -79.5477 | SAMEA104585087 | Van Belleghem *et al.*[13] | 4,157,116 | 78.45 | 2.14 | WGS |
| ros.CAM2552 | CAM002552 | *Heliconius melpomene ssp. rosina* | Panamá | 9.0109 | -79.5477 | SAMEA104585088 | Van Belleghem *et al.*[13] | 3,520,446 | 66.52 | 2.02 | WGS |
| ros.CJ2071 | CAM002071 | *Heliconius melpomene ssp. rosina* | Panamá | 9.1103 | -79.6907 | SAMEA1919257 | Martin *et al.*[14] | 4,089,428 | 77.27 | 2.02 | WGS |
| ros.CJ533 | CAM000533 | *Heliconius melpomene ssp. rosina* | Panamá | 9.1103 | -79.6907 | SAMEA1919260 | Martin *et al.*[14] | 3,442,199 | 65.09 | 1.94 | WGS |
| ros.CJ531 | CAM000531 | *Heliconius melpomene ssp. rosina* | Panamá | 9.1103 | -79.6907 | SAMEA1919271 | Martin *et al.*[14] | 3,556,153 | 67.21 | 1.99 | WGS |
| ros.CJ546 | CAM000546 | *Heliconius melpomene ssp. rosina* | Panamá | 9.1253 | -79.6932 | SAMEA1919279 | Martin *et al.*[14] | 3,445,216 | 65.14 | 1.96 | WGS |
| CS32533 | CS032533 | *Heliconius melpomene ssp. vicina* | Colombia | -4.1003 | -70.0417 | pending | | 3,927,898 | 74.07 | 2.21 | CS |
| CS31744 | CS031744 | *Heliconius melpomene ssp. vicina* | Colombia | -4.1334 | -69.9415 | pending | | 3,957,710 | 74.58 | 2.29 | CS |
| CS31755 | CS031755 | *Heliconius melpomene ssp. vicina* | Colombia | -4.1334 | -69.9415 | pending | | 3,932,967 | 74.13 | 2.26 | CS |
| vul.CS3603 | CS003603 | *Heliconius melpomene ssp. vulcanus* | Colombia | 3.5175 | -76.7572 | SAMEA104585091 | Van Belleghem *et al.*[13] | 4,205,795 | 79.50 | 1.98 | WGS |
| vul.CS3605 | CS003605 | *Heliconius melpomene ssp. vulcanus* | Colombia | 3.5175 | -76.7572 | SAMEA104585092 | Van Belleghem *et al.*[13] | 4,193,786 | 79.25 | 2.01 | WGS |
| vul.CS3606 | CS003606 | *Heliconius melpomene ssp. vulcanus* | Colombia | 3.5175 | -76.7572 | SAMEA104585093 | Van Belleghem *et al.*[13] | 4,162,523 | 78.55 | 2.14 | WGS |
| vul.CS3612 | CS003612 | *Heliconius melpomene ssp. vulcanus* | Colombia | 3.5175 | -76.7572 | SAMEA104585094 | Van Belleghem *et al.*[13] | 4,201,547 | 79.77 | 1.54 | WGS |
| vul.CS3614 | CS003614 | *Heliconius melpomene ssp. vulcanus* | Colombia | 3.5175 | -76.7572 | SAMEA104585095 | Van Belleghem *et al.*[13] | 3,861,940 | 72.92 | 2.09 | WGS |
| vul.CS3615 | CS003615 | *Heliconius melpomene ssp. vulcanus* | Colombia | 3.5175 | -76.7572 | SAMEA104585096 | Van Belleghem *et al.*[13] | 3,717,208 | 70.41 | 1.78 | WGS |
| vul.CS3617 | CS003617 | *Heliconius melpomene ssp. vulcanus* | Colombia | 3.5175 | -76.7572 | SAMEA104585097 | Van Belleghem *et al.*[13] | 3,805,245 | 71.87 | 2.06 | WGS |
| vul.CS3618 | CS003618 | *Heliconius melpomene ssp. vulcanus* | Colombia | 3.5175 | -76.7572 | SAMEA104585098 | Van Belleghem *et al.*[13] | 3,820,701 | 72.46 | 1.66 | WGS |
| vul.CS3621 | CS003621 | *Heliconius melpomene ssp. vulcanus* | Colombia | 3.5175 | -76.7572 | SAMEA104585099 | Van Belleghem *et al.*[13] | 3,742,085 | 70.61 | 2.15 | WGS |
| vul.CS10 | CS000710 | *Heliconius melpomene ssp. vulcanus* | Colombia | 3.9000 | -76.6325 | SAMEA3723391 | Enciso-Romero *et al.*[6]; Martin *et al.*[16] | 4,255,149 | 80.43 | 1.98 | WGS |

| SequenceID | EarthCapeID | Taxon Name | Country | Latitude | Longitude | Accession | Publication | Called sites | Genotyped sites after filters (%) | Proportion of hetero-zygous genotypes (%) | Data Type (WGS…whole genome re-sequencing, CS…capture sequencing) |
|---|---|---|---|---|---|---|---|---|---|---|---|
| vul.CJ14632 | CAM014632 | *Heliconius melpomene ssp. vulcanus* | Panamá | 8.6136 | -78.1398 | SAMEA3670560 | Wallbank *et al.*[8] | 2,402,825 | 45.39 | 2.03 | WGS |
| MJ12-3636 | MJ12-3636 | *Heliconius melpomene ssp. xenoclea* | Peru | -11.0338 | -75.4091 | pending | | 3,516,391 | 66.53 | 1.89 | CS |
| MJ12-3653 | MJ12-3653 | *Heliconius melpomene ssp. xenoclea* | Peru | -11.0338 | -75.4091 | pending | | 3,659,935 | 69.21 | 1.93 | CS |
| MJ12-3605 | MJ12-3605 | *Heliconius melpomene ssp. xenoclea* | Peru | -11.1745 | -75.4035 | pending | | 3,608,285 | 68.26 | 1.90 | CS |
| MJ12-3648 | MJ12-3648 | *Heliconius melpomene ssp. xenoclea* | Peru | -11.0338 | -75.4091 | pending | | 3,603,354 | 68.16 | 1.92 | CS |
| MJ12-3647 | MJ12-3647 | *Heliconius melpomene ssp. xenoclea* | Peru | -11.0446 | -75.4133 | pending | | 3,538,622 | 66.93 | 1.92 | CS |
| MJ12-3763 | MJ12-3763 | *Heliconius melpomene ssp. xenoclea* | Peru | -11.0338 | -75.4091 | pending | | 3,584,335 | 67.83 | 1.86 | CS |
| MJ12-3651 | MJ12-3651 | *Heliconius melpomene ssp. xenoclea* | Peru | -11.0338 | -75.4091 | pending | | 3,399,714 | 64.38 | 1.80 | CS |
| MJ12-3608 | MJ12-3608 | *Heliconius melpomene ssp. xenoclea* | Peru | -11.1745 | -75.4035 | pending | | 3,379,925 | 64.01 | 1.80 | CS |
| MJ12-3606 | MJ12-3606 | *Heliconius melpomene ssp. xenoclea* | Peru | -11.1745 | -75.4035 | pending | | 3,527,682 | 66.74 | 1.89 | CS |
| MJ12-3638 | MJ12-3638 | *Heliconius melpomene ssp. xenoclea* | Peru | -11.0364 | -75.4080 | pending | | 3,593,153 | 67.99 | 1.88 | CS |
| nu_sil.MJ09-4125 | MJ09-4125 | *Heliconius numata ssp. numata* | French Guiana | 4.0833 | -52.6753 | SAMEA3888884 | Nadeau *et al.*[7] | 3,429,209 | 64.05 | 3.15 | WGS |
| nu_sil.MJ09-4184 | MJ09-4184 | *Heliconius numata ssp. silvana* | French Guiana | 4.0833 | -52.6753 | SAMEA3888889 | Nadeau *et al.*[7] | 3,514,463 | 65.52 | 3.33 | WGS |
| nu_sil.MJ05-1240 | JM-05-1240 | *Heliconius numata ssp. silvana* | Peru | -6.4782 | -76.3258 | SAMEA3888886 | Nadeau *et al.*[7] | 2,133,564 | 39.94 | 2.93 | WGS |
| nu_sil.MJ05-1271 | JM-05-1271 | *Heliconius numata ssp. silvana* | Peru | -6.4782 | -76.3258 | SAMEA3888887 | Nadeau *et al.*[7] | 3,383,291 | 63.11 | 3.28 | WGS |
| nu_sil.MJ05-124 | MJ05-124 | *Heliconius numata ssp. silvana* | Peru | -6.4782 | -76.3258 | SAMEA3888888 | Nadeau *et al.*[7] | 3,336,988 | 62.22 | 3.32 | WGS |
| MK000516 | MK000516 | *Heliconius pachinus* | Costa Rica | 9.8500 | -84.3167 | SAMN02384468 | Kronforst *et al.*[18] | 3,399,813 | 64.39 | 1.78 | WGS |
| CAM008062 | CAM008062 | *Heliconius pachinus* | Panamá | 8.8388 | -82.7148 | pending | | 3,803,511 | 72.07 | 1.75 | CS |
| CAM008036 | CAM008036 | *Heliconius pachinus* | Panamá | 8.8388 | -82.7148 | pending | | 3,813,469 | 72.02 | 2.07 | CS |
| CAM008034 | CAM008034 | *Heliconius pachinus* | Panamá | 8.8388 | -82.7148 | pending | | 3,810,903 | 72.01 | 2.02 | CS |
| CAM008033 | CAM008033 | *Heliconius pachinus* | Panamá | 8.8388 | -82.7148 | pending | | 3,789,692 | 71.80 | 1.75 | CS |
| CAM008032 | CAM008032 | *Heliconius pachinus* | Panamá | 8.8388 | -82.7148 | pending | | 3,812,716 | 72.26 | 1.73 | CS |
| CAM008023 | CAM008023 | *Heliconius pachinus* | Panamá | 8.8388 | -82.7148 | pending | | 3,782,656 | 71.58 | 1.87 | CS |
| CAM008022 | CAM008022 | *Heliconius pachinus* | Panamá | 8.8388 | -82.7148 | pending | | 3,824,865 | 72.43 | 1.81 | CS |
| CAM008019 | CAM008019 | *Heliconius pachinus* | Panamá | 8.8388 | -82.7148 | pending | | 3,800,262 | 71.62 | 2.28 | CS |
| CAM008035 | CAM008035 | *Heliconius pachinus* | Panamá | 8.8388 | -82.7148 | SAMEA3670565; pending | Wallbank *et al.*[8] | 3,804,700 | 72.10 | 1.73 | CS |
| ser.JM202 | JM-09-202 | *Heliconius pardalinus ssp. sergestus* | Peru | -6.4667 | -76.3347 | SAMEA1919268 | Martin *et al.*[14] | 3,723,595 | 71.15 | 0.92 | WGS |
| par.JM371 | JM-09-371 | *Heliconius pardalinus ssp. ssp.nov.P* | Peru | -8.3422 | -74.5922 | SAMEA1919253 | Martin *et al.*[14] | 3,672,441 | 68.65 | 3.06 | WGS |
| CS2394 | CS002394 | *Heliconius timareta ssp. florencia* | Colombia | 1.7108 | -75.7089 | pending | | 3,794,165 | 71.82 | 1.84 | CS |
| CS2343 | CS002343 | *Heliconius timareta ssp. florencia* | Colombia | 1.8136 | -75.6686 | pending | | 3,878,536 | 73.42 | 1.83 | CS |
| CS2333 | CS002333 | *Heliconius timareta ssp. florencia* | Colombia | 1.7108 | -75.7089 | pending | | 3,835,744 | 72.64 | 1.79 | CS |
| CS2327 | CS002327 | *Heliconius timareta ssp. florencia* | Colombia | 1.7108 | -75.7089 | pending | | 3,849,177 | 72.91 | 1.78 | CS |
| CS419 | CS000419 | *Heliconius timareta ssp. florencia* | Colombia | 1.8033 | -75.6553 | pending | | 3,736,703 | 70.74 | 1.83 | CS |

| SequenceID | EarthCapeID | Taxon Name | Country | Latitude | Longitude | Accession | Publication | Called sites | Genotyped sites after filters (%) | Proportion of hetero-zygous genotypes (%) | Data Type (WGS…whole genome re-sequencing, CS…capture sequencing) |
|---|---|---|---|---|---|---|---|---|---|---|---|
| CS399 | CS000399 | *Heliconius timareta ssp. florencia* | Colombia | 1.8033 | -75.6553 | pending | | 3,698,884 | 70.15 | 1.66 | CS |
| CS2349 | CS002349 | *Heliconius timareta ssp. florencia* | Colombia | 1.7108 | -75.7089 | pending | | 3,836,299 | 72.64 | 1.81 | CS |
| CS1009 | CS001009 | *Heliconius timareta ssp. florencia* | Colombia | 1.8033 | -75.6553 | pending | | 3,847,891 | 72.85 | 1.82 | CS |
| CS473 | CS000473 | *Heliconius timareta ssp. florencia* | Colombia | 1.8033 | -75.6553 | pending | | 3,828,399 | 72.39 | 1.95 | CS |
| CS2354 | CS002354 | *Heliconius timareta ssp. florencia* | Colombia | 1.7108 | -75.7089 | pending | | 3,871,665 | 73.42 | 1.67 | CS |
| flo.CS12 | CS002395 | *Heliconius timareta ssp. florencia* | Colombia | 1.7097 | -75.6976 | SAMEA104585100 | Van Belleghem *et al.*[13] | 3,897,565 | 73.76 | 1.87 | WGS |
| flo.CS13 | CS002402 | *Heliconius timareta ssp. florencia* | Colombia | 1.7097 | -75.6976 | SAMEA104585101 | Van Belleghem *et al.*[13] | 3,872,164 | 73.23 | 1.93 | WGS |
| flo.CS14 | CS002403 | *Heliconius timareta ssp. florencia* | Colombia | 1.7097 | -75.6976 | SAMEA104585102 | Van Belleghem *et al.*[13] | 3,910,873 | 73.99 | 1.89 | WGS |
| flo.CS15 | CS002406 | *Heliconius timareta ssp. florencia* | Colombia | 1.7097 | -75.6976 | SAMEA104585103 | Van Belleghem *et al.*[13] | 3,904,353 | 73.79 | 1.99 | WGS |
| flo.CS2337 | CS002337 | *Heliconius timareta ssp. florencia* | Colombia | 1.7108 | -75.7089 | SAMEA104585104 | Van Belleghem *et al.*[13] | 3,681,700 | 69.62 | 1.94 | WGS |
| flo.CS2338 | CS002338 | *Heliconius timareta ssp. florencia* | Colombia | 1.7108 | -75.7089 | SAMEA104585105 | Van Belleghem *et al.*[13] | 3,723,102 | 70.48 | 1.83 | WGS |
| flo.CS2341 | CS002341 | *Heliconius timareta ssp. florencia* | Colombia | 1.8136 | -75.6686 | SAMEA104585106 | Van Belleghem *et al.*[13] | 3,579,194 | 67.68 | 1.94 | WGS |
| flo.CS2350 | CS002350 | *Heliconius timareta ssp. florencia* | Colombia | 1.7108 | -75.7089 | SAMEA104585107 | Van Belleghem *et al.*[13] | 3,741,211 | 70.75 | 1.93 | WGS |
| flo.CS2358 | CS002358 | *Heliconius timareta ssp. florencia* | Colombia | 1.7108 | -75.7089 | SAMEA104585108 | Van Belleghem *et al.*[13] | 3,829,666 | 72.45 | 1.89 | WGS |
| flo.CS2359 | CS002359 | *Heliconius timareta ssp. florencia* | Colombia | 1.7108 | -75.7089 | SAMEA104585109 | Van Belleghem *et al.*[13] | 3,693,632 | 69.88 | 1.90 | WGS |
| CS3009 | CS003009 | *Heliconius timareta ssp. linaresi* | Colombia | 2.6844 | -74.8881 | pending | | 3,820,489 | 72.41 | 1.71 | CS |
| CS2429 | CS002429 | *Heliconius timareta ssp. linaresi* | Colombia | 2.6090 | -74.7776 | pending | | 3,797,759 | 71.93 | 1.78 | CS |
| CS2436 | CS002436 | *Heliconius timareta ssp. linaresi* | Colombia | 2.6090 | -74.7776 | pending | | 3,822,561 | 72.41 | 1.76 | CS |
| CS3014 | CS003014 | *Heliconius timareta ssp. linaresi* | Colombia | 2.6844 | -74.8881 | pending | | 3,781,757 | 71.51 | 1.95 | CS |
| CS3012 | CS003012 | *Heliconius timareta ssp. linaresi* | Colombia | 2.6844 | -74.8881 | pending | | 3,800,749 | 71.99 | 1.78 | CS |
| CS3760 | CS003760 | *Heliconius timareta ssp. linaresi* | Colombia | 2.6844 | -74.8881 | pending | | 3,793,257 | 72.04 | 1.51 | CS |
| CS3050 | CS003050 | *Heliconius timareta ssp. linaresi* | Colombia | 2.6844 | -74.8881 | pending | | 3,814,986 | 72.15 | 1.93 | CS |
| CS3032 | CS003032 | *Heliconius timareta ssp. linaresi* | Colombia | 2.6844 | -74.8881 | pending | | 3,821,017 | 72.40 | 1.74 | CS |
| CS2422 | CS002422 | *Heliconius timareta ssp. linaresi* | Colombia | 2.6090 | -74.7776 | pending | | 3,819,912 | 72.39 | 1.74 | CS |
| CS3013 | CS003013 | *Heliconius timareta ssp. linaresi* | Colombia | 2.6844 | -74.8881 | pending | | 3,827,502 | 72.40 | 1.91 | CS |
| CS3046 | CS003046 | *Heliconius timareta ssp. linaresi* | Colombia | 2.6844 | -74.8881 | pending | | 3,741,134 | 70.90 | 1.72 | CS |
| CS3762 | CS003762 | *Heliconius timareta ssp. linaresi* | Colombia | 2.6844 | -74.8881 | pending | | 3,803,044 | 72.08 | 1.71 | CS |
| CS3367 | CS003367 | *Heliconius timareta ssp. linaresi* | Colombia | 2.6844 | -74.8881 | pending | | 3,804,131 | 72.04 | 1.79 | CS |
| CS3763 | CS003763 | *Heliconius timareta ssp. linaresi* | Colombia | 2.6844 | -74.8881 | pending | | 3,822,941 | 72.35 | 1.86 | CS |
| CS3372 | CS003372 | *Heliconius timareta ssp. linaresi* | Colombia | 2.6844 | -74.8881 | pending | | 3,805,156 | 72.26 | 1.53 | CS |
| CS3051 | CS003051 | *Heliconius timareta ssp. linaresi* | Colombia | 2.6844 | -74.8881 | pending | | 3,820,696 | 72.31 | 1.86 | CS |
| CS3761 | CS003761 | *Heliconius timareta ssp. linaresi* | Colombia | 2.6844 | -74.8881 | pending | | 3,823,361 | 72.53 | 1.63 | CS |
| CS3086 | CS003086 | *Heliconius timareta ssp. linaresi* | Colombia | 2.6844 | -74.8881 | pending | | 3,821,604 | 72.35 | 1.82 | CS |
| CS2435 | CS002435 | *Heliconius timareta ssp. linaresi* | Colombia | 2.6090 | -74.7776 | SAMEA1094485; pending | Nadeau *et al.*[19]; SureSelect | 3,833,747 | 72.55 | 1.87 | CS |
| PS16870 | CAM016870 | *Heliconius timareta ssp. nov. ECU* | Ecuador | -1.3712 | -77.8574 | pending | | 3,750,493 | 71.04 | 1.78 | CS |

| SequenceID | EarthCapeID | Taxon Name | Country | Latitude | Longitude | Accession | Publication | Called sites | Genotyped sites after filters (%) | Proportion of hetero-zygous genotypes (%) | Data Type (WGS…whole genome re-sequencing, CS…capture sequencing) |
|---|---|---|---|---|---|---|---|---|---|---|---|
| PS16857 | CAM016857 | *Heliconius timareta ssp. nov. ECU* | Ecuador | -1.4021 | -77.7974 | pending | | 3,764,862 | 71.30 | 1.80 | CS |
| CAM016605 | CAM016605 | *Heliconius timareta ssp. nov. ECU* | Ecuador | -1.1156 | -77.7783 | pending | | 3,685,825 | 69.87 | 1.71 | CS |
| CAM016603 | CAM016603 | *Heliconius timareta ssp. nov. ECU* | Ecuador | -1.1156 | -77.7783 | pending | | 3,611,835 | 68.46 | 1.71 | CS |
| CAM016600 | CAM016600 | *Heliconius timareta ssp. nov. ECU* | Ecuador | -1.1156 | -77.7783 | pending | | 3,668,367 | 69.47 | 1.80 | CS |
| CAM016593 | CAM016593 | *Heliconius timareta ssp. nov. ECU* | Ecuador | -1.1156 | -77.7783 | pending | | 3,662,851 | 69.38 | 1.77 | CS |
| CAM016502 | CAM016502 | *Heliconius timareta ssp. nov. ECU* | Ecuador | -1.2908 | -77.8419 | pending | | 3,675,153 | 69.63 | 1.76 | CS |
| CAM016501 | CAM016501 | *Heliconius timareta ssp. nov. ECU* | Ecuador | -1.2908 | -77.8419 | pending | | 3,692,687 | 69.93 | 1.80 | CS |
| CAM016443 | CAM016443 | *Heliconius timareta ssp. nov. ECU* | Ecuador | -1.1878 | -77.8311 | pending | | 3,679,600 | 69.67 | 1.82 | CS |
| CAM016442 | CAM016442 | *Heliconius timareta ssp. nov. ECU* | Ecuador | -1.1878 | -77.8311 | pending | | 3,670,424 | 69.55 | 1.74 | CS |
| CAM016304 | CAM016304 | *Heliconius timareta ssp. nov. ECU* | Ecuador | -1.3333 | -77.9341 | pending | | 3,724,010 | 70.50 | 1.83 | CS |
| PS16269 | CAM016269 | *Heliconius timareta ssp. nov. ECU* | Ecuador | -1.2510 | -77.6989 | pending | | 3,764,038 | 71.37 | 1.68 | CS |
| CAM016260 | CAM016260 | *Heliconius timareta ssp. nov. ECU* | Ecuador | -1.2510 | -77.6989 | pending | | 3,695,564 | 70.04 | 1.72 | CS |
| CAM016257 | CAM016257 | *Heliconius timareta ssp. nov. ECU* | Ecuador | -1.2510 | -77.6989 | pending | | 3,694,177 | 70.01 | 1.72 | CS |
| CAM016255 | CAM016255 | *Heliconius timareta ssp. nov. ECU* | Ecuador | -1.2510 | -77.6989 | pending | | 3,688,248 | 69.89 | 1.74 | CS |
| PS16252 | CAM016252 | *Heliconius timareta ssp. nov. ECU* | Ecuador | -1.8176 | -77.9601 | pending | | 3,754,683 | 71.13 | 1.76 | CS |
| CAM016218 | CAM016218 | *Heliconius timareta ssp. nov. ECU* | Ecuador | -1.1156 | -77.7783 | pending | | 3,654,949 | 69.19 | 1.83 | CS |
| CAM016214 | CAM016214 | *Heliconius timareta ssp. nov. ECU* | Ecuador | -1.1156 | -77.7783 | pending | | 3,682,926 | 69.78 | 1.75 | CS |
| PS16201 | CAM016201 | *Heliconius timareta ssp. nov. ECU* | Ecuador | -1.1156 | -77.7783 | pending | | 3,756,272 | 71.14 | 1.79 | CS |
| PS16200 | CAM016200 | *Heliconius timareta ssp. nov. ECU* | Ecuador | -1.1156 | -77.7783 | pending | | 3,746,590 | 70.99 | 1.74 | CS |
| PS16173 | CAM016173 | *Heliconius timareta ssp. nov. ECU* | Ecuador | -1.2408 | -77.9609 | pending | | 3,755,655 | 71.09 | 1.85 | CS |
| CAM016037 | CAM016037 | *Heliconius timareta ssp. nov. ECU* | Ecuador | -1.2908 | -77.8419 | pending | | 3,696,783 | 69.94 | 1.89 | CS |
| PS16011 | CAM016011 | *Heliconius timareta ssp. nov. ECU* | Ecuador | -1.3333 | -77.9341 | pending | | 3,765,599 | 71.30 | 1.82 | CS |
| PS16425 | CAM016425 | *Heliconius timareta ssp. nov. ECU* | Ecuador | -1.1878 | -77.8311 | SAMEA2240078; pending | Nadeau *et al.*[20]; RAD | 3,754,500 | 71.13 | 1.75 | CS |
| PS17380 | CAM017380 | *Heliconius timareta ssp. nov. ECU* | Ecuador | -1.0983 | -77.5839 | SAMEA2240096; pending | Nadeau *et al.*[20]; RAD | 3,759,205 | 71.20 | 1.78 | CS |
| CS31766 | CS031766 | *Heliconius timareta ssp. nov. 'vicina'* | Colombia | -4.1334 | -69.9415 | pending | | 3,881,511 | 73.68 | 1.57 | CS |
| CS31733 | CS031733 | *Heliconius timareta ssp. nov. 'vicina'* | Colombia | -3.7701 | -70.3398 | pending | | 3,917,016 | 74.32 | 1.61 | CS |
| CS31809 | CS031809 | *Heliconius timareta ssp. nov. 'vicina'* | Colombia | -3.7701 | -70.3398 | pending | | 3,876,918 | 73.61 | 1.54 | CS |
| CS31711 | CS031711 | *Heliconius timareta ssp. nov. 'vicina'* | Colombia | -3.7701 | -70.3398 | pending | | 3,911,288 | 74.17 | 1.66 | CS |
| CS32537 | CS032537 | *Heliconius timareta ssp. nov. 'vicina'* | Colombia | -3.8833 | -70.1881 | pending | | 3,898,267 | 73.99 | 1.57 | CS |
| CS32535 | CS032535 | *Heliconius timareta ssp. nov. 'vicina'* | Colombia | -4.0406 | -70.0997 | pending | | 3,864,120 | 73.25 | 1.70 | CS |
| MJ12-3388 | MJ12-3388 | *Heliconius timareta ssp. thelxinoe* | Peru | -6.4515 | -76.2977 | pending | | 3,681,338 | 69.66 | 1.88 | CS |
| MJ12-3310 | MJ12-3310 | *Heliconius timareta ssp. thelxinoe* | Peru | -6.4519 | -76.2985 | pending | | 3,462,285 | 65.77 | 1.50 | CS |
| MJ12-3232 | MJ12-3232 | *Heliconius timareta ssp. thelxinoe* | Peru | -6.4515 | -76.2977 | pending | | 3,664,260 | 69.42 | 1.76 | CS |
| MJ12-3412 | MJ12-3412 | *Heliconius timareta ssp. thelxinoe* | Peru | -6.4537 | -76.2981 | pending | | 3,746,879 | 70.93 | 1.84 | CS |
| MJ12-3160 | MJ12-3160 | *Heliconius timareta ssp. thelxinoe* | Peru | -6.4519 | -76.2985 | pending | | 3,692,631 | 69.94 | 1.78 | CS |
| MJ12-3212 | MJ12-3212 | *Heliconius timareta ssp. thelxinoe* | Peru | -6.4547 | -76.2994 | pending | | 3,657,709 | 69.37 | 1.65 | CS |
| MJ12-3402 | MJ12-3402 | *Heliconius timareta ssp. thelxinoe* | Peru | -6.4528 | -76.2862 | pending | | 3,684,144 | 69.69 | 1.91 | CS |
| MJ12-3222 | MJ12-3222 | *Heliconius timareta ssp. thelxinoe* | Peru | -6.4519 | -76.2985 | pending | | 3,690,826 | 69.90 | 1.79 | CS |
| MJ12-3138 | MJ12-3138 | *Heliconius timareta ssp. thelxinoe* | Peru | -6.4547 | -76.2994 | pending | | 3,682,422 | 69.97 | 1.47 | CS |

| SequenceID | EarthCapeID | Taxon Name | Country | Latitude | Longitude | Accession | Publication | Called sites | Genotyped sites after filters (%) | Proportion of hetero-zygous genotypes (%) | Data Type (WGS…whole genome re-sequencing, CS…capture sequencing) |
|---|---|---|---|---|---|---|---|---|---|---|---|
| MJ12-3398 | MJ12-3398 | *Heliconius timareta ssp. thelxinoe* | Peru | -6.4515 | -76.2977 | pending | | 3,678,752 | 69.61 | 1.87 | CS |
| MJ12-3220 | MJ12-3220 | *Heliconius timareta ssp. thelxinoe* | Peru | -6.4515 | -76.2977 | pending | | 3,492,663 | 66.21 | 1.70 | CS |
| MJ12-3135 | MJ12-3135 | *Heliconius timareta ssp. thelxinoe* | Peru | -6.4547 | -76.2994 | pending | | 3,708,004 | 70.29 | 1.70 | CS |
| MJ12-3405 | MJ12-3405 | *Heliconius timareta ssp. thelxinoe* | Peru | -6.4540 | -76.3002 | pending | | 3,653,870 | 69.17 | 1.84 | CS |
| MJ12-3409 | MJ12-3409 | *Heliconius timareta ssp. thelxinoe* | Peru | -6.4537 | -76.2981 | pending | | 3,671,396 | 69.59 | 1.70 | CS |
| MJ12-3231 | MJ12-3231 | *Heliconius timareta ssp. thelxinoe* | Peru | -6.4515 | -76.2977 | pending | | 3,449,406 | 65.42 | 1.66 | CS |
| MJ12-3219 | MJ12-3219 | *Heliconius timareta ssp. thelxinoe* | Peru | -6.4519 | -76.2985 | pending | | 3,673,933 | 69.66 | 1.68 | CS |
| MJ12-3370 | MJ12-3370 | *Heliconius timareta ssp. thelxinoe* | Peru | -6.4528 | -76.2862 | pending | | 3,662,416 | 69.24 | 1.96 | CS |
| MJ12-3408 | MJ12-3408 | *Heliconius timareta ssp. thelxinoe* | Peru | -6.4519 | -76.2985 | pending | | 3,737,857 | 70.86 | 1.70 | CS |
| MJ12-3136 | MJ12-3136 | *Heliconius timareta ssp. thelxinoe* | Peru | -6.4547 | -76.2994 | pending | | 3,563,214 | 67.57 | 1.67 | CS |
| MJ12-3223 | MJ12-3223 | *Heliconius timareta ssp. thelxinoe* | Peru | -6.4515 | -76.2977 | pending | | 3,450,844 | 65.47 | 1.62 | CS |
| MJ12-3401 | MJ12-3401 | *Heliconius timareta ssp. thelxinoe* | Peru | -6.4528 | -76.2862 | pending | | 3,695,405 | 70.18 | 1.52 | CS |
| MJ12-3313 | MJ12-3313 | *Heliconius timareta ssp. thelxinoe* | Peru | -6.4547 | -76.2994 | pending | | 3,433,603 | 64.96 | 1.89 | CS |
| MJ12-3400 | MJ12-3400 | *Heliconius timareta ssp. thelxinoe* | Peru | -6.4515 | -76.2977 | pending | | 3,728,573 | 70.66 | 1.72 | CS |
| MJ12-3318 | MJ12-3318 | *Heliconius timareta ssp. thelxinoe* | Peru | -6.4547 | -76.2994 | pending | | 3,598,328 | 68.15 | 1.79 | CS |
| MJ12-3378 | MJ12-3378 | *Heliconius timareta ssp. thelxinoe* | Peru | -6.4519 | -76.2985 | pending | | 3,691,987 | 69.98 | 1.72 | CS |
| MJ12-3404 | MJ12-3404 | *Heliconius timareta ssp. thelxinoe* | Peru | -6.4540 | -76.3002 | pending | | 3,687,744 | 69.92 | 1.68 | CS |
| MJ12-3406 | MJ12-3406 | *Heliconius timareta ssp. thelxinoe* | Peru | -6.4537 | -76.2981 | pending | | 3,704,412 | 70.16 | 1.79 | CS |
| MJ12-3174 | MJ12-3174 | *Heliconius timareta ssp. thelxinoe* | Peru | -6.4547 | -76.2994 | pending | | 3,658,409 | 69.17 | 1.95 | CS |
| MJ12-3390 | MJ12-3390 | *Heliconius timareta ssp. thelxinoe* | Peru | -6.4528 | -76.2862 | pending | | 3,699,234 | 70.01 | 1.86 | CS |
| thxn.MJ12-3221 | MJ12-3221 | *Heliconius timareta ssp. thelxinoe* | Peru | -6.4519 | -76.2985 | SAMEA104585110 | Van Belleghem *et al.*[13] | 3,819,091 | 72.32 | 1.81 | WGS |
| thxn.MJ12-3233 | MJ12-3233 | *Heliconius timareta ssp. thelxinoe* | Peru | -5.6546 | -77.6938 | SAMEA104585111 | Van Belleghem *et al.*[13] | 3,792,490 | 71.77 | 1.86 | WGS |
| thxn.MJ12-3308 | MJ12-3308 | *Heliconius timareta ssp. thelxinoe* | Peru | -6.4519 | -76.2985 | SAMEA104585112 | Van Belleghem *et al.*[13] | 3,760,503 | 71.06 | 2.02 | WGS |
| txn.MJ11-3339 | MJ12-3339 | *Heliconius timareta ssp. thelxinoe* | Peru | -5.6546 | -77.6938 | SAMEA104585113 | Van Belleghem *et al.*[13] | 3,523,213 | 66.66 | 1.89 | WGS |
| txn.MJ11-3340 | MJ12-3340 | *Heliconius timareta ssp. thelxinoe* | Peru | -5.6546 | -77.6938 | SAMEA104585114 | Van Belleghem *et al.*[13] | 3,437,195 | 65.07 | 1.83 | WGS |
| txn.MJ12-3460 | MJ12-3460 | *Heliconius timareta ssp. thelxinoe* | Peru | -5.6546 | -77.6938 | SAMEA104585115 | Van Belleghem *et al.*[13] | 3,651,061 | 69.03 | 1.96 | WGS |
| thxn.JM57 | JM-09-57 | *Heliconius timareta ssp. thelxinoe* | Peru | -6.4528 | -76.2987 | SAMEA1919254 | Martin *et al.*[14] | 3,964,602 | 74.96 | 1.95 | WGS |
| thxn.JM86 | JM-09-86 | *Heliconius timareta ssp. thelxinoe* | Peru | -6.4528 | -76.2987 | SAMEA1919263 | Martin *et al.*[14] | 3,969,848 | 74.91 | 2.15 | WGS |
| thxn.JM313 | JM-09-313 | *Heliconius timareta ssp. thelxinoe* | Peru | -6.4584 | -76.2877 | SAMEA1919266 | Martin *et al.*[14] | 3,953,763 | 74.65 | 2.09 | WGS |
| thxn.JM84 | JM-09-84 | *Heliconius timareta ssp. thelxinoe* | Peru | -6.4528 | -76.2987 | SAMEA1919273 | Martin *et al.*[14] | 3,819,757 | 72.27 | 1.89 | WGS |
| CAM009187 | CAM009187 | *Heliconius timareta ssp. timareta f. contigua* | Ecuador | -1.3980 | -78.1781 | pending | | 3,755,103 | 71.11 | 1.80 | CS |
| CAM009186 | CAM009186 | *Heliconius timareta ssp. timareta f. contigua* | Ecuador | -1.3980 | -78.1781 | pending | | 3,714,942 | 70.37 | 1.77 | CS |
| CAM009185 | CAM009185 | *Heliconius timareta ssp. timareta f. contigua* | Ecuador | -1.3980 | -78.1781 | pending | | 3,724,200 | 70.56 | 1.76 | CS |
| CAM009183 | CAM009183 | *Heliconius timareta ssp. timareta f. contigua* | Ecuador | -1.3980 | -78.1781 | pending | | 3,732,851 | 70.71 | 1.77 | CS |
| CAM009177 | CAM009177 | *Heliconius timareta ssp. timareta f. contigua* | Ecuador | -1.3980 | -78.1781 | pending | | 3,768,012 | 71.35 | 1.80 | CS |

| SequenceID | EarthCapeID | Taxon Name | Country | Latitude | Longitude | Accession | Publication | Called sites | Genotyped sites after filters (%) | Proportion of hetero-zygous genotypes (%) | Data Type (WGS…whole genome re-sequencing, CS…capture sequencing) |
|---|---|---|---|---|---|---|---|---|---|---|---|
| CAM009175 | CAM009175 | *Heliconius timareta ssp. timareta f. contigua* | Ecuador | -1.3980 | -78.1781 | pending | | 3,742,771 | 70.91 | 1.75 | CS |
| CAM009180 | CAM009180 | *Heliconius timareta ssp. timareta f. contigua* | Ecuador | -1.3980 | -78.1781 | SAMEA1322935; pending | The Heliconius Consortium[15]; RAD; SureSelect | 3,752,197 | 71.02 | 1.85 | CS |
| CAM016719 | CAM016719 | *Heliconius timareta ssp. timareta f. contigua* | Ecuador | -1.3980 | -78.1781 | pending | | 3,759,873 | 71.19 | 1.82 | CS |
| CAM016716 | CAM016716 | *Heliconius timareta ssp. timareta f. contigua* | Ecuador | -1.3980 | -78.1781 | pending | | 3,730,788 | 70.61 | 1.85 | CS |
| CAM016715 | CAM016715 | *Heliconius timareta ssp. timareta f. contigua* | Ecuador | -1.3980 | -78.1781 | pending | | 3,599,854 | 68.22 | 1.73 | CS |
| CAM011413 | CAM011413 | *Heliconius timareta ssp. timareta f. contigua* | Ecuador | -1.3980 | -78.1781 | pending | | 3,726,722 | 70.57 | 1.80 | CS |
| CAM011439 | CAM011439 | *Heliconius timareta ssp. timareta f. timareta* | Ecuador | -1.3980 | -78.1781 | pending | | 3,789,280 | 71.74 | 1.83 | CS |
| CAM009440 | CAM009440 | *Heliconius timareta ssp. timareta f. timareta* | Ecuador | -1.3980 | -78.1781 | pending | | 3,822,542 | 72.44 | 1.73 | CS |
| CAM009426 | CAM009426 | *Heliconius timareta ssp. timareta f. timareta* | Ecuador | -1.3980 | -78.1781 | pending | | 3,816,404 | 72.23 | 1.86 | CS |
| CAM009421 | CAM009421 | *Heliconius timareta ssp. timareta f. timareta* | Ecuador | -1.3980 | -78.1781 | pending | | 3,801,719 | 72.11 | 1.64 | CS |
| CAM009419 | CAM009419 | *Heliconius timareta ssp. timareta f. timareta* | Ecuador | -1.3980 | -78.1781 | pending | | 3,788,443 | 71.73 | 1.81 | CS |
| CAM009227 | CAM009227 | *Heliconius timareta ssp. timareta f. timareta* | Ecuador | -1.4530 | -78.1070 | pending | | 3,726,694 | 70.61 | 1.75 | CS |
| CAM009169 | CAM009169 | *Heliconius timareta ssp. timareta f. timareta* | Ecuador | -1.3980 | -78.1781 | pending | | 3,756,296 | 71.12 | 1.81 | CS |
| CAM008533 | CAM008533 | *Heliconius timareta ssp. timareta f. timareta* | Ecuador | -1.3980 | -78.1781 | SAMEA1094484; pending | Nadeau *et al.*[19]; SureSelect | 3,841,553 | 72.69 | 1.88 | CS |
| CAM009178 | CAM009178 | *Heliconius timareta ssp. timareta f. timareta* | Ecuador | -1.3980 | -78.1781 | SAMEA3670574; pending | Wallbank *et al.*[8] | 3,678,631 | 69.72 | 1.73 | CS |

**Supplementary Table 2:** Position, composite likelihood-ratio statistics (CLR) and strength of selection ($\alpha$, $2N_es$, and $s$) for the highest CLR and the smallest $\alpha$ value on each colour pattern scaffold ($\alpha_{min}$) for the *H. melpomene*-clade. Additional relevant peaks on scaffolds are also given. Data are from SweepFinder2[3,4] runs with background site frequency spectrum estimated from background scaffolds.

| Population | Locus | Scaffold | Position | CLR | $\alpha$ | $2N_es$ | $s$ | Position ($\alpha_{min}$) | CLR ($\alpha_{min}$) | $\alpha_{min}$ | $2N_es$ ($\alpha_{min}$) | $s$ ($\alpha_{min}$) |
|---|---|---|---|---|---|---|---|---|---|---|---|---|
| *H. besckei* | *aristaless* | Hmel201011 | 2488332 | 14 | 244.24 | 848 | 0.001 | 2623214 | 8 | 79.09 | 2618 | 0.003 |
| *H. c. chioneus* | *aristaless* | Hmel201011 | 2637747 | 41 | 129.13 | 4529 | 0.002 | 2638447 | 38 | 126.42 | 4626 | 0.002 |
| *H. c. cydnides* | *aristaless* | Hmel201011 | 2613872 | 106 | 63 | 8052 | 0.004 | 2613822 | 101 | 62.97 | 8056 | 0.004 |
| *H. c. weymeri gustavi* | *aristaless* | Hmel201011 | 2598794 | 64 | 109.51 | 4750 | 0.002 | 2600094 | 39 | 87.83 | 5923 | 0.003 |
| *H. c. weymeri weymeri* | *aristaless* | Hmel201011 | 2595343 | 122 | 74.63 | 6265 | 0.003 | 2600194 | 68 | 52.06 | 8981 | 0.004 |
| *H. c. zelinde* | *aristaless* | Hmel201011 | 2637640 | 39 | 132.71 | 4674 | 0.002 | 2637640 | 39 | 132.71 | 4674 | 0.002 |
| *H. elevatus Ecuador* | *aristaless* | Hmel201011 | 2671769 | 64 | 140.74 | 6525 | 0.002 | 2671769 | 64 | 140.74 | 6525 | 0.002 |
| *H. heurippa* | *aristaless* | Hmel201011 | 2648749 | 125 | 43.49 | 7946 | 0.005 | 2647499 | 97 | 40.42 | 8550 | 0.006 |
| *H. m. amaryllis* | *aristaless* | Hmel201011 | 2638339 | 19 | 254.65 | 3387 | 0.001 | 2725509 | 7 | 198.75 | 4340 | 0.001 |
| *H. m. cythera* | *aristaless* | Hmel201011 | 2616846 | 80 | 71.96 | 8271 | 0.003 | 2623247 | 72 | 30.58 | 19461 | 0.008 |
| *H. m. ECU* | *aristaless* | Hmel201011 | 2641567 | 31 | 202.82 | 3773 | 0.001 | 2641617 | 31 | 201.69 | 3794 | 0.001 |
| *H. m. malleti COL* | *aristaless* | Hmel201011 | 2563116 | 33 | 436.43 | 1688 | 0.001 | 2725394 | 15 | 214.26 | 3439 | 0.001 |
| *H. m. malleti ECU* | *aristaless* | Hmel201011 | 2643054 | 25 | 606.14 | 1501 | 0 | 2725514 | 4 | 194.61 | 4675 | 0.001 |
| *H. m. melpomene COL* | *aristaless* | Hmel201011 | 2623375 | 200 | 33.27 | 22438 | 0.007 | 2623575 | 196 | 33.07 | 22575 | 0.007 |
| *H. m. melpomene FG* | *aristaless* | Hmel201011 | 2518352 | 27 | 489.17 | 1083 | 0 | 2583809 | 7 | 171.11 | 3097 | 0.001 |
| *H. m. melpomene PAN* | *aristaless* | Hmel201011 | 2601790 | 44 | 216.07 | 3070 | 0.001 | 2600890 | 41 | 111.65 | 5942 | 0.002 |
| *H. m. meriana* | *aristaless* | Hmel201011 | 2584942 | 16 | 983.69 | 388 | 0 | 2483584 | 1 | 251.96 | 1516 | 0.001 |
| *H. m. nanna NORTH* | *aristaless* | Hmel201011 | 2542627 | 46 | 158.72 | 3705 | 0.001 | 2545828 | 21 | 120.49 | 4881 | 0.002 |
| *H. m. nanna SOUTH* | *aristaless* | Hmel201011 | 2753296 | 30 | 82.03 | 7169 | 0.003 | 2636583 | 27 | 64.35 | 9140 | 0.004 |
| *H. m. plesseni* | *aristaless* | Hmel201011 | 2641847 | 21 | 276.52 | 2612 | 0.001 | 2740705 | 9 | 264.91 | 2727 | 0.001 |
| *H. m. rosina* | *aristaless* | Hmel201011 | 2648668 | 57 | 164.68 | 3222 | 0.001 | 2661869 | 28 | 95.48 | 5558 | 0.002 |
| *H. m. vicina* | *aristaless* | Hmel201011 | 2641654 | 33 | 94.72 | 7881 | 0.003 | 2641504 | 31 | 93.92 | 7948 | 0.003 |
| *H. m. vulcanus* | *aristaless* | Hmel201011 | 2669703 | 43 | 249.33 | 1953 | 0.001 | 2661802 | 13 | 142.8 | 3410 | 0.002 |
| *H. m. xenoclea* | *aristaless* | Hmel201011 | 2642090 | 24 | 233.79 | 2904 | 0.001 | 2641640 | 21 | 162.04 | 4189 | 0.001 |
| *H. pachinus* | *aristaless* | Hmel201011 | 2702891 | 99 | 37.43 | 14517 | 0.006 | 2704441 | 44 | 35.6 | 15263 | 0.007 |
| *H. t. florencia* | *aristaless* | Hmel201011 | 2669550 | 193 | 23.58 | 23345 | 0.01 | 2668800 | 108 | 23.34 | 23584 | 0.01 |
| | | | 2640195 | 180 | 38.35 | 14352 | 0.006 | 2640045 | 176 | 37.69 | 14603 | 0.006 |
| *H. t. linaresi* | *aristaless* | Hmel201011 | 2634480 | 180 | 27.08 | 18189 | 0.009 | 2634180 | 127 | 26.92 | 18297 | 0.009 |
| *H. t. ssp. nov. ECU* | *aristaless* | Hmel201011 | 2666249 | 159 | 35.77 | 16212 | 0.007 | 2664749 | 140 | 31.57 | 18367 | 0.007 |
| *H. t. thelxinoe* | *aristaless* | Hmel201011 | 2674171 | 59 | 103.18 | 4818 | 0.002 | 2631367 | 37 | 98.32 | 5056 | 0.002 |
| *H. t. timareta f. contigua* | *aristaless* | Hmel201011 | 2674005 | 146 | 35.27 | 14908 | 0.007 | 2673455 | 115 | 34.82 | 15102 | 0.007 |
| *H. t. timareta f. timareta* | *aristaless* | Hmel201011 | 2674076 | 77 | 58.86 | 9727 | 0.004 | 2631017 | 64 | 47.35 | 12090 | 0.005 |
| *H. t. ssp. nov. COL* | *aristaless* | Hmel201011 | 2669779 | 164 | 26.14 | 17359 | 0.009 | 2672479 | 77 | 23.57 | 19257 | 0.01 |
| *H. besckei* | *WntA* | Hmel210004 | 1559377 | 27 | 108.61 | 1983 | 0.002 | 1566878 | 19 | 40.6 | 5304 | 0.005 |
| *H. c. chioneus* | *WntA* | Hmel210004 | 1620799 | 161 | 28.33 | 18989 | 0.008 | 1626550 | 119 | 25.19 | 21352 | 0.009 |
| *H. c. cydnides* | *WntA* | Hmel210004 | 1811842 | 173 | 27.05 | 21129 | 0.008 | 1817892 | 143 | 22.09 | 25874 | 0.01 |
| *H. c. weymeri gustavi* | *WntA* | Hmel210004 | 1806398 | 348 | 13.78 | 38564 | 0.016 | 1805348 | 344 | 13.57 | 39160 | 0.016 |

| Population | Locus | Scaffold | Position | CLR | $\alpha$ | $2N_es$ | $s$ | Position ($\alpha_{min}$) | CLR ($\alpha_{min}$) | $\alpha_{min}$ | $2N_es$ ($\alpha_{min}$) | $s$ ($\alpha_{min}$) |
|---|---|---|---|---|---|---|---|---|---|---|---|---|
| *H. c. weymeri weymeri* | *WntA* | Hmel210004 | 1810765 | 237 | 17.17 | 32190 | 0.013 | 1811415 | 222 | 17.04 | 32431 | 0.013 |
| *H. c. zelinde* | *WntA* | Hmel210004 | 1620832 | 128 | 31.44 | 18426 | 0.007 | 1620482 | 120 | 31.14 | 18602 | 0.007 |
| *H. elevatus ECU* | *WntA* | Hmel210004 | 1828231 | 339 | 25.25 | 33900 | 0.009 | 1827631 | 324 | 24.76 | 34574 | 0.009 |
| *H. heurippa* | *WntA* | Hmel210004 | 1560084 | 247 | 21.52 | 15644 | 0.01 | 1567084 | 239 | 20.14 | 16718 | 0.011 |
| *H. m. amaryllis* | *WntA* | Hmel210004 | 1809664 | 186 | 44.02 | 17940 | 0.005 | 1810014 | 180 | 40.31 | 19589 | 0.006 |
| *H. m. cythera* | *WntA* | Hmel210004 | 1811224 | 174 | 29.08 | 19900 | 0.008 | 1811424 | 171 | 28.97 | 19975 | 0.008 |
| *H. m. ecuadoriensis* | *WntA* | Hmel210004 | 1848718 | 304 | 32.77 | 20532 | 0.007 | 1809867 | 113 | 24.11 | 27915 | 0.009 |
| | | | 1808717 | 226 | 24.59 | 27368 | 0.009 | 1809867 | 113 | 24.11 | 27915 | 0.009 |
| *H. m. malleti COL* | *WntA* | Hmel210004 | 1621626 | 235 | 24.39 | 28389 | 0.009 | 1621326 | 170 | 24.26 | 28533 | 0.009 |
| *H. m. malleti ECU* | *WntA* | Hmel210004 | 1809624 | 306 | 30.86 | 27223 | 0.007 | 1810125 | 302 | 30.03 | 27982 | 0.008 |
| *H. m. melpomene COL* | *WntA* | Hmel210004 | 1631681 | 531 | 12.33 | 58771 | 0.018 | 1630681 | 251 | 12.3 | 58890 | 0.018 |
| *H. m. melpomene FG* | *WntA* | Hmel210004 | 1568298 | 101 | 21.42 | 19513 | 0.01 | 1567298 | 96 | 21.08 | 19829 | 0.01 |
| *H. m. melpomene PAN* | *WntA* | Hmel210004 | 1629713 | 479 | 12.61 | 51277 | 0.018 | 1629363 | 444 | 12.56 | 51484 | 0.018 |
| *H. m. meriana* | *WntA* | Hmel210004 | 1855774 | 100 | 93.21 | 3613 | 0.002 | 1623664 | 80 | 48.71 | 6914 | 0.004 |
| *H. m. nanna NORTH* | *WntA* | Hmel210004 | 1806251 | 237 | 10.14 | 53581 | 0.022 | 1807451 | 183 | 10.06 | 54028 | 0.022 |
| *H. m. nanna SOUTH* | *WntA* | Hmel210004 | 1783553 | 58 | 15.1 | 35991 | 0.015 | 1784553 | 41 | 15.01 | 36193 | 0.015 |
| *H. m. plesseni* | *WntA* | Hmel210004 | 1829355 | 1098 | 6.3 | 95215 | 0.035 | 1830905 | 1081 | 6.28 | 95549 | 0.035 |
| *H. m. rosina* | *WntA* | Hmel210004 | 1568042 | 195 | 24.31 | 18977 | 0.009 | 1627296 | 44 | 22.33 | 20659 | 0.01 |
| *H. m. vicina* | *WntA* | Hmel210004 | 1622166 | 179 | 24.43 | 29654 | 0.009 | 1620566 | 151 | 23.21 | 31214 | 0.01 |
| *H. m. vulcanus* | *WntA* | Hmel210004 | 1625201 | 231 | 19.04 | 21328 | 0.011 | 1624201 | 126 | 18.97 | 21418 | 0.011 |
| *H. m. xenoclea* | *WntA* | Hmel210004 | 1811430 | 971 | 4.54 | 118013 | 0.049 | 1812130 | 965 | 4.54 | 118070 | 0.049 |
| *H. pachinus* | *WntA* | Hmel210004 | 1805390 | 245 | 11.06 | 47778 | 0.02 | 1805340 | 245 | 11.06 | 47780 | 0.02 |
| *H. t. florencia* | *WntA* | Hmel210004 | 1566574 | 542 | 11.94 | 43044 | 0.018 | 1566474 | 542 | 11.93 | 43059 | 0.018 |
| *H. t. linaresi* | *WntA* | Hmel210004 | 1825099 | 215 | 23.03 | 21135 | 0.01 | 1825249 | 215 | 23.01 | 21150 | 0.01 |
| *H. t. ssp. nov. ECU* | *WntA* | Hmel210004 | 1571479 | 414 | 20.96 | 22770 | 0.01 | 1566878 | 397 | 16.37 | 29147 | 0.013 |
| *H. t. thelxinoe* | *WntA* | Hmel210004 | 1765448 | 170 | 23.36 | 17044 | 0.009 | 1625441 | 113 | 14.67 | 27142 | 0.015 |
| *H. t. timareta f. contigua* | *WntA* | Hmel210004 | 1572726 | 216 | 20.37 | 18943 | 0.011 | 1629379 | 78 | 12.71 | 30349 | 0.017 |
| *H. t. timareta f. timareta* | *WntA* | Hmel210004 | 1567741 | 300 | 13.42 | 31394 | 0.016 | 1568691 | 221 | 13.38 | 31496 | 0.016 |
| *H. t. ssp. nov. COL* | *WntA* | Hmel210004 | 1567277 | 421 | 12.56 | 37584 | 0.017 | 1880146 | 149 | 9.41 | 50157 | 0.023 |
| *H. besckei* | *cortex* | Hmel215006 | 662442 | 28 | 56.91 | 5712 | 0.006 | 925699 | 11 | 10.58 | 30722 | 0.031 |
| *H. c. chioneus* | *cortex* | Hmel215006 | 1209780 | 459 | 16.81 | 50815 | 0.021 | 924667 | 29 | 13.46 | 63454 | 0.026 |
| *H. c. cydnides* | *cortex* | Hmel215006 | 1238215 | 236 | 35.36 | 22363 | 0.01 | 824294 | 8 | 15.26 | 51828 | 0.023 |
| *H. c. weymeri gustavi* | *cortex* | Hmel215006 | 1329024 | 696 | 9.75 | 68950 | 0.036 | 1326374 | 343 | 9.57 | 70208 | 0.036 |
| | | | 1220571 | 627 | 14.4 | 46666 | 0.024 | 1225721 | 265 | 14.1 | 47661 | 0.025 |
| *H. c. weymeri weymeri* | *cortex* | Hmel215006 | 1337975 | 2411 | 5.3 | 115568 | 0.065 | 1334724 | 2289 | 5.28 | 116071 | 0.065 |
| | | | 1218021 | 367 | 20.74 | 29538 | 0.017 | 1215221 | 325 | 20.08 | 30501 | 0.017 |
| *H. c. zelinde* | *cortex* | Hmel215006 | 1218589 | 262 | 25.2 | 35877 | 0.014 | 924529 | 71 | 9.32 | 97037 | 0.038 |
| *H. elevatus ECU* | *cortex* | Hmel215006 | 1563763 | 242 | 55.24 | 25155 | 0.007 | 923334 | 34 | 17 | 81744 | 0.021 |
| *H. heurippa* | *cortex* | Hmel215006 | 1446977 | 724 | 16.71 | 26945 | 0.02 | 924205 | 211 | 7.68 | 58659 | 0.044 |
| *H. m. amaryllis* | *cortex* | Hmel215006 | 1575917 | 979 | 10.69 | 75277 | 0.033 | 1581867 | 835 | 9.75 | 82558 | 0.036 |
| *H. m. cythera* | *cortex* | Hmel215006 | 1100490 | 1713 | 3.35 | 205197 | 0.104 | 1090939 | 1369 | 3.29 | 209382 | 0.106 |
| | | | 1229846 | 1484 | 6.58 | 104613 | 0.053 | 1224346 | 1323 | 6.48 | 106226 | 0.054 |
| *H. m. ecuadoriensis* | *cortex* | Hmel215006 | 1459579 | 335 | 30.74 | 30944 | 0.012 | 924906 | 110 | 7.51 | 126705 | 0.047 |

| Population | Locus | Scaffold | Position | CLR | $\alpha$ | $2N_es$ | $s$ | Position ($\alpha_{min}$) | CLR ($\alpha_{min}$) | $\alpha_{min}$ | $2N_es$ ($\alpha_{min}$) | $s$ ($\alpha_{min}$) |
|---|---|---|---|---|---|---|---|---|---|---|---|---|
| *H. m. malleti COL* | *cortex* | Hmel215006 | 1105290 | 335 | 16.2 | 60827 | 0.022 | 923581 | 88 | 10.88 | 90534 | 0.033 |
| *H. m. malleti ECU* | *cortex* | Hmel215006 | 1098623 | 234 | 29.97 | 40361 | 0.012 | 924714 | 145 | 9.5 | 127251 | 0.038 |
| *H. m. melpomene COL* | *cortex* | Hmel215006 | 1095016 | 604 | 12.97 | 68498 | 0.027 | 1095166 | 602 | 12.96 | 68537 | 0.027 |
| *H. m. melpomene FG* | *cortex* | Hmel215006 | 1557309 | 506 | 18.61 | 36692 | 0.019 | 826094 | 12 | 13.69 | 49863 | 0.025 |
| *H. m. melpomene PAN* | *cortex* | Hmel215006 | 1096916 | 857 | 5.36 | 164465 | 0.066 | 1089216 | 722 | 5.03 | 175240 | 0.07 |
| *H. m. meriana* | *cortex* | Hmel215006 | 1220307 | 235 | 28.78 | 17789 | 0.012 | 830538 | 3 | 21.45 | 23866 | 0.016 |
| *H. m. nanna NORTH* | *cortex* | Hmel215006 | 1460407 | 1350 | 6.79 | 108135 | 0.051 | 1057899 | 1141 | 2.47 | 297164 | 0.141 |
| | | | 1226202 | 1316 | 4.29 | 171062 | 0.081 | 1230152 | 811 | 4.25 | 172801 | 0.082 |
| | | | 1062599 | 1198 | 2.49 | 295257 | 0.14 | 1057899 | 1141 | 2.47 | 297164 | 0.141 |
| | | | 1577710 | 1170 | 6.83 | 107604 | 0.051 | 1570059 | 938 | 6.7 | 109603 | 0.052 |
| *H. m. nanna SOUTH* | *cortex* | Hmel215006 | 1130468 | 173 | 7.86 | 93493 | 0.044 | 1123618 | 75 | 7.69 | 95532 | 0.045 |
| *H. m. plesseni* | *cortex* | Hmel215006 | 1237265 | 2989 | 4.7 | 157635 | 0.074 | 1235965 | 2981 | 4.7 | 157702 | 0.074 |
| | | | 1366372 | 1090 | 17.92 | 41316 | 0.02 | 1304818 | 214 | 7.93 | 93420 | 0.044 |
| | | | 1447976 | 958 | 25.36 | 29196 | 0.014 | 1380022 | 742 | 23.55 | 31445 | 0.015 |
| *H. m. rosina* | *cortex* | Hmel215006 | 1069455 | 1051 | 2.77 | 225945 | 0.125 | 1066255 | 871 | 2.75 | 227241 | 0.126 |
| *H. m. vicina* | *cortex* | Hmel215006 | 1105326 | 372 | 6.1 | 145612 | 0.058 | 1091675 | 317 | 5.39 | 164792 | 0.066 |
| *H. m. vulcanus* | *cortex* | Hmel215006 | 1200605 | 615 | 12.93 | 47085 | 0.027 | 1088852 | 171 | 6.18 | 98467 | 0.056 |
| *H. m. xenoclea* | *cortex* | Hmel215006 | 1543692 | 1166 | 8.72 | 91594 | 0.04 | 1055866 | 486 | 6.56 | 121755 | 0.054 |
| | | | 1459887 | 1045 | 12.73 | 62761 | 0.028 | 1459687 | 1044 | 12.72 | 62769 | 0.028 |
| *H. pachinus* | *cortex* | Hmel215006 | 1458120 | 570 | 16.83 | 44000 | 0.021 | 925602 | 111 | 6.76 | 109519 | 0.052 |
| *H. t. florencia* | *cortex* | Hmel215006 | 1060718 | 639 | 8.09 | 93800 | 0.043 | 1068818 | 462 | 7.89 | 96195 | 0.044 |
| *H. t. linaresi* | *cortex* | Hmel215006 | 1328608 | 871 | 11.56 | 60838 | 0.03 | 1329908 | 838 | 11.47 | 61324 | 0.03 |
| *H. t. ssp. nov. ECU* | *cortex* | Hmel215006 | 1056672 | 532 | 8.3 | 87549 | 0.042 | 1057872 | 444 | 8.18 | 88791 | 0.043 |
| *H. t. thelxinoe* | *cortex* | Hmel215006 | 1245780 | 189 | 69.92 | 9208 | 0.005 | 926617 | 21 | 19.85 | 32434 | 0.017 |
| *H. t. timareta f. contigua* | *cortex* | Hmel215006 | 1458076 | 523 | 21.79 | 29909 | 0.016 | 930404 | 21 | 8.21 | 79348 | 0.042 |
| *H. t. timareta f. timareta* | *cortex* | Hmel215006 | 1098821 | 613 | 8.35 | 77661 | 0.041 | 1091971 | 572 | 7.51 | 86375 | 0.046 |
| *H. t. ssp. nov. COL* | *cortex* | Hmel215006 | 1459519 | 599 | 16.07 | 39527 | 0.022 | 1070451 | 554 | 5.89 | 107840 | 0.059 |
| *H. besckei* | *optix* | Hmel218003 | 736567 | 26 | 211.68 | 905 | 0.001 | 868677 | 22 | 66.88 | 2865 | 0.003 |
| *H. c. chioneus* | *optix* | Hmel218003 | 788367 | 132 | 45.54 | 11564 | 0.005 | 789017 | 129 | 45.12 | 11671 | 0.005 |
| *H. c. cydnides* | *optix* | Hmel218003 | 637588 | 155 | 46.8 | 10480 | 0.005 | 789794 | 131 | 42.03 | 11670 | 0.005 |
| *H. c. weymeri gustavi* | *optix* | Hmel218003 | 624737 | 176 | 38.09 | 11148 | 0.006 | 625137 | 173 | 38 | 11175 | 0.006 |
| *H. c. weymeri weymeri* | *optix* | Hmel218003 | 1019652 | 220 | 102.97 | 3284 | 0.002 | 786128 | 122 | 53.71 | 6297 | 0.004 |
| *H. c. zelinde* | *optix* | Hmel218003 | 789114 | 116 | 48.94 | 10602 | 0.004 | 789764 | 112 | 48.27 | 10749 | 0.005 |
| *H. elevatus ECU* | *optix* | Hmel218003 | 787076 | 224 | 44.11 | 15829 | 0.005 | 857131 | 68 | 39.13 | 17844 | 0.006 |
| *H. heurippa* | *optix* | Hmel218003 | 857729 | 525 | 18.23 | 11544 | 0.011 | 853979 | 489 | 17.77 | 11848 | 0.012 |
| | | | 781223 | 481 | 27.33 | 7701 | 0.008 | 785574 | 435 | 24.94 | 8438 | 0.008 |
| *H. m. amaryllis* | *optix* | Hmel218003 | 786183 | 357 | 32.58 | 14300 | 0.007 | 784733 | 279 | 32.07 | 14528 | 0.007 |
| *H. m. cythera* | *optix* | Hmel218003 | 838145 | 736 | 11.82 | 33955 | 0.018 | 833645 | 509 | 11.76 | 34137 | 0.018 |
| *H. m. ECU* | *optix* | Hmel218003 | 811137 | 200 | 47.73 | 10123 | 0.005 | 842388 | 17 | 44.12 | 10951 | 0.005 |
| *H. m. malleti COL* | *optix* | Hmel218003 | 814383 | 329 | 28.73 | 17414 | 0.008 | 843034 | 206 | 22.26 | 22480 | 0.01 |
| *H. m. malleti ECU* | *optix* | Hmel218003 | 814624 | 255 | 53.69 | 12199 | 0.004 | 843427 | 189 | 31.04 | 21102 | 0.007 |
| *H. m. melpomene COL* | *optix* | Hmel218003 | 672284 | 119 | 138.06 | 3609 | 0.002 | 623580 | 95 | 55.09 | 9046 | 0.004 |
| *H. m. melpomene FG* | *optix* | Hmel218003 | 649462 | 141 | 49.87 | 7416 | 0.004 | 645511 | 80 | 45.72 | 8089 | 0.005 |

| **Population** | **Locus** | **Scaffold** | **Position** | **CLR** | $\alpha$ | $2N_e s$ | *s* | **Position** ($\alpha_{min}$) | **CLR** ($\alpha_{min}$) | $\alpha_{min}$ | $2N_e s$ ($\alpha_{min}$) | *s* ($\alpha_{min}$) |
|---|---|---|---|---|---|---|---|---|---|---|---|---|
| *H. m. melpomene PAN* | *optix* | Hmel218003 | 838170 | 647 | 13.74 | 33226 | 0.016 | 837219 | 529 | 13.72 | 33277 | 0.016 |
| *H. m. meriana* | *optix* | Hmel218003 | 801534 | 1250 | 9.45 | 35360 | 0.023 | 811585 | 826 | 9.28 | 35993 | 0.023 |
| *H. m. nanna NORTH* | *optix* | Hmel218003 | 782525 | 306 | 38.5 | 12092 | 0.006 | 646316 | 90 | 27.67 | 16823 | 0.008 |
| *H. m. nanna SOUTH* | *optix* | Hmel218003 | 726637 | 41 | 65.72 | 7084 | 0.003 | 727838 | 38 | 62.99 | 7391 | 0.003 |
| *H. m. plesseni* | *optix* | Hmel218003 | 783431 | 2371 | 6.97 | 41978 | 0.03 | 655174 | 1937 | 5.91 | 49501 | 0.036 |
| | | | 643924 | 2174 | 6.07 | 48223 | 0.035 | 655174 | 1937 | 5.91 | 49501 | 0.036 |
| | | | 732278 | 1638 | 6.21 | 47109 | 0.034 | 728328 | 1466 | 6.17 | 47423 | 0.034 |
| *H. m. rosina* | *optix* | Hmel218003 | 847932 | 585 | 13.15 | 26357 | 0.016 | 840982 | 463 | 12.78 | 27129 | 0.017 |
| *H. m. vicina* | *optix* | Hmel218003 | 790999 | 475 | 10.58 | 47091 | 0.021 | 796800 | 320 | 10.33 | 48254 | 0.021 |
| *H. m. vulcanus* | *optix* | Hmel218003 | 848005 | 759 | 8.83 | 32135 | 0.024 | 840154 | 609 | 8.53 | 33270 | 0.025 |
| *H. m. xenoclea* | *optix* | Hmel218003 | 727532 | 1182 | 9.74 | 37910 | 0.022 | 725882 | 917 | 9.73 | 37935 | 0.022 |
| *H. pachinus* | *optix* | Hmel218003 | 648265 | 289 | 30.16 | 14716 | 0.007 | 646315 | 193 | 29.92 | 14838 | 0.007 |
| *H. t. florencia* | *optix* | Hmel218003 | 705381 | 201 | 84.98 | 4616 | 0.003 | 674830 | 70 | 44.94 | 8729 | 0.005 |
| *H. t. linaresi* | *optix* | Hmel218003 | 803436 | 408 | 27.11 | 12138 | 0.008 | 802136 | 218 | 26.9 | 12235 | 0.008 |
| *H. t. ssp. nov. ECU* | *optix* | Hmel218003 | 705531 | 186 | 99.32 | 3833 | 0.002 | 671379 | 46 | 56.44 | 6745 | 0.004 |
| *H. t. thelxinoe* | *optix* | Hmel218003 | 788276 | 66 | 117.15 | 2773 | 0.002 | 788276 | 66 | 117.15 | 2773 | 0.002 |
| *H. t. timareta f. contigua* | *optix* | Hmel218003 | 941544 | 152 | 52.4 | 6196 | 0.004 | 941244 | 147 | 52.33 | 6205 | 0.004 |
| *H. t. timareta f. timareta* | *optix* | Hmel218003 | 864840 | 157 | 50.27 | 6313 | 0.004 | 864840 | 157 | 50.27 | 6313 | 0.004 |
| *H. t. ssp. nov. COL* | *optix* | Hmel218003 | 597476 | 284 | 36.8 | 8193 | 0.006 | 616977 | 186 | 24.29 | 12411 | 0.009 |

**Supplementary Table 3:** Position, composite likelihood-ratio statistics (CLR) and strength of selection ($\alpha$, $2N_es$, and $s$) for the highest CLR and the smallest $\alpha$ value on each background scaffold ($\alpha_{min}$) for the *H. melpomene*-clade. Data are from SweepFinder2[3,4] runs with background site frequency spectrum estimated from background scaffolds.

| Population | Scaffold | Position | CLR | $\alpha$ | $2N_es$ | $s$ | Position ($\alpha_{min}$) | CLR ($\alpha_{min}$) | $\alpha_{min}$ | $2N_es$ ($\alpha_{min}$) | $s$ ($\alpha_{min}$) |
|---|---|---|---|---|---|---|---|---|---|---|---|
| *H. besckei* | Hmel204017 | 2202171 | 10 | 549.99 | 649 | 0.001 | 2198120 | 7 | 42.25 | 8445 | 0.008 |
| *H. c. chioneus* | Hmel204017 | 2299755 | 24 | 461.75 | 2371 | 0.001 | 1959329 | 13 | 163.28 | 6705 | 0.002 |
| *H. c. cydnides* | Hmel204017 | 2298643 | 25 | 194.08 | 6303 | 0.002 | 1958116 | 20 | 89.26 | 13704 | 0.004 |
| *H. c. weymeri gustavi* | Hmel204017 | 2003132 | 40 | 121.06 | 10105 | 0.003 | 2003332 | 35 | 118.42 | 10330 | 0.003 |
| *H. c. weymeri weymeri* | Hmel204017 | 2019842 | 42 | 192.6 | 6144 | 0.002 | 2004090 | 29 | 91.35 | 12954 | 0.004 |
| *H. c. zelinde* | Hmel204017 | 2016795 | 31 | 193.11 | 5709 | 0.002 | 2053101 | 10 | 75.43 | 14614 | 0.005 |
| *H. elevatus Ecuador* | Hmel204017 | 2255583 | 12 | 484.36 | 2327 | 0.001 | 1958500 | 4 | 103.58 | 10880 | 0.004 |
| *H. heurippa* | Hmel204017 | 2117854 | 35 | 206.51 | 8319 | 0.002 | 2109703 | 18 | 67.29 | 25532 | 0.006 |
| *H. m. amaryllis* | Hmel204017 | 2095349 | 92 | 72.69 | 11239 | 0.005 | 2110351 | 4 | 54.96 | 14865 | 0.007 |
| *H. m. cythera* | Hmel204017 | 1957699 | 23 | 226.1 | 6867 | 0.002 | 2109212 | 3 | 101.71 | 15266 | 0.004 |
| *H. m. ECU* | Hmel204017 | 2115425 | 28 | 163.27 | 7507 | 0.002 | 2109574 | 11 | 77.18 | 15879 | 0.005 |
| *H. m. malleti COL* | Hmel204017 | 2119953 | 19 | 540.47 | 2674 | 0.001 | 1958384 | 14 | 99.86 | 14472 | 0.004 |
| *H. m. malleti ECU* | Hmel204017 | 1940231 | 23 | 406.94 | 3468 | 0.001 | 1955983 | 10 | 149.31 | 9451 | 0.003 |
| *H. m. melpomene COL* | Hmel204017 | 1940194 | 21 | 490.45 | 3550 | 0.001 | 2108911 | 1 | 111.29 | 15644 | 0.003 |
| *H. m. melpomene FG* | Hmel204017 | 2070510 | 17 | 1052.51 | 1326 | 0 | 2054159 | 6 | 159.26 | 8763 | 0.002 |
| *H. m. melpomene PAN* | Hmel204017 | 1958471 | 37 | 102.29 | 13645 | 0.004 | 1958371 | 37 | 101.91 | 13696 | 0.004 |
| *H. m. meriana* | Hmel204017 | 1938159 | 36 | 314.03 | 1732 | 0.001 | 2054221 | 4 | 67.64 | 8043 | 0.005 |
| *H. m. nanna NORTH* | Hmel204017 | 1962630 | 17 | 206.08 | 5183 | 0.002 | 2110187 | 1 | 104.45 | 10226 | 0.004 |
| *H. m. nanna SOUTH* | Hmel204017 | 1987449 | 23 | 85.41 | 12506 | 0.004 | 1959797 | 20 | 30.51 | 35006 | 0.012 |
| *H. m. plesseni* | Hmel204017 | 1960131 | 24 | 138 | 10351 | 0.003 | 1958431 | 14 | 107.91 | 13237 | 0.004 |
| *H. m. rosina* | Hmel204017 | 2070277 | 19 | 565.94 | 1716 | 0.001 | 2197537 | 1 | 191.2 | 5078 | 0.002 |
| *H. m. vicina* | Hmel204017 | 2028490 | 25 | 250.5 | 5571 | 0.002 | 1957880 | 21 | 69.35 | 20124 | 0.005 |
| *H. m. vulcanus* | Hmel204017 | 1958592 | 45 | 68.73 | 12826 | 0.005 | 1958592 | 45 | 68.73 | 12826 | 0.005 |
| *H. m. xenoclea* | Hmel204017 | 1940108 | 20 | 427.87 | 3013 | 0.001 | 1957909 | 7 | 220.11 | 5858 | 0.002 |
| *H. pachinus* | Hmel204017 | 2115316 | 40 | 102.78 | 11624 | 0.004 | 2109316 | 37 | 44.2 | 27031 | 0.008 |
| *H. t. florencia* | Hmel204017 | 1967332 | 37 | 143.92 | 7590 | 0.003 | 2065692 | 10 | 108.37 | 10080 | 0.003 |
| *H. t. linaresi* | Hmel204017 | 1859422 | 44 | 75.94 | 15138 | 0.005 | 1859622 | 41 | 74.32 | 15468 | 0.005 |
| *H. t. ssp. nov. ECU* | Hmel204017 | 1967446 | 23 | 164.5 | 6944 | 0.002 | 2053904 | 2 | 142.86 | 7996 | 0.003 |
| *H. t. thelxinoe* | Hmel204017 | 1992794 | 13 | 370.4 | 2639 | 0.001 | 2200268 | 0 | 205.27 | 4762 | 0.002 |
| *H. t. timareta f. contigua* | Hmel204017 | 1967333 | 29 | 177.71 | 5673 | 0.002 | 1894174 | 27 | 153.85 | 6553 | 0.002 |
| *H. t. timareta f. timareta* | Hmel204017 | 1967317 | 29 | 139.95 | 7689 | 0.003 | 1967917 | 28 | 129.38 | 8316 | 0.003 |
| *H. t. ssp. nov. COL* | Hmel204017 | 1967382 | 31 | 129.08 | 7875 | 0.003 | 2066095 | 26 | 67.81 | 14992 | 0.005 |
| *H. besckei* | Hmel206006 | 623821 | 32 | 96.67 | 2936 | 0.003 | 446344 | 6 | 78.83 | 3600 | 0.003 |
| *H. c. chioneus* | Hmel206006 | 552281 | 35 | 206.11 | 4284 | 0.001 | 635186 | 13 | 106.84 | 8264 | 0.003 |
| *H. c. cydnides* | Hmel206006 | 621246 | 49 | 185.05 | 4819 | 0.001 | 636047 | 17 | 90.71 | 9830 | 0.003 |
| *H. c. weymeri gustavi* | Hmel206006 | 551591 | 24 | 285.25 | 2877 | 0.001 | 635597 | 6 | 141.7 | 5793 | 0.002 |
| *H. c. weymeri weymeri* | Hmel206006 | 358310 | 38 | 313.61 | 2521 | 0.001 | 602687 | 4 | 211.68 | 3735 | 0.001 |

| Population | Scaffold | Position | CLR | $\alpha$ | $2N_es$ | $s$ | Position ($\alpha_{min}$) | CLR ($\alpha_{min}$) | $\alpha_{min}$ | $2N_es$ ($\alpha_{min}$) | $s$ ($\alpha_{min}$) |
|---|---|---|---|---|---|---|---|---|---|---|---|
| *H. c. zelinde* | Hmel206006 | 625531 | 20 | 322.96 | 2516 | 0.001 | 547874 | 12 | 195.33 | 4160 | 0.001 |
| *H. elevatus Ecuador* | Hmel206006 | 625339 | 67 | 165.96 | 7938 | 0.002 | 625089 | 56 | 158.23 | 8326 | 0.002 |
| *H. heurippa* | Hmel206006 | 692174 | 209 | 68.12 | 8410 | 0.004 | 622815 | 113 | 56 | 10232 | 0.005 |
| *H. m. amaryllis* | Hmel206006 | 792036 | 13 | 6498.34 | 199 | 0 | 602713 | 3 | 266.71 | 4857 | 0.001 |
| *H. m. cythera* | Hmel206006 | 327697 | 35 | 313.66 | 2881 | 0.001 | 582730 | 1 | 294.86 | 3064 | 0.001 |
| *H. m. ECU* | Hmel206006 | 754627 | 18 | 1234.34 | 843 | 0 | 604468 | 6 | 289.65 | 3594 | 0.001 |
| *H. m. malleti COL* | Hmel206006 | 519734 | 18 | 772.31 | 1331 | 0 | 330318 | 6 | 330.27 | 3112 | 0.001 |
| *H. m. malleti ECU* | Hmel206006 | 520036 | 23 | 608.97 | 2203 | 0 | 604299 | 11 | 168.4 | 7967 | 0.002 |
| *H. m. melpomene COL* | Hmel206006 | 435316 | 77 | 286.56 | 3472 | 0.001 | 635727 | 14 | 142 | 7006 | 0.002 |
| *H. m. melpomene FG* | Hmel206006 | 748824 | 30 | 400.07 | 1622 | 0.001 | 603065 | 21 | 82.02 | 7913 | 0.003 |
| *H. m. melpomene PAN* | Hmel206006 | 550698 | 67 | 79.7 | 12387 | 0.003 | 550848 | 42 | 79.35 | 12441 | 0.003 |
| *H. m. meriana* | Hmel206006 | 398903 | 43 | 178.98 | 2537 | 0.001 | 399353 | 32 | 168.46 | 2696 | 0.002 |
| *H. m. nanna NORTH* | Hmel206006 | 624824 | 52 | 71.95 | 11590 | 0.004 | 624874 | 52 | 71.88 | 11601 | 0.004 |
| *H. m. nanna SOUTH* | Hmel206006 | 697150 | 29 | 65.89 | 12655 | 0.004 | 593689 | 20 | 55.63 | 14990 | 0.005 |
| *H. m. plesseni* | Hmel206006 | 542803 | 28 | 533.96 | 1896 | 0.001 | 604409 | 9 | 195.65 | 5174 | 0.001 |
| *H. m. rosina* | Hmel206006 | 435387 | 31 | 485.67 | 1638 | 0.001 | 485890 | 11 | 311.11 | 2558 | 0.001 |
| *H. m. vicina* | Hmel206006 | 596917 | 19 | 304.47 | 3268 | 0.001 | 635572 | 8 | 102.6 | 9697 | 0.003 |
| *H. m. vulcanus* | Hmel206006 | 620645 | 58 | 160.28 | 4176 | 0.002 | 635647 | 13 | 137.72 | 4860 | 0.002 |
| *H. m. xenoclea* | Hmel206006 | 602412 | 24 | 189.92 | 5286 | 0.001 | 602712 | 20 | 173.66 | 5781 | 0.002 |
| *H. pachinus* | Hmel206006 | 602775 | 56 | 87.16 | 9762 | 0.003 | 603575 | 53 | 82.35 | 10333 | 0.003 |
| *H. t. florencia* | Hmel206006 | 619768 | 71 | 144.46 | 5701 | 0.002 | 603667 | 7 | 91.83 | 8967 | 0.003 |
| *H. t. linaresi* | Hmel206006 | 647619 | 44 | 266.93 | 2946 | 0.001 | 647569 | 41 | 266.8 | 2947 | 0.001 |
| *H. t. ssp. nov. ECU* | Hmel206006 | 622609 | 76 | 118.17 | 7126 | 0.002 | 604357 | 24 | 103.87 | 8107 | 0.003 |
| *H. t. thelxinoe* | Hmel206006 | 619714 | 48 | 198.45 | 3860 | 0.001 | 619764 | 47 | 197.9 | 3871 | 0.001 |
| *H. t. timareta f. contigua* | Hmel206006 | 739720 | 27 | 395.99 | 1932 | 0.001 | 736870 | 5 | 207.58 | 3685 | 0.001 |
| *H. t. timareta f. timareta* | Hmel206006 | 739820 | 24 | 384.53 | 2082 | 0.001 | 503640 | 7 | 342.81 | 2335 | 0.001 |
| *H. t. ssp. nov. COL* | Hmel206006 | 600649 | 106 | 54.24 | 12566 | 0.005 | 599899 | 105 | 53.68 | 12697 | 0.005 |
| *H. besckei* | Hmel208051 | 754245 | 17 | 64.59 | 5134 | 0.006 | 755845 | 11 | 47.33 | 7006 | 0.008 |
| *H. c. chioneus* | Hmel208051 | 1042680 | 28 | 137.66 | 8279 | 0.003 | 911421 | 19 | 66.71 | 17083 | 0.006 |
| *H. c. cydnides* | Hmel208051 | 1043501 | 33 | 101.13 | 10528 | 0.004 | 1043451 | 30 | 99.58 | 10692 | 0.004 |
| *H. c. weymeri gustavi* | Hmel208051 | 1039926 | 62 | 59.32 | 17382 | 0.007 | 1040376 | 4 | 55.87 | 18455 | 0.007 |
| *H. c. weymeri weymeri* | Hmel208051 | 1040038 | 29 | 146.11 | 6744 | 0.003 | 638197 | 13 | 123.9 | 7953 | 0.003 |
| *H. c. zelinde* | Hmel208051 | 1042582 | 23 | 171.2 | 6554 | 0.002 | 911322 | 8 | 87.76 | 12785 | 0.005 |
| *H. elevatus Ecuador* | Hmel208051 | 1044514 | 68 | 86.19 | 18877 | 0.005 | 929499 | 23 | 61.66 | 26386 | 0.007 |
| *H. heurippa* | Hmel208051 | 982755 | 447 | 11.85 | 63379 | 0.033 | 984105 | 268 | 11.46 | 65530 | 0.034 |
| *H. m. amaryllis* | Hmel208051 | 1048965 | 19 | 674.01 | 2446 | 0.001 | 1045265 | 13 | 186.14 | 8856 | 0.002 |
| *H. m. cythera* | Hmel208051 | 846537 | 27 | 395.77 | 3243 | 0.001 | 955998 | 25 | 117.17 | 10955 | 0.003 |
| *H. m. ECU* | Hmel208051 | 1024299 | 40 | 82.15 | 16900 | 0.005 | 1024299 | 40 | 82.15 | 16900 | 0.005 |
| *H. m. malleti COL* | Hmel208051 | 1049946 | 19 | 183.34 | 7932 | 0.002 | 1068899 | 14 | 182.57 | 7965 | 0.002 |
| *H. m. malleti ECU* | Hmel208051 | 1050196 | 14 | 908.1 | 1964 | 0 | 1023894 | 2 | 115.65 | 15426 | 0.004 |
| *H. m. melpomene COL* | Hmel208051 | 1031498 | 23 | 184.04 | 7475 | 0.002 | 745420 | 4 | 161.2 | 8534 | 0.003 |
| *H. m. melpomene FG* | Hmel208051 | 1022462 | 13 | 136.64 | 6828 | 0.003 | 1023112 | 8 | 118.53 | 7871 | 0.003 |
| *H. m. melpomene PAN* | Hmel208051 | 992145 | 28 | 374.68 | 3580 | 0.001 | 905289 | 12 | 124.84 | 10743 | 0.003 |

| Population | Scaffold | Position | CLR | $\alpha$ | $2N_e s$ | $s$ | Position ($\alpha_{min}$) | CLR ($\alpha_{min}$) | $\alpha_{min}$ | $2N_e s$ ($\alpha_{min}$) | $s$ ($\alpha_{min}$) |
|---|---|---|---|---|---|---|---|---|---|---|---|
| *H. m. meriana* | Hmel208051 | 1024688 | 41 | 84.4 | 8036 | 0.005 | 1053241 | 34 | 77.92 | 8705 | 0.005 |
| *H. m. nanna NORTH* | Hmel208051 | 1039701 | 268 | 15.12 | 71234 | 0.026 | 1042701 | 165 | 14.81 | 72711 | 0.027 |
| *H. m. nanna SOUTH* | Hmel208051 | 1073274 | 25 | 40.14 | 26835 | 0.01 | 761403 | 14 | 18.38 | 58609 | 0.022 |
| *H. m. plesseni* | Hmel208051 | 1049999 | 19 | 230.55 | 6532 | 0.002 | 1023996 | 18 | 91.79 | 16404 | 0.004 |
| *H. m. rosina* | Hmel208051 | 903527 | 24 | 205.05 | 4694 | 0.002 | 904777 | 6 | 132.63 | 7256 | 0.003 |
| *H. m. vicina* | Hmel208051 | 1053006 | 45 | 48.28 | 28493 | 0.008 | 1053256 | 43 | 47.58 | 28915 | 0.009 |
| *H. m. vulcanus* | Hmel208051 | 986477 | 43 | 59.01 | 14524 | 0.007 | 986427 | 43 | 58.93 | 14544 | 0.007 |
| *H. m. xenoclea* | Hmel208051 | 1049965 | 26 | 185.01 | 7211 | 0.002 | 910549 | 4 | 86.3 | 15459 | 0.005 |
| *H. pachinus* | Hmel208051 | 957473 | 52 | 62.25 | 17632 | 0.006 | 957473 | 52 | 62.25 | 17632 | 0.006 |
| *H. t. florencia* | Hmel208051 | 1044370 | 163 | 27.82 | 36817 | 0.014 | 1034118 | 78 | 25.36 | 40393 | 0.016 |
| *H. t. linaresi* | Hmel208051 | 1053301 | 78 | 61.85 | 16854 | 0.006 | 1048751 | 37 | 58.96 | 17679 | 0.007 |
| *H. t. ssp. nov. ECU* | Hmel208051 | 1045877 | 144 | 29.56 | 35705 | 0.014 | 1046027 | 73 | 29.51 | 35764 | 0.014 |
| *H. t. thelxinoe* | Hmel208051 | 909967 | 59 | 61.29 | 14753 | 0.006 | 910717 | 55 | 51.43 | 17581 | 0.008 |
| *H. t. timareta f. contigua* | Hmel208051 | 1045354 | 83 | 42.15 | 21556 | 0.009 | 1045254 | 82 | 42.07 | 21597 | 0.009 |
| *H. t. timareta f. timareta* | Hmel208051 | 1043033 | 87 | 34.74 | 28376 | 0.011 | 1042433 | 35 | 34.64 | 28465 | 0.011 |
| *H. t. ssp. nov. COL* | Hmel208051 | 1024784 | 151 | 23.07 | 40454 | 0.017 | 1031634 | 97 | 22.06 | 42310 | 0.018 |
| *H. besckei* | Hmel219003 | 5392785 | 10 | 250.9 | 995 | 0.001 | 5656167 | 7 | 239.49 | 1043 | 0.001 |
| *H. c. chioneus* | Hmel219003 | 5569780 | 86 | 88.48 | 8060 | 0.003 | 5569830 | 86 | 88.47 | 8061 | 0.003 |
| *H. c. cydnides* | Hmel219003 | 5396768 | 123 | 85.52 | 7800 | 0.003 | 5396818 | 123 | 85.49 | 7803 | 0.003 |
| *H. c. weymeri gustavi* | Hmel219003 | 5538911 | 101 | 112.36 | 5544 | 0.002 | 5286426 | 32 | 98.65 | 6314 | 0.002 |
| *H. c. weymeri weymeri* | Hmel219003 | 5553535 | 283 | 38.09 | 15583 | 0.006 | 5554435 | 278 | 37.93 | 15647 | 0.006 |
| *H. c. zelinde* | Hmel219003 | 5570079 | 125 | 62.53 | 11161 | 0.004 | 5570329 | 82 | 62.3 | 11203 | 0.004 |
| *H. elevatus Ecuador* | Hmel219003 | 5568930 | 92 | 252.97 | 4436 | 0.001 | 5576831 | 37 | 143.22 | 7835 | 0.002 |
| *H. heurippa* | Hmel219003 | 5611280 | 129 | 142.67 | 2523 | 0.002 | 5428619 | 118 | 91.69 | 3926 | 0.002 |
| *H. m. amaryllis* | Hmel219003 | 5555079 | 39 | 336.76 | 3004 | 0.001 | 5254364 | 7 | 224.1 | 4514 | 0.001 |
| *H. m. cythera* | Hmel219003 | 5246608 | 277 | 39.17 | 19565 | 0.006 | 5245458 | 210 | 38.8 | 19754 | 0.006 |
| *H. m. ECU* | Hmel219003 | 5576937 | 87 | 80.03 | 11156 | 0.003 | 5577787 | 84 | 78.91 | 11315 | 0.003 |
| *H. m. malleti COL* | Hmel219003 | 5252414 | 33 | 289.02 | 2908 | 0.001 | 5254464 | 30 | 157.13 | 5350 | 0.002 |
| *H. m. malleti ECU* | Hmel219003 | 5567800 | 50 | 384.41 | 2743 | 0.001 | 5576601 | 42 | 116.57 | 9045 | 0.002 |
| *H. m. melpomene COL* | Hmel219003 | 5588298 | 31 | 788.28 | 1086 | 0 | 5286460 | 2 | 201.48 | 4249 | 0.001 |
| *H. m. melpomene FG* | Hmel219003 | 5554226 | 57 | 154.81 | 3433 | 0.002 | 5554426 | 54 | 154.6 | 3438 | 0.002 |
| *H. m. melpomene PAN* | Hmel219003 | 5552111 | 99 | 99.07 | 8155 | 0.002 | 5254367 | 91 | 95 | 8505 | 0.003 |
| *H. m. meriana* | Hmel219003 | 5475294 | 51 | 322.83 | 1212 | 0.001 | 5244725 | 34 | 142.27 | 2750 | 0.002 |
| *H. m. nanna NORTH* | Hmel219003 | 5576562 | 243 | 25.98 | 25617 | 0.009 | 5573512 | 121 | 25.67 | 25933 | 0.009 |
| *H. m. nanna SOUTH* | Hmel219003 | 5597249 | 88 | 51.91 | 12823 | 0.005 | 5607750 | 49 | 42.09 | 15816 | 0.006 |
| *H. m. plesseni* | Hmel219003 | 5254487 | 56 | 99.24 | 8569 | 0.002 | 5254537 | 56 | 99.09 | 8582 | 0.002 |
| *H. m. rosina* | Hmel219003 | 5551670 | 76 | 138.55 | 4565 | 0.002 | 5255279 | 55 | 120.95 | 5229 | 0.002 |
| *H. m. vicina* | Hmel219003 | 5550947 | 50 | 111.2 | 7699 | 0.002 | 5551797 | 47 | 107.56 | 7960 | 0.002 |
| *H. m. vulcanus* | Hmel219003 | 5553765 | 71 | 200.24 | 2712 | 0.001 | 5577768 | 39 | 135.95 | 3994 | 0.002 |
| *H. m. xenoclea* | Hmel219003 | 5576910 | 52 | 59 | 13791 | 0.004 | 5577810 | 49 | 57.61 | 14123 | 0.004 |
| *H. pachinus* | Hmel219003 | 5551837 | 138 | 78.62 | 8787 | 0.003 | 5555187 | 133 | 66.15 | 10443 | 0.004 |
| *H. t. florencia* | Hmel219003 | 5715417 | 155 | 86.73 | 6303 | 0.003 | 5251087 | 83 | 44.97 | 12156 | 0.005 |
| *H. t. linaresi* | Hmel219003 | 5481153 | 128 | 135.29 | 3870 | 0.002 | 5388590 | 90 | 77.48 | 6757 | 0.003 |

| Population | Scaffold | Position | CLR | $\alpha$ | $2N_es$ | $s$ | Position ($\alpha_{min}$) | CLR ($\alpha_{min}$) | $\alpha_{min}$ | $2N_es$ ($\alpha_{min}$) | $s$ ($\alpha_{min}$) |
|---|---|---|---|---|---|---|---|---|---|---|---|
| *H. t. ssp. nov. ECU* | Hmel219003 | 5715333 | 177 | 78.73 | 7015 | 0.003 | 5251188 | 57 | 59.79 | 9238 | 0.004 |
| *H. t. thelxinoe* | Hmel219003 | 5611298 | 177 | 88.47 | 4333 | 0.003 | 5611048 | 161 | 88.12 | 4350 | 0.003 |
| *H. t. timareta f. contigua* | Hmel219003 | 5250567 | 242 | 28.08 | 16568 | 0.008 | 5250067 | 200 | 27.78 | 16749 | 0.008 |
| *H. t. timareta f. timareta* | Hmel219003 | 5551811 | 161 | 75.74 | 6270 | 0.003 | 5251121 | 70 | 59.27 | 8012 | 0.004 |
| *H. t. ssp. nov. COL* | Hmel219003 | 5583888 | 553 | 18.15 | 24628 | 0.013 | 5575787 | 420 | 16.8 | 26600 | 0.014 |

**Supplementary Table 4:** Position, composite likelihood-ratio statistics (CLR) and strength of selection ($\alpha$, $2N_es$, and $s$) for the highest CLR and the smallest $\alpha$ value on each colour pattern scaffold ($\alpha_{min}$) for the *H. melpomene*-clade. Additional relevant peaks on scaffolds are also given. Data are from SweepFinder2[3,4] runs with background site frequency spectrum estimated from background and colour pattern scaffolds.

| Population | Locus | Scaffold | Position | CLR | $\alpha$ | $2N_es$ | $s$ | Position ($\alpha_{min}$) | CLR ($\alpha_{min}$) | $\alpha_{min}$ | $2N_es$ ($\alpha_{min}$) | $s$ ($\alpha_{min}$) |
|---|---|---|---|---|---|---|---|---|---|---|---|---|
| *H. besckei* | *aristaless* | Hmel201011 | 2488332 | 15 | 245.38 | 844 | 0.001 | 2623264 | 8 | 78.98 | 2621 | 0.003 |
| *H. c. chioneus* | *aristaless* | Hmel201011 | 2637797 | 28 | 162.3 | 3603 | 0.001 | 2637697 | 27 | 157.39 | 3716 | 0.001 |
| *H. c. cydnides* | *aristaless* | Hmel201011 | 2613872 | 82 | 70.24 | 7222 | 0.003 | 2613872 | 82 | 70.24 | 7222 | 0.003 |
| *H. c. weymeri gustavi* | *aristaless* | Hmel201011 | 2598794 | 52 | 129.02 | 4032 | 0.002 | 2600094 | 27 | 100.1 | 5197 | 0.002 |
| *H. c. weymeri weymeri* | *aristaless* | Hmel201011 | 2595343 | 95 | 87.66 | 5334 | 0.003 | 2600144 | 37 | 62.72 | 7455 | 0.004 |
| *H. c. zelinde* | *aristaless* | Hmel201011 | 2637640 | 28 | 163.96 | 3783 | 0.001 | 2637640 | 28 | 163.96 | 3783 | 0.001 |
| *H. elevatus Ecuador* | *aristaless* | Hmel201011 | 2671769 | 42 | 162.82 | 5641 | 0.001 | 2671769 | 42 | 162.82 | 5641 | 0.001 |
| *H. heurippa* | *aristaless* | Hmel201011 | 2648749 | 85 | 50.23 | 6880 | 0.005 | 2647699 | 62 | 46.58 | 7420 | 0.005 |
| *H. m. amaryllis* | *aristaless* | Hmel201011 | 2538415 | 9 | 652817.2 | 1 | 0 | 2740662 | 4 | 338.7 | 2546 | 0.001 |
| *H. m. cythera* | *aristaless* | Hmel201011 | 2616846 | 54 | 91.95 | 6473 | 0.003 | 2623397 | 19 | 49.58 | 12004 | 0.005 |
| *H. m. ECU* | *aristaless* | Hmel201011 | 2642217 | 21 | 353.41 | 2165 | 0.001 | 2641617 | 20 | 247.76 | 3088 | 0.001 |
| *H. m. malleti COL* | *aristaless* | Hmel201011 | 2563116 | 26 | 471.86 | 1562 | 0.001 | 2725444 | 9 | 233.21 | 3160 | 0.001 |
| *H. m. malleti ECU* | *aristaless* | Hmel201011 | 2643054 | 19 | 657.64 | 1383 | 0 | 2740666 | 0 | 291.95 | 3116 | 0.001 |
| *H. m. melpomene COL* | *aristaless* | Hmel201011 | 2623275 | 116 | 43.1 | 17319 | 0.006 | 2623525 | 112 | 42.4 | 17604 | 0.006 |
| *H. m. melpomene FG* | *aristaless* | Hmel201011 | 2518352 | 23 | 521.78 | 1016 | 0 | 2583859 | 3 | 216.89 | 2444 | 0.001 |
| *H. m. melpomene PAN* | *aristaless* | Hmel201011 | 2601790 | 31 | 317.04 | 2092 | 0.001 | 2600840 | 20 | 155.34 | 4271 | 0.002 |
| *H. m. meriana* | *aristaless* | Hmel201011 | 2584942 | 15 | 1029.57 | 371 | 0 | 2483534 | 3 | 275.37 | 1387 | 0.001 |
| *H. m. nanna NORTH* | *aristaless* | Hmel201011 | 2542627 | 29 | 206 | 2855 | 0.001 | 2545678 | 10 | 196.89 | 2987 | 0.001 |
| *H. m. nanna SOUTH* | *aristaless* | Hmel201011 | 2635733 | 24 | 77.1 | 7628 | 0.003 | 2636533 | 21 | 75.23 | 7818 | 0.003 |
| *H. m. plesseni* | *aristaless* | Hmel201011 | 2716853 | 11 | 667.16 | 1083 | 0 | 2740655 | 1 | 325.36 | 2220 | 0.001 |
| *H. m. rosina* | *aristaless* | Hmel201011 | 2648668 | 40 | 192.06 | 2763 | 0.001 | 2661869 | 15 | 120.39 | 4408 | 0.002 |
| *H. m. vicina* | *aristaless* | Hmel201011 | 2643104 | 19 | 266.38 | 2802 | 0.001 | 2641554 | 16 | 133.94 | 5573 | 0.002 |
| *H. m. vulcanus* | *aristaless* | Hmel201011 | 2669653 | 31 | 319.09 | 1526 | 0.001 | 2661802 | 4 | 236.26 | 2061 | 0.001 |
| *H. m. xenoclea* | *aristaless* | Hmel201011 | 2643390 | 16 | 981.57 | 692 | 0 | 2641640 | 7 | 250.65 | 2708 | 0.001 |
| *H. pachinus* | *aristaless* | Hmel201011 | 2637780 | 52 | 158.95 | 3419 | 0.001 | 2703291 | 40 | 68.95 | 7882 | 0.003 |
| *H. t. florencia* | *aristaless* | Hmel201011 | 2640195 | 141 | 42.99 | 12804 | 0.005 | 2669300 | 130 | 27.3 | 20158 | 0.009 |
| | | | 2669550 | 134 | 27.48 | 20032 | 0.009 | 2669300 | 130 | 27.3 | 20158 | 0.009 |
| *H. t. linaresi* | *aristaless* | Hmel201011 | 2634430 | 114 | 31.16 | 15805 | 0.007 | 2634180 | 62 | 30.95 | 15915 | 0.008 |
| *H. t. ssp. nov. ECU* | *aristaless* | Hmel201011 | 2666249 | 101 | 49.94 | 11611 | 0.005 | 2664899 | 79 | 44.49 | 13033 | 0.005 |
| *H. t. thelxinoe* | *aristaless* | Hmel201011 | 2674221 | 51 | 113.09 | 4396 | 0.002 | 2631467 | 36 | 108.05 | 4601 | 0.002 |
| *H. t. timareta f. contigua* | *aristaless* | Hmel201011 | 2674055 | 105 | 44.53 | 11810 | 0.005 | 2673705 | 102 | 43.95 | 11966 | 0.005 |
| *H. t. timareta f. timareta* | *aristaless* | Hmel201011 | 2674126 | 53 | 69.97 | 8183 | 0.003 | 2630917 | 34 | 54.97 | 10415 | 0.004 |
| *H. t. ssp. nov. COL* | *aristaless* | Hmel201011 | 2669729 | 110 | 36.41 | 12465 | 0.006 | 2672229 | 81 | 34.16 | 13286 | 0.007 |
| *H. besckei* | *WntA* | Hmel210004 | 1559377 | 28 | 110.61 | 1947 | 0.002 | 1566878 | 21 | 40.39 | 5331 | 0.005 |
| *H. c. chioneus* | *WntA* | Hmel210004 | 1620799 | 111 | 34.86 | 15432 | 0.006 | 1626500 | 70 | 32.03 | 16795 | 0.007 |
| *H. c. cydnides* | *WntA* | Hmel210004 | 1811792 | 134 | 31.47 | 18160 | 0.007 | 1817892 | 101 | 25.37 | 22521 | 0.009 |

| Population | Locus | Scaffold | Position | CLR | $\alpha$ | $2N_es$ | $s$ | Position ($\alpha_{min}$) | CLR ($\alpha_{min}$) | $\alpha_{min}$ | $2N_es$ ($\alpha_{min}$) | $s$ ($\alpha_{min}$) |
|---|---|---|---|---|---|---|---|---|---|---|---|---|
| *H. c. weymeri gustavi* | *WntA* | Hmel210004 | 1806398 | 257 | 16.57 | 32081 | 0.013 | 1805198 | 251 | 16.18 | 32860 | 0.014 |
| *H. c. weymeri weymeri* | *WntA* | Hmel210004 | 1810315 | 166 | 22.13 | 24964 | 0.01 | 1811415 | 151 | 21.65 | 25519 | 0.01 |
| *H. c. zelinde* | *WntA* | Hmel210004 | 1620832 | 95 | 36.08 | 16055 | 0.006 | 1620482 | 87 | 35.61 | 16269 | 0.006 |
| *H. elevatus ECU* | *WntA* | Hmel210004 | 1828331 | 247 | 29.65 | 28873 | 0.008 | 1827831 | 239 | 29.18 | 29337 | 0.008 |
| *H. heurippa* | *WntA* | Hmel210004 | 1560284 | 186 | 24.93 | 13503 | 0.009 | 1567234 | 179 | 23.17 | 14532 | 0.009 |
| *H. m. amaryllis* | *WntA* | Hmel210004 | 1809664 | 121 | 67.34 | 11727 | 0.003 | 1810164 | 104 | 58.95 | 13396 | 0.004 |
| *H. m. cythera* | *WntA* | Hmel210004 | 1811174 | 96 | 44.81 | 12915 | 0.005 | 1811424 | 94 | 44.32 | 13059 | 0.005 |
| *H. m. ecuadoriensis* | *WntA* | Hmel210004 | 1848718 | 232 | 38.28 | 17577 | 0.006 | 1621560 | 132 | 30.73 | 21896 | 0.007 |
| | | | 1808717 | 139 | 34.32 | 19609 | 0.007 | 1810117 | 85 | 33.01 | 20386 | 0.007 |
| *H. m. malleti COL* | *WntA* | Hmel210004 | 1621626 | 164 | 37.02 | 18699 | 0.006 | 1626276 | 20 | 33.64 | 20578 | 0.007 |
| *H. m. malleti ECU* | *WntA* | Hmel210004 | 1809624 | 238 | 34.8 | 24145 | 0.007 | 1810125 | 234 | 34.02 | 24694 | 0.007 |
| *H. m. melpomene COL* | *WntA* | Hmel210004 | 1631681 | 334 | 14.47 | 50043 | 0.016 | 1631731 | 333 | 14.47 | 50070 | 0.016 |
| *H. m. melpomene FG* | *WntA* | Hmel210004 | 1568298 | 63 | 25.08 | 16668 | 0.009 | 1567398 | 59 | 24.61 | 16987 | 0.009 |
| *H. m. melpomene PAN* | *WntA* | Hmel210004 | 1629713 | 314 | 16.29 | 39705 | 0.014 | 1629363 | 279 | 16.23 | 39842 | 0.014 |
| *H. m. meriana* | *WntA* | Hmel210004 | 1855774 | 88 | 105.24 | 3200 | 0.002 | 1623664 | 61 | 54.24 | 6209 | 0.004 |
| *H. m. nanna NORTH* | *WntA* | Hmel210004 | 1806151 | 99 | 17.3 | 31418 | 0.013 | 1816252 | 65 | 16.76 | 32425 | 0.013 |
| *H. m. nanna SOUTH* | *WntA* | Hmel210004 | 1783553 | 38 | 19.94 | 27254 | 0.011 | 1782353 | 8 | 19.78 | 27473 | 0.011 |
| *H. m. plesseni* | *WntA* | Hmel210004 | 1851807 | 649 | 9.5 | 63153 | 0.023 | 1828905 | 631 | 8.18 | 73376 | 0.027 |
| *H. m. rosina* | *WntA* | Hmel210004 | 1567842 | 123 | 29.7 | 15534 | 0.007 | 1568692 | 73 | 29.45 | 15666 | 0.007 |
| *H. m. vicina* | *WntA* | Hmel210004 | 1622166 | 125 | 33.27 | 21769 | 0.007 | 1620716 | 98 | 31.37 | 23090 | 0.007 |
| *H. m. vulcanus* | *WntA* | Hmel210004 | 1625201 | 132 | 23.5 | 17282 | 0.009 | 1626051 | 101 | 23.15 | 17550 | 0.009 |
| *H. m. xenoclea* | *WntA* | Hmel210004 | 1811380 | 487 | 6.17 | 86938 | 0.036 | 1811780 | 485 | 6.17 | 86959 | 0.036 |
| *H. pachinus* | *WntA* | Hmel210004 | 1810240 | 118 | 20.15 | 26224 | 0.011 | 1805240 | 110 | 17.34 | 30483 | 0.013 |
| *H. t. florencia* | *WntA* | Hmel210004 | 1566624 | 406 | 13.48 | 38127 | 0.016 | 1566424 | 405 | 13.47 | 38158 | 0.016 |
| *H. t. linaresi* | *WntA* | Hmel210004 | 1825099 | 156 | 26.85 | 18128 | 0.008 | 1825199 | 156 | 26.84 | 18132 | 0.008 |
| *H. t. ssp. nov. ECU* | *WntA* | Hmel210004 | 1571479 | 266 | 24.47 | 19504 | 0.009 | 1567328 | 242 | 20.54 | 23229 | 0.011 |
| *H. t. thelxinoe* | *WntA* | Hmel210004 | 1765448 | 124 | 26.67 | 14929 | 0.008 | 1765348 | 122 | 26.64 | 14946 | 0.008 |
| *H. t. timareta f. contigua* | *WntA* | Hmel210004 | 1573226 | 148 | 27.87 | 13841 | 0.008 | 1567926 | 125 | 21.56 | 17892 | 0.01 |
| *H. t. timareta f. timareta* | *WntA* | Hmel210004 | 1567591 | 197 | 17.02 | 24745 | 0.013 | 1567891 | 197 | 17 | 24774 | 0.013 |
| *H. t. ssp. nov. COL* | *WntA* | Hmel210004 | 1567227 | 303 | 15.57 | 30304 | 0.014 | 1866345 | 169 | 13.17 | 35834 | 0.017 |
| *H. besckei* | *cortex* | Hmel215006 | 662442 | 27 | 58.7 | 5537 | 0.006 | 925499 | 11 | 10.3 | 31569 | 0.032 |
| *H. c. chioneus* | *cortex* | Hmel215006 | 1209730 | 323 | 20.92 | 40839 | 0.017 | 923867 | 7 | 19.71 | 43356 | 0.018 |
| *H. c. cydnides* | *cortex* | Hmel215006 | 1238415 | 167 | 51.71 | 15293 | 0.007 | 923849 | 9 | 19.71 | 40124 | 0.018 |
| *H. c. weymeri gustavi* | *cortex* | Hmel215006 | 1220571 | 444 | 19.58 | 34335 | 0.018 | 1326374 | 34 | 13.15 | 51123 | 0.026 |
| | | | 1329174 | 393 | 13.31 | 50508 | 0.026 | 1326374 | 34 | 13.15 | 51123 | 0.026 |
| *H. c. weymeri weymeri* | *cortex* | Hmel215006 | 1337825 | 1791 | 6.21 | 98669 | 0.056 | 1334724 | 1673 | 6.2 | 98841 | 0.056 |
| | | | 1218021 | 254 | 30.64 | 19992 | 0.011 | 1215371 | 217 | 30.09 | 20361 | 0.011 |
| *H. c. zelinde* | *cortex* | Hmel215006 | 1218639 | 189 | 30.72 | 29434 | 0.012 | 923979 | 36 | 12.76 | 70863 | 0.028 |
| *H. elevatus ECU* | *cortex* | Hmel215006 | 1563763 | 150 | 71.78 | 19358 | 0.005 | 923634 | 6 | 24.91 | 55787 | 0.015 |
| *H. heurippa* | *cortex* | Hmel215006 | 1447077 | 482 | 20.11 | 22385 | 0.017 | 924905 | 146 | 8.93 | 50418 | 0.038 |
| *H. m. amaryllis* | *cortex* | Hmel215006 | 1575917 | 453 | 13.88 | 57985 | 0.025 | 1581267 | 296 | 12.47 | 64578 | 0.028 |
| *H. m. cythera* | *cortex* | Hmel215006 | 1105740 | 828 | 5.43 | 126643 | 0.064 | 1094539 | 588 | 5.04 | 136563 | 0.069 |
| | | | 1231296 | 789 | 10.23 | 67256 | 0.034 | 1227646 | 592 | 10.01 | 68720 | 0.035 |

| Population | Locus | Scaffold | Position | CLR | $\alpha$ | $2N_es$ | $s$ | Position ($\alpha_{min}$) | CLR ($\alpha_{min}$) | $\alpha_{min}$ | $2N_es$ ($\alpha_{min}$) | $s$ ($\alpha_{min}$) |
|---|---|---|---|---|---|---|---|---|---|---|---|---|
| *H. m. ecuadoriensis* | *cortex* | Hmel215006 | 1459529 | 187 | 38.58 | 24655 | 0.009 | 923556 | 49 | 13.77 | 69097 | 0.026 |
| *H. m. malleti COL* | *cortex* | Hmel215006 | 1105390 | 237 | 19.09 | 51609 | 0.019 | 924031 | 51 | 12.91 | 76296 | 0.028 |
| *H. m. malleti ECU* | *cortex* | Hmel215006 | 1098573 | 150 | 40.38 | 29953 | 0.009 | 924964 | 73 | 11.52 | 104998 | 0.031 |
| *H. m. melpomene COL* | *cortex* | Hmel215006 | 1095016 | 413 | 15.2 | 58430 | 0.023 | 1095166 | 411 | 15.19 | 58466 | 0.023 |
| *H. m. melpomene FG* | *cortex* | Hmel215006 | 1557309 | 356 | 22.33 | 30582 | 0.016 | 827794 | 1 | 17.85 | 38264 | 0.019 |
| *H. m. melpomene PAN* | *cortex* | Hmel215006 | 1096966 | 434 | 8.02 | 109921 | 0.044 | 1093116 | 286 | 7.68 | 114713 | 0.046 |
| *H. m. meriana* | *cortex* | Hmel215006 | 1220357 | 193 | 37.36 | 13704 | 0.009 | 921392 | 18 | 25.52 | 20061 | 0.013 |
| *H. m. nanna NORTH* | *cortex* | Hmel215006 | 1460407 | 772 | 11.55 | 63590 | 0.03 | 1065699 | 408 | 4.68 | 157001 | 0.075 |
| | | | 1226102 | 655 | 7.43 | 98857 | 0.047 | 1225352 | 600 | 7.41 | 99216 | 0.047 |
| | | | 1062599 | 561 | 4.72 | 155677 | 0.074 | 1065699 | 408 | 4.68 | 157001 | 0.075 |
| | | | 1578060 | 619 | 10.59 | 69363 | 0.033 | 1574709 | 411 | 10.42 | 70512 | 0.034 |
| *H. m. nanna SOUTH* | *cortex* | Hmel215006 | 1130468 | 118 | 10.85 | 67707 | 0.032 | 1123618 | 22 | 10.65 | 69004 | 0.033 |
| *H. m. plesseni* | *cortex* | Hmel215006 | 1237065 | 1564 | 6.45 | 114884 | 0.054 | 1236815 | 1563 | 6.44 | 114895 | 0.054 |
| | | | 1366372 | 635 | 26.2 | 28268 | 0.013 | 1366822 | 572 | 26.17 | 28298 | 0.013 |
| | | | 1447976 | 557 | 35.77 | 20702 | 0.01 | 1448876 | 542 | 35.58 | 20809 | 0.01 |
| *H. m. rosina* | *cortex* | Hmel215006 | 1069405 | 378 | 4.56 | 137130 | 0.076 | 1069005 | 338 | 4.55 | 137581 | 0.076 |
| *H. m. vicina* | *cortex* | Hmel215006 | 1396187 | 207 | 30.26 | 29354 | 0.012 | 1094225 | 47 | 11.11 | 79959 | 0.032 |
| *H. m. vulcanus* | *cortex* | Hmel215006 | 1200505 | 383 | 18.62 | 32697 | 0.019 | 1106402 | 51 | 12.84 | 47409 | 0.027 |
| *H. m. xenoclea* | *cortex* | Hmel215006 | 1543692 | 607 | 15.34 | 52052 | 0.023 | 1055866 | 165 | 10.76 | 74197 | 0.033 |
| | | | 1459937 | 563 | 18.7 | 42714 | 0.019 | 1459787 | 563 | 18.7 | 42717 | 0.019 |
| *H. pachinus* | *cortex* | Hmel215006 | 1457620 | 321 | 24.43 | 30316 | 0.014 | 924302 | 41 | 10.62 | 69733 | 0.033 |
| *H. t. florencia* | *cortex* | Hmel215006 | 1060668 | 486 | 10.3 | 73712 | 0.034 | 1060918 | 464 | 10.28 | 73839 | 0.034 |
| *H. t. linaresi* | *cortex* | Hmel215006 | 1328458 | 539 | 14.27 | 49286 | 0.024 | 1329958 | 499 | 14.04 | 50085 | 0.025 |
| *H. t. ssp. nov. ECU* | *cortex* | Hmel215006 | 1056672 | 368 | 10 | 72662 | 0.035 | 1057272 | 221 | 9.92 | 73247 | 0.035 |
| *H. t. thelxinoe* | *cortex* | Hmel215006 | 1245780 | 150 | 74.7 | 8618 | 0.005 | 928117 | 8 | 27.04 | 23811 | 0.013 |
| *H. t. timareta f. contigua* | *cortex* | Hmel215006 | 1457826 | 356 | 25.71 | 25342 | 0.013 | 1094861 | 327 | 11.82 | 55133 | 0.029 |
| *H. t. timareta f. timareta* | *cortex* | Hmel215006 | 1098871 | 412 | 10.51 | 61702 | 0.033 | 1094021 | 383 | 9.85 | 65876 | 0.035 |
| *H. t. ssp. nov. COL* | *cortex* | Hmel215006 | 1062350 | 410 | 8.82 | 72045 | 0.039 | 1069851 | 397 | 8.11 | 78320 | 0.043 |
| *H. besckei* | *optix* | Hmel218003 | 736567 | 27 | 206.69 | 927 | 0.001 | 868677 | 22 | 69.91 | 2741 | 0.003 |
| *H. c. chioneus* | *optix* | Hmel218003 | 804168 | 87 | 125.11 | 4209 | 0.002 | 788617 | 76 | 64.26 | 8194 | 0.003 |
| *H. c. cydnides* | *optix* | Hmel218003 | 637588 | 98 | 63.47 | 7727 | 0.003 | 789494 | 78 | 55.25 | 8877 | 0.004 |
| *H. c. weymeri gustavi* | *optix* | Hmel218003 | 624687 | 119 | 45.62 | 9309 | 0.005 | 624737 | 119 | 45.61 | 9309 | 0.005 |
| *H. c. weymeri weymeri* | *optix* | Hmel218003 | 1019902 | 158 | 123.37 | 2741 | 0.002 | 786278 | 73 | 71.45 | 4733 | 0.003 |
| *H. c. zelinde* | *optix* | Hmel218003 | 789014 | 76 | 64.69 | 8021 | 0.003 | 789414 | 74 | 64.1 | 8094 | 0.003 |
| *H. elevatus ECU* | *optix* | Hmel218003 | 773075 | 146 | 150.85 | 4629 | 0.001 | 857481 | 39 | 51.17 | 13645 | 0.004 |
| *H. heurippa* | *optix* | Hmel218003 | 857779 | 360 | 22.39 | 9401 | 0.009 | 854779 | 306 | 22.17 | 9495 | 0.009 |
| | | | 781223 | 341 | 31.12 | 6763 | 0.007 | 785474 | 300 | 27.81 | 7570 | 0.007 |
| *H. m. amaryllis* | *optix* | Hmel218003 | 786133 | 182 | 59.66 | 7809 | 0.004 | 786483 | 180 | 59.45 | 7837 | 0.004 |
| *H. m. cythera* | *optix* | Hmel218003 | 838195 | 342 | 21.03 | 19087 | 0.01 | 838995 | 251 | 20.98 | 19137 | 0.01 |
| *H. m. ECU* | *optix* | Hmel218003 | 813087 | 130 | 83.53 | 5784 | 0.003 | 810887 | 125 | 73.15 | 6605 | 0.003 |
| *H. m. malleti COL* | *optix* | Hmel218003 | 814383 | 234 | 38.35 | 13047 | 0.006 | 843034 | 74 | 30.45 | 16428 | 0.007 |
| *H. m. malleti ECU* | *optix* | Hmel218003 | 814624 | 197 | 69.18 | 9467 | 0.003 | 842527 | 70 | 40.15 | 16312 | 0.006 |
| *H. m. melpomene COL* | *optix* | Hmel218003 | 672284 | 72 | 192.34 | 2591 | 0.001 | 624731 | 32 | 75.58 | 6593 | 0.003 |

| **Population** | **Locus** | **Scaffold** | **Position** | **CLR** | **$\alpha$** | **$2N_es$** | **$s$** | **Position ($\alpha_{min}$)** | **CLR ($\alpha_{min}$)** | **$\alpha_{min}$** | **$2N_es$ ($\alpha_{min}$)** | **$s$ ($\alpha_{min}$)** |
|---|---|---|---|---|---|---|---|---|---|---|---|---|
| *H. m. melpomene FG* | *optix* | Hmel218003 | 649562 | 81 | 61.99 | 5967 | 0.003 | 648312 | 70 | 60.86 | 6077 | 0.004 |
| *H. m. melpomene PAN* | *optix* | Hmel218003 | 855571 | 378 | 24.29 | 18797 | 0.009 | 841770 | 278 | 20.58 | 22189 | 0.011 |
| *H. m. meriana* | *optix* | Hmel218003 | 801534 | 1085 | 10.59 | 31543 | 0.02 | 811585 | 656 | 10.4 | 32131 | 0.021 |
| *H. m. nanna NORTH* | *optix* | Hmel218003 | 782525 | 191 | 50.12 | 9288 | 0.004 | 859480 | 133 | 38.08 | 12225 | 0.006 |
| *H. m. nanna SOUTH* | *optix* | Hmel218003 | 726487 | 30 | 78.35 | 5942 | 0.003 | 727738 | 28 | 73.94 | 6296 | 0.003 |
| *H. m. plesseni* | *optix* | Hmel218003 | 784931 | 1228 | 9.4 | 31127 | 0.023 | 647624 | 879 | 8.61 | 33962 | 0.025 |
| | | | 640874 | 1199 | 8.72 | 33528 | 0.024 | 647624 | 879 | 8.61 | 33962 | 0.025 |
| | | | 732328 | 673 | 16.42 | 17818 | 0.013 | 729378 | 612 | 16.1 | 18165 | 0.013 |
| *H. m. rosina* | *optix* | Hmel218003 | 847982 | 281 | 20.56 | 16860 | 0.01 | 847682 | 279 | 20.55 | 16871 | 0.01 |
| *H. m. vicina* | *optix* | Hmel218003 | 790899 | 210 | 19.94 | 24988 | 0.011 | 795400 | 106 | 19.06 | 26147 | 0.012 |
| *H. m. vulcanus* | *optix* | Hmel218003 | 848105 | 324 | 14.4 | 19699 | 0.015 | 842254 | 206 | 13.65 | 20781 | 0.016 |
| *H. m. xenoclea* | *optix* | Hmel218003 | 727532 | 519 | 14.82 | 24921 | 0.015 | 728633 | 409 | 14.79 | 24974 | 0.015 |
| *H. pachinus* | *optix* | Hmel218003 | 648265 | 181 | 40.25 | 11029 | 0.005 | 646315 | 86 | 39.49 | 11240 | 0.006 |
| *H. t. florencia* | *optix* | Hmel218003 | 705381 | 164 | 93.56 | 4193 | 0.002 | 679980 | 87 | 70.48 | 5566 | 0.003 |
| *H. t. linaresi* | *optix* | Hmel218003 | 804236 | 288 | 33.26 | 9897 | 0.006 | 802686 | 159 | 32.42 | 10151 | 0.007 |
| *H. t. ssp. nov. ECU* | *optix* | Hmel218003 | 705531 | 142 | 109.22 | 3486 | 0.002 | 671029 | 24 | 71.52 | 5324 | 0.003 |
| *H. t. thelxinoe* | *optix* | Hmel218003 | 542307 | 51 | 385.24 | 843 | 0.001 | 788326 | 46 | 131.68 | 2467 | 0.002 |
| *H. t. timareta f. contigua* | *optix* | Hmel218003 | 868890 | 98 | 68.01 | 4774 | 0.003 | 941544 | 88 | 64.05 | 5070 | 0.003 |
| *H. t. timareta f. timareta* | *optix* | Hmel218003 | 864840 | 103 | 76.84 | 4131 | 0.003 | 864990 | 102 | 76.6 | 4144 | 0.003 |
| *H. t. ssp. nov. COL* | *optix* | Hmel218003 | 597376 | 209 | 45.38 | 6643 | 0.005 | 615177 | 114 | 34.99 | 8616 | 0.006 |

**Supplementary Table 5:** Position, composite likelihood-ratio statistics (CLR) and strength of selection ($\alpha$, $2N_es$, and $s$) for the highest CLR and the smallest $\alpha$ value on each background scaffold ($\alpha_{min}$) for the *H. melpomene*-clade. Data are from SweepFinder2[3,4] runs with background site frequency spectrum estimated from background and colour pattern scaffolds.

| Population | Scaffold | Position | CLR | $\alpha$ | $2N_es$ | $s$ | Position ($\alpha_{min}$) | CLR ($\alpha_{min}$) | $\alpha_{min}$ | $2N_es$ ($\alpha_{min}$) | $s$ ($\alpha_{min}$) |
|---|---|---|---|---|---|---|---|---|---|---|---|
| *H. besckei* | Hmel204017 | 2202171 | 10 | 545.28 | 654 | 0.001 | 2198120 | 7 | 41.71 | 8554 | 0.008 |
| *H. c. chioneus* | Hmel204017 | 2299755 | 19 | 697.25 | 1570 | 0.001 | 2298955 | 16 | 219.31 | 4992 | 0.002 |
| *H. c. cydnides* | Hmel204017 | 2003132 | 31 | 135.83 | 9006 | 0.003 | 2003932 | 22 | 131.74 | 9286 | 0.003 |
| *H. c. weymeri gustavi* | Hmel204017 | 2019892 | 36 | 220.12 | 5376 | 0.002 | 2004140 | 22 | 100.88 | 11731 | 0.004 |
| *H. c. weymeri weymeri* | Hmel204017 | 2016795 | 23 | 228.35 | 4828 | 0.002 | 2053101 | 0 | 96.24 | 11455 | 0.004 |
| *H. c. zelinde* | Hmel204017 | 2255583 | 10 | 533.35 | 2113 | 0.001 | 2117268 | 7 | 292.38 | 3854 | 0.001 |
| *H. elevatus Ecuador* | Hmel204017 | 2119154 | 24 | 437.64 | 3926 | 0.001 | 2110153 | 5 | 84.32 | 20375 | 0.005 |
| *H. heurippa* | Hmel204017 | 2097849 | 76 | 70.77 | 11544 | 0.005 | 2096449 | 71 | 67.12 | 12171 | 0.005 |
| *H. m. amaryllis* | Hmel204017 | 1957699 | 13 | 253.74 | 6119 | 0.001 | 2108612 | 0 | 240.68 | 6451 | 0.002 |
| *H. m. cythera* | Hmel204017 | 2070522 | 17 | 952.37 | 1287 | 0 | 2115325 | 15 | 194.2 | 6311 | 0.002 |
| *H. m. ECU* | Hmel204017 | 2033793 | 16 | 823.66 | 1755 | 0 | 1957884 | 3 | 176.14 | 8205 | 0.002 |
| *H. m. malleti COL* | Hmel204017 | 1940231 | 16 | 451.95 | 3123 | 0.001 | 1957983 | 9 | 195.98 | 7201 | 0.002 |
| *H. m. malleti ECU* | Hmel204017 | 1940094 | 16 | 866.84 | 2008 | 0 | 2107461 | 0 | 258.44 | 6736 | 0.001 |
| *H. m. melpomene COL* | Hmel204017 | 2070510 | 13 | 1260.27 | 1107 | 0 | 2054109 | 1 | 180.77 | 7720 | 0.002 |
| *H. m. melpomene FG* | Hmel204017 | 2003188 | 23 | 140.16 | 5579 | 0.003 | 2001888 | 19 | 92.68 | 8437 | 0.004 |
| *H. m. melpomene PAN* | Hmel204017 | 1958471 | 21 | 127.92 | 10911 | 0.003 | 1958371 | 21 | 127.07 | 10984 | 0.003 |
| *H. m. meriana* | Hmel204017 | 1938159 | 32 | 336.68 | 1616 | 0.001 | 2054271 | 0 | 79.19 | 6871 | 0.004 |
| *H. m. nanna NORTH* | Hmel204017 | 1962630 | 12 | 288.27 | 3705 | 0.001 | 2110887 | 0 | 163.6 | 6529 | 0.002 |
| *H. m. nanna SOUTH* | Hmel204017 | 1987449 | 19 | 95.72 | 11159 | 0.004 | 1959497 | 11 | 49.03 | 21786 | 0.008 |
| *H. m. plesseni* | Hmel204017 | 1960131 | 13 | 192.83 | 7408 | 0.002 | 1960181 | 12 | 190.87 | 7484 | 0.002 |
| *H. m. rosina* | Hmel204017 | 2070277 | 14 | 634.92 | 1529 | 0.001 | 2117181 | 2 | 277.77 | 3495 | 0.001 |
| *H. m. vicina* | Hmel204017 | 2028490 | 19 | 321.68 | 4339 | 0.001 | 1957980 | 9 | 100.14 | 13937 | 0.004 |
| *H. m. vulcanus* | Hmel204017 | 1958592 | 29 | 106.22 | 8299 | 0.003 | 1960792 | 24 | 98.95 | 8909 | 0.004 |
| *H. m. xenoclea* | Hmel204017 | 1940108 | 14 | 607.94 | 2121 | 0.001 | 2004412 | 3 | 329.68 | 3911 | 0.001 |
| *H. pachinus* | Hmel204017 | 2113366 | 25 | 261.52 | 4568 | 0.001 | 2109366 | 12 | 61.19 | 19522 | 0.006 |
| *H. t. florencia* | Hmel204017 | 1967332 | 29 | 163.58 | 6677 | 0.002 | 2065642 | 3 | 117.55 | 9292 | 0.003 |
| *H. t. linaresi* | Hmel204017 | 1851421 | 22 | 765.3 | 1502 | 0 | 1859622 | 13 | 99.62 | 11540 | 0.004 |
| *H. t. ssp. nov. ECU* | Hmel204017 | 1967446 | 14 | 227.52 | 5021 | 0.002 | 2089008 | 12 | 217.47 | 5252 | 0.002 |
| *H. t. thelxinoe* | Hmel204017 | 1897232 | 12 | 1669.44 | 586 | 0 | 2199918 | 0 | 179.64 | 5442 | 0.002 |
| *H. t. timareta f. contigua* | Hmel204017 | 1967333 | 21 | 219.12 | 4601 | 0.002 | 1894174 | 20 | 168.63 | 5979 | 0.002 |
| *H. t. timareta f. timareta* | Hmel204017 | 2290611 | 19 | 396.19 | 2716 | 0.001 | 1967917 | 17 | 155.17 | 6934 | 0.002 |
| *H. t. ssp. nov. COL* | Hmel204017 | 1967382 | 21 | 147.35 | 6899 | 0.003 | 2065945 | 15 | 82.77 | 12282 | 0.004 |
| *H. besckei* | Hmel206006 | 623821 | 32 | 97.56 | 2909 | 0.003 | 446294 | 8 | 74.45 | 3812 | 0.003 |
| *H. c. chioneus* | Hmel206006 | 552281 | 26 | 237.71 | 3714 | 0.001 | 635236 | 5 | 144.84 | 6096 | 0.002 |
| *H. c. cydnides* | Hmel206006 | 621246 | 35 | 230.14 | 3875 | 0.001 | 636247 | 7 | 111.31 | 8011 | 0.002 |
| *H. c. weymeri gustavi* | Hmel206006 | 551591 | 22 | 302.96 | 2709 | 0.001 | 635797 | 2 | 166.98 | 4916 | 0.002 |
| *H. c. weymeri weymeri* | Hmel206006 | 358310 | 31 | 369.33 | 2141 | 0.001 | 581335 | 0 | 319.73 | 2473 | 0.001 |

| Population | Scaffold | Position | CLR | $\alpha$ | $2N_es$ | $s$ | Position ($\alpha_{min}$) | CLR ($\alpha_{min}$) | $\alpha_{min}$ | $2N_es$ ($\alpha_{min}$) | $s$ ($\alpha_{min}$) |
|---|---|---|---|---|---|---|---|---|---|---|---|
| *H. c. zelinde* | Hmel206006 | 625531 | 16 | 354.12 | 2294 | 0.001 | 625781 | 14 | 312.76 | 2598 | 0.001 |
| *H. elevatus Ecuador* | Hmel206006 | 625339 | 44 | 206.24 | 6388 | 0.001 | 625089 | 34 | 193.59 | 6805 | 0.001 |
| *H. heurippa* | Hmel206006 | 692324 | 168 | 74.34 | 7707 | 0.004 | 622715 | 85 | 66.49 | 8616 | 0.004 |
| *H. m. amaryllis* | Hmel206006 | 792036 | 10 | 6846.76 | 189 | 0 | 624816 | 1 | 663.78 | 1952 | 0 |
| *H. m. cythera* | Hmel206006 | 327697 | 24 | 363.9 | 2483 | 0.001 | 327697 | 24 | 363.9 | 2483 | 0.001 |
| *H. m. ECU* | Hmel206006 | 754627 | 14 | 2689.83 | 387 | 0 | 604518 | 4 | 369.31 | 2819 | 0.001 |
| *H. m. malleti COL* | Hmel206006 | 519734 | 13 | 860.75 | 1194 | 0 | 330368 | 3 | 377.11 | 2726 | 0.001 |
| *H. m. malleti ECU* | Hmel206006 | 520036 | 17 | 670.59 | 2001 | 0 | 604349 | 1 | 256.42 | 5232 | 0.001 |
| *H. m. melpomene COL* | Hmel206006 | 435316 | 57 | 323.64 | 3074 | 0.001 | 635627 | 2 | 182.56 | 5450 | 0.001 |
| *H. m. melpomene FG* | Hmel206006 | 748824 | 25 | 446.97 | 1452 | 0.001 | 603265 | 10 | 113.82 | 5702 | 0.002 |
| *H. m. melpomene PAN* | Hmel206006 | 550748 | 34 | 108.81 | 9072 | 0.002 | 550448 | 3 | 106.69 | 9253 | 0.003 |
| *H. m. meriana* | Hmel206006 | 398903 | 39 | 187.53 | 2422 | 0.001 | 399353 | 28 | 176.16 | 2578 | 0.001 |
| *H. m. nanna NORTH* | Hmel206006 | 624824 | 26 | 114.64 | 7274 | 0.002 | 624874 | 26 | 114.5 | 7282 | 0.002 |
| *H. m. nanna SOUTH* | Hmel206006 | 697150 | 19 | 74.81 | 11146 | 0.004 | 593689 | 12 | 71 | 11745 | 0.004 |
| *H. m. plesseni* | Hmel206006 | 542803 | 20 | 607.55 | 1666 | 0 | 604459 | 1 | 287.69 | 3519 | 0.001 |
| *H. m. rosina* | Hmel206006 | 435387 | 21 | 579.17 | 1374 | 0 | 485890 | 4 | 372.9 | 2134 | 0.001 |
| *H. m. vicina* | Hmel206006 | 596517 | 13 | 555.86 | 1790 | 0 | 635772 | 2 | 150.84 | 6596 | 0.002 |
| *H. m. vulcanus* | Hmel206006 | 620745 | 37 | 204.59 | 3271 | 0.001 | 619745 | 33 | 161.41 | 4146 | 0.002 |
| *H. m. xenoclea* | Hmel206006 | 602412 | 12 | 256.85 | 3909 | 0.001 | 602712 | 8 | 224.35 | 4475 | 0.001 |
| *H. pachinus* | Hmel206006 | 602775 | 30 | 114.42 | 7436 | 0.002 | 603575 | 26 | 103.26 | 8240 | 0.003 |
| *H. t. florencia* | Hmel206006 | 619768 | 54 | 158.05 | 5210 | 0.002 | 604217 | 12 | 115.02 | 7160 | 0.002 |
| *H. t. linaresi* | Hmel206006 | 647619 | 30 | 294.32 | 2672 | 0.001 | 647569 | 28 | 294.19 | 2673 | 0.001 |
| *H. t. ssp. nov. ECU* | Hmel206006 | 622609 | 54 | 148.24 | 5681 | 0.002 | 635360 | 12 | 134.25 | 6273 | 0.002 |
| *H. t. thelxinoe* | Hmel206006 | 619714 | 49 | 200.32 | 3824 | 0.001 | 619764 | 48 | 199.77 | 3834 | 0.001 |
| *H. t. timareta f. contigua* | Hmel206006 | 700365 | 20 | 884.42 | 865 | 0 | 736970 | 0 | 249.31 | 3068 | 0.001 |
| *H. t. timareta f. timareta* | Hmel206006 | 739820 | 17 | 439.77 | 1821 | 0.001 | 503640 | 3 | 409.39 | 1956 | 0.001 |
| *H. t. ssp. nov. COL* | Hmel206006 | 600699 | 80 | 63.47 | 10739 | 0.004 | 599899 | 77 | 62.48 | 10908 | 0.004 |
| *H. besckei* | Hmel208051 | 754245 | 18 | 63.65 | 5209 | 0.006 | 755845 | 12 | 46.58 | 7119 | 0.008 |
| *H. c. chioneus* | Hmel208051 | 1042680 | 19 | 158.11 | 7208 | 0.003 | 911321 | 11 | 91.25 | 12489 | 0.004 |
| *H. c. cydnides* | Hmel208051 | 1043501 | 22 | 148.57 | 7166 | 0.003 | 1043501 | 22 | 148.57 | 7166 | 0.003 |
| *H. c. weymeri gustavi* | Hmel208051 | 1039926 | 45 | 81.67 | 12625 | 0.005 | 934568 | 21 | 72.99 | 14126 | 0.005 |
| *H. c. weymeri weymeri* | Hmel208051 | 1043638 | 21 | 341.79 | 2883 | 0.001 | 638197 | 9 | 132.55 | 7435 | 0.003 |
| *H. c. zelinde* | Hmel208051 | 687356 | 18 | 424.56 | 2643 | 0.001 | 948375 | 7 | 136.57 | 8216 | 0.003 |
| *H. elevatus Ecuador* | Hmel208051 | 1044514 | 49 | 97.68 | 16657 | 0.004 | 929249 | 9 | 90.21 | 18036 | 0.005 |
| *H. heurippa* | Hmel208051 | 982755 | 352 | 13.28 | 56565 | 0.029 | 983505 | 284 | 12.93 | 58060 | 0.03 |
| *H. m. amaryllis* | Hmel208051 | 1048965 | 13 | 801.01 | 2058 | 0.001 | 1045165 | 4 | 223.02 | 7391 | 0.002 |
| *H. m. cythera* | Hmel208051 | 846537 | 21 | 442.22 | 2903 | 0.001 | 955998 | 17 | 136.1 | 9432 | 0.003 |
| *H. m. ECU* | Hmel208051 | 1024299 | 26 | 94.47 | 14695 | 0.004 | 1024249 | 26 | 94.32 | 14719 | 0.004 |
| *H. m. malleti COL* | Hmel208051 | 1049946 | 11 | 209.12 | 6954 | 0.002 | 1049946 | 11 | 209.12 | 6954 | 0.002 |
| *H. m. malleti ECU* | Hmel208051 | 1050196 | 11 | 1123.99 | 1587 | 0 | 637708 | 0 | 359.68 | 4960 | 0.001 |
| *H. m. melpomene COL* | Hmel208051 | 694915 | 15 | 955.82 | 1439 | 0 | 745370 | 1 | 186.47 | 7377 | 0.002 |
| *H. m. melpomene FG* | Hmel208051 | 706418 | 10 | 602.4 | 1549 | 0.001 | 1029013 | 7 | 177.13 | 5267 | 0.002 |
| *H. m. melpomene PAN* | Hmel208051 | 992145 | 20 | 446.35 | 3005 | 0.001 | 905339 | 6 | 175.09 | 7660 | 0.002 |

| **Population** | **Scaffold** | **Position** | **CLR** | ***α*** | **2*N*e*s*** | ***s*** | **Position ($α_{min}$)** | **CLR ($α_{min}$)** | **$α_{min}$** | **2*N*e*s* ($α_{min}$)** | ***s* ($α_{min}$)** |
|---|---|---|---|---|---|---|---|---|---|---|---|
| *H. m. meriana* | Hmel208051 | 1024688 | 41 | 85.45 | 7938 | 0.005 | 1053241 | 33 | 79.51 | 8530 | 0.005 |
| *H. m. nanna NORTH* | Hmel208051 | 1037151 | 165 | 21.66 | 49737 | 0.018 | 1040451 | 129 | 20.15 | 53472 | 0.02 |
| *H. m. nanna SOUTH* | Hmel208051 | 945416 | 17 | 35.09 | 30698 | 0.011 | 761453 | 5 | 23.47 | 45887 | 0.017 |
| *H. m. plesseni* | Hmel208051 | 1100404 | 11 | 658.06 | 2288 | 0.001 | 1023896 | 3 | 135.07 | 11149 | 0.003 |
| *H. m. rosina* | Hmel208051 | 903527 | 16 | 253.68 | 3794 | 0.002 | 956032 | 10 | 159.2 | 6045 | 0.002 |
| *H. m. vicina* | Hmel208051 | 1069609 | 27 | 95.31 | 14433 | 0.004 | 1053206 | 20 | 78.74 | 17470 | 0.005 |
| *H. m. vulcanus* | Hmel208051 | 981677 | 25 | 185 | 4633 | 0.002 | 986477 | 19 | 92.31 | 9286 | 0.004 |
| *H. m. xenoclea* | Hmel208051 | 1048964 | 13 | 662.39 | 2014 | 0.001 | 1052915 | 4 | 178.7 | 7466 | 0.002 |
| *H. pachinus* | Hmel208051 | 958623 | 33 | 124.22 | 8836 | 0.003 | 957523 | 33 | 78.89 | 13913 | 0.005 |
| *H. t. florencia* | Hmel208051 | 1046270 | 132 | 32.25 | 31761 | 0.012 | 1031368 | 98 | 30.46 | 33621 | 0.013 |
| *H. t. linaresi* | Hmel208051 | 1053301 | 48 | 72.19 | 14440 | 0.006 | 1053251 | 47 | 72.19 | 14440 | 0.006 |
| *H. t. ssp. nov. ECU* | Hmel208051 | 1045877 | 92 | 35.49 | 29744 | 0.011 | 1046027 | 21 | 35.22 | 29971 | 0.011 |
| *H. t. thelxinoe* | Hmel208051 | 909917 | 58 | 68.56 | 13188 | 0.006 | 910767 | 54 | 53.6 | 16868 | 0.007 |
| *H. t. timareta f. contigua* | Hmel208051 | 1045354 | 53 | 51.84 | 17529 | 0.008 | 1045254 | 53 | 51.71 | 17571 | 0.008 |
| *H. t. timareta f. timareta* | Hmel208051 | 1040433 | 50 | 47.32 | 20836 | 0.008 | 1043433 | 39 | 44.08 | 22366 | 0.009 |
| *H. t. ssp. nov. COL* | Hmel208051 | 1024784 | 108 | 27.27 | 34225 | 0.015 | 1024684 | 108 | 27.26 | 34240 | 0.015 |
| *H. besckei* | Hmel219003 | 5636365 | 10 | 382.71 | 653 | 0.001 | 5656167 | 9 | 234.87 | 1063 | 0.001 |
| *H. c. chioneus* | Hmel219003 | 5569780 | 55 | 103.99 | 6858 | 0.002 | 5569780 | 55 | 103.99 | 6858 | 0.002 |
| *H. c. cydnides* | Hmel219003 | 5396768 | 101 | 93.28 | 7151 | 0.003 | 5396768 | 101 | 93.28 | 7151 | 0.003 |
| *H. c. weymeri gustavi* | Hmel219003 | 5538911 | 79 | 126.32 | 4931 | 0.002 | 5286326 | 25 | 109.02 | 5713 | 0.002 |
| *H. c. weymeri weymeri* | Hmel219003 | 5553535 | 205 | 45.63 | 13006 | 0.005 | 5553235 | 164 | 45.5 | 13044 | 0.005 |
| *H. c. zelinde* | Hmel219003 | 5570029 | 85 | 75.96 | 9187 | 0.003 | 5570329 | 42 | 75.18 | 9283 | 0.003 |
| *H. elevatus Ecuador* | Hmel219003 | 5568930 | 68 | 314.01 | 3574 | 0.001 | 5576931 | 16 | 192.94 | 5816 | 0.001 |
| *H. heurippa* | Hmel219003 | 5611280 | 98 | 154.57 | 2329 | 0.001 | 5428619 | 91 | 100.02 | 3599 | 0.002 |
| *H. m. amaryllis* | Hmel219003 | 5552829 | 22 | 640.11 | 1581 | 0 | 5555129 | 19 | 441.36 | 2292 | 0.001 |
| *H. m. cythera* | Hmel219003 | 5246608 | 183 | 46.31 | 16549 | 0.005 | 5245608 | 144 | 45.94 | 16683 | 0.005 |
| *H. m. ECU* | Hmel219003 | 5576937 | 56 | 96.52 | 9250 | 0.003 | 5577787 | 52 | 95.49 | 9350 | 0.003 |
| *H. m. malleti COL* | Hmel219003 | 5252414 | 26 | 323.26 | 2600 | 0.001 | 5254414 | 19 | 196.08 | 4287 | 0.001 |
| *H. m. malleti ECU* | Hmel219003 | 5567800 | 39 | 469.27 | 2247 | 0.001 | 5576701 | 19 | 160.88 | 6554 | 0.002 |
| *H. m. melpomene COL* | Hmel219003 | 5588298 | 23 | 1036.13 | 826 | 0 | 5254406 | 6 | 426.4 | 2008 | 0.001 |
| *H. m. melpomene FG* | Hmel219003 | 5553326 | 40 | 184.69 | 2878 | 0.001 | 5554226 | 40 | 181.55 | 2928 | 0.001 |
| *H. m. melpomene PAN* | Hmel219003 | 5254467 | 58 | 122.63 | 6589 | 0.002 | 5254417 | 58 | 122.44 | 6599 | 0.002 |
| *H. m. meriana* | Hmel219003 | 5452642 | 46 | 241.13 | 1623 | 0.001 | 5244725 | 29 | 148.28 | 2639 | 0.002 |
| *H. m. nanna NORTH* | Hmel219003 | 5576562 | 96 | 37.54 | 17732 | 0.006 | 5576712 | 95 | 37.52 | 17742 | 0.006 |
| *H. m. nanna SOUTH* | Hmel219003 | 5597149 | 76 | 53.93 | 12342 | 0.004 | 5607500 | 18 | 45.49 | 14634 | 0.005 |
| *H. m. plesseni* | Hmel219003 | 5567856 | 29 | 333.57 | 2549 | 0.001 | 5254487 | 26 | 145.16 | 5858 | 0.002 |
| *H. m. rosina* | Hmel219003 | 5551670 | 47 | 201.16 | 3144 | 0.001 | 5255329 | 30 | 145.08 | 4360 | 0.002 |
| *H. m. vicina* | Hmel219003 | 5586901 | 36 | 297.39 | 2879 | 0.001 | 5551797 | 24 | 155.19 | 5517 | 0.002 |
| *H. m. vulcanus* | Hmel219003 | 5553765 | 50 | 237.42 | 2287 | 0.001 | 5577818 | 26 | 212.72 | 2553 | 0.001 |
| *H. m. xenoclea* | Hmel219003 | 5569609 | 25 | 323.88 | 2512 | 0.001 | 5576810 | 6 | 145.39 | 5597 | 0.002 |
| *H. pachinus* | Hmel219003 | 5551837 | 78 | 116.07 | 5952 | 0.002 | 5555137 | 69 | 111.15 | 6215 | 0.002 |
| *H. t. florencia* | Hmel219003 | 5715417 | 124 | 98.33 | 5559 | 0.002 | 5251087 | 41 | 48.88 | 11184 | 0.005 |
| *H. t. linaresi* | Hmel219003 | 5610520 | 98 | 190.54 | 2748 | 0.001 | 5388490 | 69 | 89.17 | 5872 | 0.003 |

| **Population** | **Scaffold** | **Position** | **CLR** | ***α*** | **2$N_e s$** | ***s*** | **Position ($α_{min}$)** | **CLR ($α_{min}$)** | **$α_{min}$** | **2$N_e s$ ($α_{min}$)** | ***s* ($α_{min}$)** |
|---|---|---|---|---|---|---|---|---|---|---|---|
| *H. t. ssp. nov. ECU* | Hmel219003 | 5715383 | 137 | 89.06 | 6202 | 0.003 | 5251238 | 26 | 77.57 | 7120 | 0.003 |
| *H. t. thelxinoe* | Hmel219003 | 5611298 | 171 | 88.73 | 4321 | 0.003 | 5611098 | 168 | 88.41 | 4336 | 0.003 |
| *H. t. timareta f. contigua* | Hmel219003 | 5250567 | 175 | 32.56 | 14290 | 0.007 | 5250067 | 132 | 32.27 | 14420 | 0.007 |
| *H. t. timareta f. timareta* | Hmel219003 | 5551811 | 114 | 88.55 | 5363 | 0.003 | 5251171 | 38 | 75.12 | 6322 | 0.003 |
| *H. t. ssp. nov. COL* | Hmel219003 | 5583888 | 412 | 21.17 | 21110 | 0.011 | 5575787 | 274 | 19.51 | 22910 | 0.012 |

**Supplementary Table 6:** List of additional genes with significant colour pattern associations on the cortex scaffold from Nadeau *et al*.[7] that overlap with or are in proximity of selection signatures detected in this study.

| **Hm/Hmel2.5 gene IDs** | **He/H. erato demophoon v1 gene IDs** | **Putative gene name** | **Position relative to *cortex*** | **Comment** | **Number ID in Suppl. Figure 6/11** |
|---|---|---|---|---|---|
| HM00002/tba | HERA000036/tba | Acylpeptide hydrolase | downstream | | 6 |
| HM00008/tba | HERA000040/tba | | downstream | | 7 |
| HM00012/tba | HERA000042/tba | *CG2519* | downstream | | 8 |
| HM00019/tba | HERA000052/tba | *BmSuc2* | downstream | | 9 |
| HM00026/tba | HERA000077 | Poly-A specific ribonuclease *parn* | upstream | | 14 |
| HM00031/tba | HERA000083/tba | | upstream | | 15 |
| HM00032/tba | HERA000084/tba | Zinc phosphodiesterase | upstream | | 16 |
| HM00033/tba | HERA000085/tba | Serine/threonine-protein kinase *LMTK1* | upstream | strong, consistent selection signal in *H. melpomene* | 17 |
| HM00034/tba | HERA000086/tba | WD repeat domain *Wdr13* | upstream | | 18 |
| HM00035/tba | HERA000087/tba | *domeless* | upstream | truncated sequence | 19/10 |
| HM00036/tba | HERA000061/tba | *washout / WAS homologue 1* | upstream | strong evidence in Nadeau *et al.* 2016 | 20/11 |
| HM00037/tba | annotated as evm.TU.Herato1505.96 in *H. erato demophoon* v1 | *domeless* | upstream | complete sequence, not considered in Nadeau *et al.* 2016 | 21/12 |
| HM00038/tba | HERA000062/tba | *lethal (2) k05819 CG3054* | upstream | strong, consistent selection signal in *H. erato* | 22/13 |
| HM00052/tba | HERA000076/tba | | upstream | | 23 |

**Supplementary Table 7:** Sample information and genotyping statistics for all samples from the *Heliconius erato*-clade from Van Belleghem *et al.*[2].

| SequenceID | EarthCapeID | Taxon name | Country | Latitude | Longitude | Accession |
|---|---|---|---|---|---|---|
| BC2115 | BC2115 | *Heliconius erato amalfreda* | Suriname | -4.946897 | 55.183386 | SAMN05224103 |
| BC2124 | BC2124 | *Heliconius erato amalfreda* | Suriname | -5.943486 | 55.186072 | SAMN05224104 |
| STRI_WOM_5779 | STRI_WOM_5779 | *Heliconius erato amalfreda* | Suriname | -5.940653 | 55.189922 | SAMN05224208 |
| STRI_WOM_5780 | STRI_WOM_5780 | *Heliconius erato amalfreda* | Suriname | -5.940653 | 55.189922 | SAMN05224209 |
| STRI_WOM_5781 | STRI_WOM_5781 | *Heliconius erato amalfreda* | Suriname | -4.932733 | 55.200803 | SAMN05224210 |
| STRI_WOM_0057 | STRI_WOM_0057 | *Heliconius erato chestertonii* | Colombia | 3.884017 | -76.589367 | SAMN05224192 |
| STRI_WOM_0058 | STRI_WOM_0058 | *Heliconius erato chestertonii* | Colombia | 3.884017 | -76.589367 | SAMN05224193 |
| STRI_WOM_0059 | STRI_WOM_0059 | *Heliconius erato chestertonii* | Colombia | 3.884017 | -76.589367 | SAMN05224194 |
| 3661 | CS003661 | *Heliconius erato chestertonii* | Colombia | 3.884017 | -76.589367 | SAMN05224096 |
| 3662 | CS003662 | *Heliconius erato chestertonii* | Colombia | 3.884017 | -76.589367 | SAMN05224097 |
| 3663 | CS003663 | *Heliconius erato chestertonii* | Colombia | 3.884017 | -76.589367 | SAMN05224098 |
| 3664 | CS003664 | *Heliconius erato chestertonii* | Colombia | 3.884017 | -76.589367 | SAMN05224099 |
| cyrbia_004 | CYR004 | *Heliconius erato cyrbia* | Ecuador | -3.726389 | -79.836667 | SAMN05224122 |
| cyrbia_005 | CYR005 | *Heliconius erato cyrbia* | Ecuador | -3.726389 | -79.836667 | SAMN05224123 |
| cyrbia_023 | CYR023 | *Heliconius erato cyrbia* | Ecuador | -3.726389 | -79.836667 | SAMN05224124 |
| cyrbia_024 | CYR024 | *Heliconius erato cyrbia* | Ecuador | -3.726389 | -79.836667 | SAMN05224125 |
| Pet_ED3 | Pet_ED3 | *Heliconius erato demophoon* | Panama | -9.129444 | 79.715278 | SAMN05224182 |
| Pet_ED4 | Pet_ED4 | *Heliconius erato demophoon* | Panama | -9.129444 | 79.715278 | SAMN05224183 |
| Pet_ED5 | Pet_ED5 | *Heliconius erato demophoon* | Panama | -9.129444 | 79.715278 | SAMN05224184 |
| Pet_ED6 | Pet_ED6 | *Heliconius erato demophoon* | Panama | -9.129444 | 79.715278 | SAMN05224185 |
| STRI_WOM_0033 | STRI_WOM_0033 | *Heliconius erato demophoon* | Panama | -9.1525 | 78.689722 | SAMN05224188 |
| STRI_WOM_0082 | STRI_WOM_0082 | *Heliconius erato demophoon* | Panama | -9.1525 | 78.689722 | SAMN05224195 |
| STRI_WOM_0087 | STRI_WOM_0087 | *Heliconius erato demophoon* | Panama | -9.1525 | 78.689722 | SAMN05224196 |
| STRIWOM1284 | STRI_WOM_1284 | *Heliconius erato demophoon* | Panama | -9.1525 | 78.689722 | SAMN05224198 |
| STRIWOM5353 | STRI_WOM_5353 | *Heliconius erato demophoon* | Panama | -9.1525 | 78.689722 | SAMN05224202 |
| STRIWOM5362 | STRI_WOM_5362 | *Heliconius erato demophoon* | Panama | -9.1525 | 78.689722 | SAMN05224203 |
| BC2563* | BC_2563 | *Heliconius erato emma* | Peru | -5.29499 | -78.381 | SAMN08049955 |
| BC2577* | BC_2577 | *Heliconius erato emma* | Peru | -5.29499 | -78.381 | SAMN08049956 |
| BC2578* | BC_2578 | *Heliconius erato emma* | Peru | -5.29499 | -78.381 | SAMN08049957 |
| BC2579* | BC_2579 | *Heliconius erato emma* | Peru | -5.29499 | -78.381 | SAMN08049958 |
| GS020redo | GS020 | *Heliconius erato emma* | Peru | -6.181944 | -76.247222 | SAMN05224127 |
| GS021redo | GS021 | *Heliconius erato emma* | Peru | -6.181944 | -76.247222 | SAMN05224128 |
| NCS_1671 | NCS1671 | *Heliconius erato emma* | Peru | -6.181944 | -76.247222 | SAMN05224154 |
| NCS_1672 | NCS1672 | *Heliconius erato emma* | Peru | -6.181944 | -76.247222 | SAMN05224155 |
| NCS_1673 | NCS1673 | *Heliconius erato emma* | Peru | -6.181944 | -76.247222 | SAMN05224156 |
| NCS_1674 | NCS1674 | *Heliconius erato emma* | Peru | -6.181944 | -76.247222 | SAMN05224157 |
| NCS_1675 | NCS1675 | *Heliconius erato emma* | Peru | -6.181944 | -76.247222 | SAMN05224158 |
| NCS_2005 | NCS2005 | *Heliconius erato erato* | French Guiana | -4.638611 | 52.301667 | SAMN05224160 |
| NCS_2012 | NCS2012 | *Heliconius erato erato* | French Guiana | -4.638611 | 52.301667 | SAMN05224161 |
| NCS_2020 | NCS2020 | *Heliconius erato erato* | French Guiana | -4.585 | 52.245556 | SAMN05224162 |
| NCS_2023 | NCS2023 | *Heliconius erato erato* | French Guiana | -4.638611 | 52.301667 | SAMN05224163 |
| NCS_2025 | NCS2025 | *Heliconius erato erato* | French Guiana | -4.585 | 52.245556 | SAMN05224164 |

| SequenceID | EarthCapeID | Taxon name | Country | Latitude | Longitude | Accession |
|---|---|---|---|---|---|---|
| NCS_2556 | NCS2556 | *Heliconius erato erato* | French Guiana | -4.621944 | 52.376111 | SAMN05224174 |
| BC_3277 | BC_3277 | *Heliconius erato etylus* | Ecuador | -1.97786 | -78.00945 | SAMN05224110 |
| BC_3278 | BC_3278 | *Heliconius erato etylus* | Ecuador | -1.97786 | -78.00945 | SAMN05224111 |
| BC_3280 | BC_3280 | *Heliconius erato etylus* | Ecuador | -1.97786 | -78.00945 | SAMN05224112 |
| BC_3281 | BC_3281 | *Heliconius erato etylus* | Ecuador | -1.97786 | -78.00945 | SAMN05224113 |
| BC_3282 | BC_3282 | *Heliconius erato etylus* | Ecuador | -1.97786 | -78.00945 | SAMN05224114 |
| BC2635* | BC_2635 | *Heliconius erato favorinus* | Peru | -6.4174 | -77.44329 | SAMN08049959 |
| BC2637* | BC_2637 | *Heliconius erato favorinus* | Peru | -6.4174 | -77.44329 | SAMN08049960 |
| BC2638* | BC_2638 | *Heliconius erato favorinus* | Peru | -6.4174 | -77.44329 | SAMN08049961 |
| BC2639* | BC_2639 | *Heliconius erato favorinus* | Peru | -6.4174 | -77.44329 | SAMN08049962 |
| GS012redo | GS012 | *Heliconius erato favorinus* | Peru | -6.461389 | -76.341944 | SAMN05224126 |
| NCS_0471 | NCS0471 | *Heliconius erato favorinus* | Peru | -6.474167 | -76.010278 | SAMN05224148 |
| NCS_0473 | NCS0473 | *Heliconius erato favorinus* | Peru | -6.474167 | -76.010278 | SAMN05224149 |
| NCS_0476 | NCS0476 | *Heliconius erato favorinus* | Peru | -6.474167 | -76.010278 | SAMN05224150 |
| NCS_0478 | NCS0478 | *Heliconius erato favorinus* | Peru | -6.474167 | -76.010278 | SAMN05224151 |
| NCS_0479 | NCS0479 | *Heliconius erato favorinus* | Peru | -6.474167 | -76.010278 | SAMN05224152 |
| NCS_2554 | NCS2554 | *Heliconius erato favorinus* | Peru | -6.474167 | -76.010278 | SAMN05224172 |
| NCS_2555 | NCS2555 | *Heliconius erato favorinus* | Peru | -6.474167 | -76.010278 | SAMN05224173 |
| STRI_WOM_0042 | STRI_WOM_0042 | *Heliconius erato hydara* | Panama | -9.1525 | 78.689722 | SAMN05224191 |
| NCS_1179 | NCS1179 | *Heliconius erato hydara* | French Guiana | -4.703611 | 52.303611 | SAMN05224153 |
| NCS_1979 | NCS1979 | *Heliconius erato hydara* | French Guiana | -4.571667 | 52.223333 | SAMN05224159 |
| NCS_2080 | NCS2080 | *Heliconius erato hydara* | French Guiana | -4.607778 | 52.2725 | SAMN05224165 |
| NCS_2211 | NCS2211 | *Heliconius erato hydara* | French Guiana | -4.547222 | 52.170278 | SAMN05224166 |
| NCS_2217 | NCS2217 | *Heliconius erato hydara* | French Guiana | -4.544444 | 52.1525 | SAMN05224167 |
| STRI_WOM_0039 | STRI_WOM_0039 | *Heliconius erato hydara* | Panama | -9.1525 | 78.689722 | SAMN05224189 |
| STRI_WOM_0040 | STRI_WOM_0040 | *Heliconius erato hydara* | Panama | -9.1525 | 78.689722 | SAMN05224190 |
| STRI_WOM_0088 | STRI_WOM_0088 | *Heliconius erato hydara* | Panama | -9.1525 | 78.689722 | SAMN05224197 |
| STRI_WOM_5193 | STRI_WOM_5193 | *Heliconius erato hydara* | Panama | -9.1525 | 78.689722 | SAMN05224200 |
| STRI_WOM_5351 | STRI_WOM_5351 | *Heliconius erato hydara* | Panama | -9.1525 | 78.689722 | SAMN05224201 |
| BC_0411 | BC0411 | *Heliconius erato lativitta* | Ecuador | -1.098333 | 77.583889 | SAMN05224101 |
| lativitta_01 | LAT01 | *Heliconius erato lativitta* | Ecuador | -1.098333 | 77.583889 | SAMN05224137 |
| lativitta_02 | LAT02 | *Heliconius erato lativitta* | Ecuador | -1.098333 | 77.583889 | SAMN05224138 |
| lativitta_03 | LAT03 | *Heliconius erato lativitta* | Ecuador | -1.098333 | 77.583889 | SAMN05224139 |
| lativitta_04 | LAT04 | *Heliconius erato lativitta* | Ecuador | -0.7125 | 77.583889 | SAMN05224140 |
| BC_0410 | BC_0410 | *Heliconius erato notabilis* | Ecuador | -1.81337 | -78.04507 | SAMN05224100 |
| BC_3223 | BC_3223 | *Heliconius erato notabilis* | Ecuador | -1.81337 | -78.04507 | SAMN05224105 |
| BC_3224 | BC_3224 | *Heliconius erato notabilis* | Ecuador | -1.81337 | -78.04507 | SAMN05224106 |
| BC_3225 | BC_3225 | *Heliconius erato notabilis* | Ecuador | -1.81337 | -78.04507 | SAMN05224107 |
| BC_3227 | BC_3227 | *Heliconius erato notabilis* | Ecuador | -1.82259 | -78.04406 | SAMN05224108 |
| BC_3228 | BC_3228 | *Heliconius erato notabilis* | Ecuador | -1.82259 | -78.04406 | SAMN05224109 |
| notabilis_01 | NOT01 | *Heliconius erato notabilis* | Ecuador | -1.399167 | 78.181111 | SAMN05224178 |
| notabilis_02 | NOT02 | *Heliconius erato notabilis* | Ecuador | -1.399167 | 78.181111 | SAMN05224179 |
| notabilis_03 | NOT03 | *Heliconius erato notabilis* | Ecuador | -1.399167 | 78.181111 | SAMN05224180 |
| notabilis_04 | NOT04 | *Heliconius erato notabilis* | Ecuador | -1.399167 | 78.181111 | SAMN05224181 |

| SequenceID | EarthCapeID | Taxon name | Country | Latitude | Longitude | Accession |
|---|---|---|---|---|---|---|
| M_3654 | CS003654 | *Heliconius erato venus* | Colombia | 3.5311 | -76.753383 | SAMN05224141 |
| M_3655 | CS003655 | *Heliconius erato venus* | Colombia | 3.5311 | -76.753383 | SAMN05224142 |
| M_3656 | CS003656 | *Heliconius erato venus* | Colombia | 3.5311 | -76.753383 | SAMN05224143 |
| M_3657 | CS003657 | *Heliconius erato venus* | Colombia | 3.5311 | -76.753383 | SAMN05224144 |
| M_3659 | CS003659 | *Heliconius erato venus* | Colombia | 3.5311 | -76.753383 | SAMN05224145 |
| BC2565* | BC_2565 | *Heliconius himera* | Peru | -5.43724 | -78.4714 | SAMN08049963 |
| BC2566* | BC_2566 | *Heliconius himera* | Peru | -5.43724 | -78.4714 | SAMN08049964 |
| BC2567* | BC_2567 | *Heliconius himera* | Peru | -5.43724 | -78.4714 | SAMN08049965 |
| BC2570* | BC_2570 | *Heliconius himera* | Peru | -5.43724 | -78.4714 | SAMN08049966 |
| himera_001 | HIM001 | *Heliconius himera* | Ecuador | -4.276111 | -79.195833 | SAMN05224132 |
| himera_002 | HIM002 | *Heliconius himera* | Ecuador | -4.276111 | -79.195833 | SAMN05224133 |
| himera_003 | HIM003 | *Heliconius himera* | Ecuador | -4.276111 | -79.195833 | SAMN05224134 |
| himera_006 | HIM006 | *Heliconius himera* | Ecuador | -4.276111 | -79.195833 | SAMN05224135 |
| himera_030 | HIM030 | *Heliconius himera* | Ecuador | -4.276111 | -79.195833 | SAMN05224136 |
| LM CI94-13 | - | *Heliconius hermathena* | Brazil | -2.450000 | -54.700000 | SAMN05224129 |
| LM CI94-14 | - | *Heliconius hermathena* | Brazil | -2.450000 | -54.700000 | SAMN05224130 |
| LM CI94-15 | - | *Heliconius hermathena* | Brazil | -2.450000 | -54.700000 | SAMN05224131 |

**Supplementary Table 8:** Position, composite likelihood-ratio statistics (CLR) and strength of selection ($\alpha$, $2N_es$, and $s$) for the highest CLR and the smallest $\alpha$ value on each colour pattern scaffold ($\alpha_{min}$) for *H. erato*. Additional relevant peaks on scaffolds are also given. Data are from SweepFinder2[3,4] runs with background site frequency spectrum estimated from background scaffolds.

| Population | Locus | Scaffold | Position | CLR | $\alpha$ | $2N_es$ | $s$ | Position ($\alpha_{min}$) | CLR ($\alpha_{min}$) | $\alpha_{min}$ | $2N_es$ ($\alpha_{min}$) | $s$ ($\alpha_{min}$) |
|---|---|---|---|---|---|---|---|---|---|---|---|---|
| *H. e. amalfreda* | *WntA* | Herato1001 | 4642379 | 561 | 24.28 | 39792 | 0.006 | 4646429 | 514 | 23.61 | 40923 | 0.006 |
| *H. e. cyrbiaN* | *WntA* | Herato1001 | 4402129 | 123 | 50.94 | 8607 | 0.003 | 4402279 | 121 | 50.9 | 8613 | 0.003 |
| *H. e. demophoon* | *WntA* | Herato1001 | 4410429 | 146 | 119.4 | 8119 | 0.001 | 4410429 | 146 | 119.4 | 8119 | 0.001 |
| *H. e. emma* | *WntA* | Herato1001 | 4651630 | 289 | 52.24 | 19995 | 0.003 | 4649530 | 287 | 49.4 | 21146 | 0.003 |
| *H. e. erato* | *WntA* | Herato1001 | 4642325 | 478 | 27.69 | 33496 | 0.005 | 4646375 | 456 | 26.84 | 34558 | 0.006 |
| *H. e. etylus* | *WntA* | Herato1001 | 4646229 | 701 | 16.24 | 58092 | 0.009 | 4698580 | 696 | 10.22 | 92334 | 0.015 |
| *H. e. favorinus* | *WntA* | Herato1001 | 4654380 | 250 | 28.24 | 36989 | 0.005 | 4668580 | 240 | 23.16 | 45096 | 0.007 |
| *H. e. hydaraFG* | *WntA* | Herato1001 | 4668734 | 306 | 25.55 | 36301 | 0.006 | 4667884 | 298 | 25.4 | 36515 | 0.006 |
| *H. e. hydaraP* | *WntA* | Herato1001 | 5302745 | 123 | 177.44 | 5464 | 0.001 | 4719235 | 46 | 87.15 | 11124 | 0.002 |
| *H.e. lativitta* | *WntA* | Herato1001 | 4651584 | 363 | 41.07 | 23599 | 0.004 | 4649684 | 345 | 40.29 | 24056 | 0.004 |
| *H. e. notabilis* | *WntA* | Herato1001 | 4648024 | 909 | 14.09 | 66925 | 0.011 | 4677475 | 763 | 9.88 | 95453 | 0.015 |
| *H. e. venus* | *WntA* | Herato1001 | 4405179 | 208 | 40.16 | 15539 | 0.004 | 4403179 | 181 | 38.54 | 16193 | 0.004 |
| *H. e. amalfreda* | *cortex* | Herato1505 | 2494767 | 1628 | 13 | 111467 | 0.017 | 2499467 | 1543 | 12.91 | 112233 | 0.017 |
| *H. e. cyrbiaN* | *cortex* | Herato1505 | 2131207 | 436 | 6.16 | 106722 | 0.035 | 2130806 | 434 | 6.16 | 106744 | 0.035 |
| *H. e. demophoon* | *cortex* | Herato1505 | 2277009 | 1050 | 13.99 | 103964 | 0.016 | 2267559 | 1039 | 13.93 | 104361 | 0.016 |
| *H. e. emma* | *cortex* | Herato1505 | 2496694 | 1392 | 16.03 | 97743 | 0.014 | 2497894 | 1382 | 16 | 97946 | 0.014 |
| *H. e. erato* | *cortex* | Herato1505 | 2493655 | 1483 | 14.43 | 96430 | 0.016 | 2490155 | 1455 | 14.38 | 96762 | 0.016 |
| *H. e. etylus* | *cortex* | Herato1505 | 2494192 | 1312 | 15.07 | 93917 | 0.015 | 2497592 | 1271 | 15.01 | 94276 | 0.015 |
| *H. e. favorinus* | *cortex* | Herato1505 | 2495987 | 2558 | 8.04 | 194973 | 0.028 | 2494987 | 2552 | 8.04 | 195006 | 0.028 |
| *H. e. hydaraFG* | *cortex* | Herato1505 | 2490958 | 1552 | 12.45 | 111745 | 0.018 | 2488308 | 1531 | 12.39 | 112300 | 0.018 |
| *H. e. hydaraP* | *cortex* | Herato1505 | 2985526 | 289 | 29.8 | 48802 | 0.008 | 2999826 | 56 | 21.55 | 67476 | 0.01 |
| *H.e. lativitta* | *cortex* | Herato1505 | 2491864 | 1252 | 15.96 | 91073 | 0.014 | 2489914 | 1239 | 15.9 | 91434 | 0.014 |
| *H. e. notabilis* | *cortex* | Herato1505 | 2497650 | 1387 | 15.2 | 93112 | 0.015 | 2501850 | 1354 | 14.76 | 95869 | 0.015 |
| | | | 1963287 | 472 | 49.76 | 28438 | 0.005 | 2065039 | 155 | 37.53 | 37703 | 0.006 |
| *H. e. venus* | *cortex* | Herato1505 | 2069952 | 874 | 5.74 | 162992 | 0.038 | 2130504 | 791 | 4.82 | 194354 | 0.046 |
| *H. e. amalfreda* | *optix* | Herato1801 | 1304730 | 997 | 6.66 | 172295 | 0.027 | 1304030 | 995 | 6.66 | 172319 | 0.027 |
| | | | 1375134 | 936 | 11.31 | 101441 | 0.016 | 1356333 | 509 | 10.52 | 109048 | 0.017 |
| *H. e. cyrbiaN* | *optix* | Herato1801 | 921205 | 174 | 37.21 | 13990 | 0.005 | 1302732 | 68 | 23.15 | 22491 | 0.007 |
| *H. e. demophoon* | *optix* | Herato1801 | 938505 | 66 | 340.1 | 3385 | 0.001 | 1296221 | 3 | 74.7 | 15411 | 0.002 |
| *H. e. emma* | *optix* | Herato1801 | 1305984 | 449 | 13.21 | 93912 | 0.014 | 1303583 | 418 | 13.11 | 94614 | 0.014 |
| *H. e. erato* | *optix* | Herato1801 | 1380478 | 1005 | 11.02 | 99959 | 0.016 | 1305775 | 913 | 7.61 | 144693 | 0.023 |
| | | | 1303875 | 922 | 7.62 | 144522 | 0.023 | 1305775 | 913 | 7.61 | 144693 | 0.023 |
| *H. e. etylus* | *optix* | Herato1801 | 1304785 | 423 | 10.67 | 104966 | 0.017 | 1300385 | 408 | 10.58 | 105893 | 0.017 |
| | | | 1407742 | 405 | 21.46 | 52202 | 0.008 | 1384040 | 390 | 20.89 | 53626 | 0.009 |
| *H. e. favorinus* | *optix* | Herato1801 | 1381433 | 285 | 32.38 | 38310 | 0.006 | 1303679 | 182 | 24.3 | 51053 | 0.007 |
| | | | 1304379 | 184 | 24.35 | 50937 | 0.007 | 1303679 | 182 | 24.3 | 51053 | 0.007 |
| *H. e. hydaraFG* | *optix* | Herato1801 | 1426489 | 4187 | 3.98 | 276447 | 0.045 | 1431689 | 4152 | 3.98 | 276715 | 0.045 |
| *H. e. hydaraP* | *optix* | Herato1801 | 1644203 | 78 | 234.88 | 4901 | 0.001 | 1300682 | 46 | 26.83 | 42912 | 0.007 |

| **Population** | **Locus** | **Scaffold** | **Position** | **CLR** | $\alpha$ | **2$N_e s$** | $s$ | **Position ($\alpha_{min}$)** | **CLR ($\alpha_{min}$)** | $\alpha_{min}$ | **2$N_e s$ ($\alpha_{min}$)** | $s$ **($\alpha_{min}$)** |
|---|---|---|---|---|---|---|---|---|---|---|---|---|
| *H.e. lativitta* | *optix* | Herato1801 | 1381486 | 480 | 17.87 | 64400 | 0.01 | 1303132 | 470 | 9.42 | 122169 | 0.019 |
| | | | 1304582 | 478 | 9.44 | 121980 | 0.019 | 1303132 | 470 | 9.42 | 122169 | 0.019 |
| *H. e. notabilis* | *optix* | Herato1801 | 1294528 | 4690 | 3.03 | 370210 | 0.059 | 1305279 | 4604 | 2.99 | 374119 | 0.06 |
| *H. e. venus* | *optix* | Herato1801 | 921654 | 185 | 24.81 | 29867 | 0.007 | 900003 | 79 | 10.97 | 67582 | 0.016 |

**Supplementary Table 9:** Position, composite likelihood-ratio statistics (CLR) and strength of selection ($\alpha$, $2N_es$, and $s$) for the highest CLR and the smallest $\alpha$ value on each background scaffold ($\alpha_{min}$) for *H. erato*. Data are from SweepFinder2[3,4] runs with background site frequency spectrum estimated from background scaffolds.

| Population | Scaffold | Position | CLR | $\alpha$ | $2N_es$ | $s$ | Position ($\alpha_{min}$) | CLR ($\alpha_{min}$) | $\alpha_{min}$ | $2N_es$ ($\alpha_{min}$) | $s$ ($\alpha_{min}$) |
|---|---|---|---|---|---|---|---|---|---|---|---|
| *H. e. amalfreda* | Herato0411 | 4991530 | 21 | 1207.76 | 1158 | 0 | 4298985 | 13 | 51.96 | 26924 | 0.004 |
| *H. e. cyrbiaN* | Herato0411 | 4206446 | 18 | 1408.09 | 451 | 0 | 4303500 | 3 | 72.97 | 8699 | 0.003 |
| *H. e. demophoon* | Herato0411 | 4306907 | 14 | 64.78 | 21670 | 0.003 | 4302807 | 8 | 59.06 | 23769 | 0.004 |
| *H. e. emma* | Herato0411 | 4938642 | 16 | 1762.73 | 858 | 0 | 4303620 | 4 | 58.4 | 25900 | 0.004 |
| *H. e. erato* | Herato0411 | 4346808 | 21 | 163.79 | 8199 | 0.001 | 4301157 | 1 | 56.05 | 23961 | 0.004 |
| *H. e. etylus* | Herato0411 | 4938611 | 20 | 737.55 | 1852 | 0 | 4300691 | 12 | 51.04 | 26762 | 0.004 |
| *H. e. favorinus* | Herato0411 | 4985282 | 12 | 1269.09 | 1192 | 0 | 4346208 | 0 | 315.84 | 4789 | 0.001 |
| *H. e. hydaraFG* | Herato0411 | 4303406 | 28 | 48.89 | 27472 | 0.004 | 4301606 | 26 | 48.17 | 27877 | 0.005 |
| *H. e. hydaraP* | Herato0411 | 4996536 | 19 | 942.33 | 1490 | 0 | 4298358 | 7 | 50.07 | 28036 | 0.004 |
| *H.e. lativitta* | Herato0411 | 4276250 | 14 | 4479.58 | 313 | 0 | 4301551 | 1 | 60.29 | 23277 | 0.004 |
| *H. e. notabilis* | Herato0411 | 4276241 | 15 | 4491.75 | 304 | 0 | 4299742 | 10 | 43.57 | 31350 | 0.005 |
| *H. e. venus* | Herato0411 | 4892869 | 14 | 1623.69 | 556 | 0 | 4296935 | 1 | 190.87 | 4734 | 0.001 |
| *H. e. amalfreda* | Herato0601 | 1736795 | 36 | 245.53 | 4550 | 0.001 | 831240 | 3 | 234.12 | 4771 | 0.001 |
| *H. e. cyrbiaN* | Herato0601 | 1370053 | 68 | 122.11 | 4151 | 0.001 | 1366802 | 65 | 100.36 | 5051 | 0.002 |
| *H. e. demophoon* | Herato0601 | 1517280 | 30 | 516.29 | 2171 | 0 | 1517330 | 29 | 508.59 | 2204 | 0 |
| *H. e. emma* | Herato0601 | 1764992 | 24 | 613.16 | 1970 | 0 | 1354931 | 22 | 300.93 | 4014 | 0.001 |
| *H. e. erato* | Herato0601 | 1313021 | 23 | 945.95 | 1134 | 0 | 1354974 | 11 | 346.54 | 3095 | 0.001 |
| *H. e. etylus* | Herato0601 | 1764630 | 42 | 225.83 | 4830 | 0.001 | 1763930 | 34 | 213.31 | 5113 | 0.001 |
| *H. e. favorinus* | Herato0601 | 1398537 | 32 | 1419.11 | 851 | 0 | 1355285 | 13 | 294.82 | 4097 | 0.001 |
| *H. e. hydaraFG* | Herato0601 | 1650955 | 64 | 382.41 | 2804 | 0 | 831954 | 2 | 246.67 | 4348 | 0.001 |
| *H. e. hydaraP* | Herato0601 | 1398435 | 30 | 1031.75 | 1086 | 0 | 1763700 | 9 | 447.92 | 2503 | 0 |
| *H.e. lativitta* | Herato0601 | 1651545 | 35 | 1311.02 | 855 | 0 | 831454 | 2 | 293.39 | 3820 | 0.001 |
| *H. e. notabilis* | Herato0601 | 1495484 | 22 | 2175.42 | 501 | 0 | 831140 | 0 | 429.41 | 2540 | 0 |
| *H. e. venus* | Herato0601 | 1370059 | 95 | 82.5 | 8746 | 0.002 | 1370259 | 91 | 82.14 | 8784 | 0.002 |
| *H. e. amalfreda* | Herato0821 | 2893797 | 82 | 148.54 | 8876 | 0.001 | 2896497 | 55 | 117.8 | 11192 | 0.002 |
| *H. e. cyrbiaN* | Herato0821 | 2736672 | 45 | 184.13 | 3249 | 0.001 | 2972682 | 24 | 99.09 | 6038 | 0.002 |
| *H. e. demophoon* | Herato0821 | 2900543 | 31 | 703.49 | 1881 | 0 | 2972045 | 12 | 317.23 | 4170 | 0.001 |
| *H. e. emma* | Herato0821 | 2893838 | 50 | 477.6 | 2985 | 0 | 2889088 | 12 | 199.64 | 7140 | 0.001 |
| *H. e. erato* | Herato0821 | 2887040 | 32 | 495.24 | 2556 | 0 | 2889040 | 11 | 182.64 | 6930 | 0.001 |
| *H. e. etylus* | Herato0821 | 2972094 | 13 | 242.72 | 5303 | 0.001 | 2971894 | 12 | 240.33 | 5356 | 0.001 |
| *H. e. favorinus* | Herato0821 | 2885842 | 26 | 476.22 | 2993 | 0 | 2889092 | 13 | 199.31 | 7152 | 0.001 |
| *H. e. hydaraFG* | Herato0821 | 2015021 | 19 | 1657.86 | 763 | 0 | 2499406 | 3 | 399.99 | 3164 | 0.001 |
| *H. e. hydaraP* | Herato0821 | 2738537 | 32 | 331.44 | 3991 | 0.001 | 2897492 | 10 | 247.83 | 5338 | 0.001 |
| *H.e. lativitta* | Herato0821 | 2896842 | 35 | 184.24 | 7179 | 0.001 | 2896542 | 33 | 182.13 | 7262 | 0.001 |
| *H. e. notabilis* | Herato0821 | 2271120 | 10 | 1398.73 | 920 | 0 | 2889095 | 0 | 402.25 | 3200 | 0.001 |
| *H. e. venus* | Herato0821 | 2735284 | 36 | 249.95 | 3407 | 0.001 | 2735434 | 35 | 247.62 | 3439 | 0.001 |

| Population | Scaffold | Position | CLR | $\alpha$ | $2N_es$ | $s$ | Position ($\alpha_{min}$) | CLR ($\alpha_{min}$) | $\alpha_{min}$ | $2N_es$ ($\alpha_{min}$) | $s$ ($\alpha_{min}$) |
|---|---|---|---|---|---|---|---|---|---|---|---|
| *H. e. amalfreda* | Herato1901 | 3171719 | 44 | 406.91 | 2473 | 0 | 2646389 | 4 | 113.64 | 8856 | 0.001 |
| *H. e. cyrbiaN* | Herato1901 | 2846944 | 60 | 193.71 | 2358 | 0.001 | 2646632 | 30 | 68.6 | 6657 | 0.002 |
| *H. e. demophoon* | Herato1901 | 3396907 | 196 | 34.93 | 28911 | 0.004 | 3395457 | 187 | 34.29 | 29448 | 0.005 |
| *H. e. emma* | Herato1901 | 3171790 | 68 | 227.96 | 4773 | 0.001 | 3438258 | 11 | 86.67 | 12554 | 0.002 |
| *H. e. erato* | Herato1901 | 3171726 | 50 | 489.04 | 1976 | 0 | 2646607 | 6 | 129.18 | 7479 | 0.001 |
| *H. e. etylus* | Herato1901 | 3171722 | 84 | 234.95 | 4182 | 0.001 | 3366129 | 51 | 103.61 | 9484 | 0.002 |
| *H. e. favorinus* | Herato1901 | 3171441 | 51 | 333.83 | 3259 | 0 | 2647807 | 5 | 153.49 | 7089 | 0.001 |
| *H. e. hydaraFG* | Herato1901 | 3171813 | 55 | 409.96 | 2357 | 0 | 3367468 | 26 | 88.6 | 10905 | 0.002 |
| *H. e. hydaraP* | Herato1901 | 3396556 | 234 | 25.21 | 40051 | 0.006 | 3396806 | 234 | 25.21 | 40065 | 0.006 |
| *H.e. lativitta* | Herato1901 | 3171698 | 54 | 328.19 | 3076 | 0 | 3366606 | 27 | 128.31 | 7868 | 0.001 |
| *H. e. notabilis* | Herato1901 | 3171700 | 61 | 312.25 | 3147 | 0.001 | 3396758 | 9 | 161.06 | 6101 | 0.001 |
| *H. e. venus* | Herato1901 | 3337404 | 95 | 106.9 | 6081 | 0.001 | 2991894 | 67 | 44.62 | 14567 | 0.003 |

**Supplementary Table 10:** Position, composite likelihood-ratio statistics (CLR) and strength of selection ($\alpha$, $2N_es$, and $s$) for the highest CLR and the smallest $\alpha$ value on each colour pattern scaffold ($\alpha_{min}$) for *H. erato*. Additional relevant peaks on scaffolds are also given. Data are from SweepFinder2[3,4] runs with background site frequency spectrum estimated from background and colour pattern scaffolds.

| Population | Locus | Scaffold | Position | CLR | $\alpha$ | $2N_es$ | $s$ | Position ($\alpha_{min}$) | CLR ($\alpha_{min}$) | $\alpha_{min}$ | $2N_eS$ (αmin) | $s$ ($\alpha_{min}$) |
|---|---|---|---|---|---|---|---|---|---|---|---|---|
| *H. e. amalfreda* | *WntA* | Herato1001 | 4642279 | 434 | 29.58 | 32666 | 0.005 | 4644829 | 411 | 28.92 | 33408 | 0.005 |
| *H. e. cyrbiaN* | *WntA* | Herato1001 | 5466098 | 69 | 155.66 | 2816 | 0.001 | 4402079 | 57 | 72.47 | 6050 | 0.002 |
| *H. e. demophoon* | *WntA* | Herato1001 | 4410429 | 99 | 141.61 | 6846 | 0.001 | 4410479 | 98 | 141.6 | 6847 | 0.001 |
| *H. e. emma* | *WntA* | Herato1001 | 4649730 | 216 | 56.76 | 18404 | 0.003 | 4649680 | 216 | 56.75 | 18406 | 0.003 |
| *H. e. erato* | *WntA* | Herato1001 | 4642175 | 361 | 33.42 | 27754 | 0.004 | 4644875 | 332 | 32.74 | 28331 | 0.005 |
| *H. e. etylus* | *WntA* | Herato1001 | 4645879 | 511 | 20.92 | 45087 | 0.007 | 4685579 | 313 | 13.76 | 68569 | 0.011 |
| *H. e. favorinus* | *WntA* | Herato1001 | 5465699 | 150 | 122.65 | 8517 | 0.001 | 4649330 | 93 | 48.77 | 21419 | 0.003 |
| *H. e. hydaraFG* | *WntA* | Herato1001 | 4668834 | 166 | 41.74 | 22222 | 0.004 | 4668134 | 160 | 41.18 | 22522 | 0.004 |
| *H. e. hydaraP* | *WntA* | Herato1001 | 5302745 | 92 | 215.84 | 4492 | 0.001 | 4719285 | 21 | 120.98 | 8013 | 0.001 |
| *H.e. lativitta* | *WntA* | Herato1001 | 4651634 | 293 | 46.52 | 20833 | 0.003 | 4649634 | 275 | 45.42 | 21339 | 0.003 |
| *H. e. notabilis* | *WntA* | Herato1001 | 4642524 | 554 | 22.37 | 42160 | 0.007 | 4685525 | 249 | 15.91 | 59304 | 0.009 |
| *H. e. venus* | *WntA* | Herato1001 | 4405179 | 119 | 57.41 | 10869 | 0.003 | 4403229 | 94 | 53.93 | 11571 | 0.003 |
| *H. e. amalfreda* | *cortex* | Herato1505 | 2494717 | 1306 | 14.75 | 98269 | 0.015 | 2499417 | 1224 | 14.63 | 99040 | 0.015 |
| *H. e. cyrbiaN* | *cortex* | Herato1505 | 2453566 | 167 | 47.95 | 13714 | 0.004 | 2131107 | 132 | 16.5 | 39848 | 0.013 |
| *H. e. demophoon* | *cortex* | Herato1505 | 2277009 | 723 | 20.21 | 71962 | 0.011 | 2267559 | 704 | 19.84 | 73287 | 0.011 |
| *H. e. emma* | *cortex* | Herato1505 | 2496694 | 1120 | 18.13 | 86428 | 0.013 | 2497944 | 1110 | 18.07 | 86717 | 0.013 |
| *H. e. erato* | *cortex* | Herato1505 | 2493705 | 1151 | 16.7 | 83292 | 0.013 | 2491005 | 1134 | 16.67 | 83477 | 0.014 |
| *H. e. etylus* | *cortex* | Herato1505 | 2494192 | 1016 | 17.5 | 80873 | 0.013 | 2497192 | 986 | 17.43 | 81167 | 0.013 |
| *H. e. favorinus* | *cortex* | Herato1505 | 2496137 | 1734 | 9.92 | 157969 | 0.023 | 2494937 | 1725 | 9.91 | 158072 | 0.023 |
| *H. e. hydaraFG* | *cortex* | Herato1505 | 2493808 | 1107 | 15.64 | 88979 | 0.014 | 2488308 | 1081 | 15.31 | 90896 | 0.015 |
| *H. e. hydaraP* | *cortex* | Herato1505 | 2985526 | 205 | 35.05 | 41484 | 0.006 | 2985526 | 205 | 35.05 | 41484 | 0.006 |
| *H.e. lativitta* | *cortex* | Herato1505 | 2491914 | 1030 | 17.92 | 81126 | 0.013 | 2490014 | 1018 | 17.85 | 81455 | 0.013 |
| *H. e. notabilis* | *cortex* | Herato1505 | 2497600 | 909 | 18.52 | 76386 | 0.012 | 2501750 | 879 | 17.92 | 78973 | 0.013 |
| | | | 1963287 | 311 | 60.24 | 23489 | 0.004 | 1962437 | 301 | 59.92 | 23612 | 0.004 |
| *H. e. venus* | *cortex* | Herato1505 | 2069802 | 429 | 9.57 | 97818 | 0.023 | 2130154 | 252 | 8.34 | 112198 | 0.026 |
| *H. e. amalfreda* | *optix* | Herato1801 | 1375134 | 663 | 15.79 | 72638 | 0.011 | 1303280 | 544 | 8.22 | 139501 | 0.022 |
| | | | 1304780 | 552 | 8.23 | 139362 | 0.022 | 1303280 | 544 | 8.22 | 139501 | 0.022 |
| *H. e. cyrbiaN* | *optix* | Herato1801 | 916554 | 102 | 74.8 | 6960 | 0.002 | 921655 | 96 | 49.41 | 10537 | 0.003 |
| *H. e. demophoon* | *optix* | Herato1801 | 938505 | 47 | 408.01 | 2822 | 0 | 1293521 | 4 | 117.67 | 9784 | 0.002 |
| *H. e. emma* | *optix* | Herato1801 | 1381539 | 287 | 39.53 | 31378 | 0.005 | 1303933 | 216 | 16.34 | 75896 | 0.011 |
| *H. e. erato* | *optix* | Herato1801 | 1380328 | 672 | 14.57 | 75574 | 0.012 | 1305325 | 467 | 9.61 | 114604 | 0.019 |
| | | | 1303775 | 472 | 9.62 | 114444 | 0.019 | 1305325 | 467 | 9.61 | 114604 | 0.019 |
| *H. e. etylus* | *optix* | Herato1801 | 1382440 | 261 | 32.26 | 34721 | 0.006 | 1300235 | 172 | 15.3 | 73220 | 0.012 |
| | | | 1305285 | 193 | 15.56 | 71968 | 0.011 | 1300235 | 172 | 15.3 | 73220 | 0.012 |
| *H. e. favorinus* | *optix* | Herato1801 | 1381333 | 142 | 44.88 | 27639 | 0.004 | 1304029 | 55 | 37.14 | 33398 | 0.005 |
| | | | 1251876 | 88 | 139.79 | 8874 | 0.001 | 1304029 | 55 | 37.14 | 33398 | 0.005 |
| *H. e. hydaraFG* | *optix* | Herato1801 | 1425389 | 2890 | 4.95 | 222666 | 0.036 | 1429539 | 2881 | 4.94 | 223002 | 0.036 |
| *H. e. hydaraP* | *optix* | Herato1801 | 1643303 | 55 | 356.23 | 3232 | 0.001 | 1305282 | 5 | 70.22 | 16395 | 0.003 |

| Population | Locus | Scaffold | Position | CLR | *α* | 2$N_e$*s* | *s* | Position ($α_{min}$) | CLR ($α_{min}$) | $α_{min}$ | 2$N_e$*s* (αmin) | *s* ($α_{min}$) |
|---|---|---|---|---|---|---|---|---|---|---|---|---|
| *H.e. lativitta* | *optix* | Herato1801 | 1381286 | 350 | 24.23 | 47504 | 0.007 | 1302982 | 247 | 12.27 | 93822 | 0.015 |
| | | | 1304732 | 258 | 12.31 | 93489 | 0.015 | 1302982 | 247 | 12.27 | 93822 | 0.015 |
| *H. e. notabilis* | *optix* | Herato1801 | 1293428 | 2938 | 3.74 | 299762 | 0.048 | 1305629 | 2836 | 3.67 | 305008 | 0.049 |
| *H. e. venus* | *optix* | Herato1801 | 1067013 | 94 | 89.46 | 8283 | 0.002 | 1300277 | 9 | 30.24 | 24506 | 0.006 |

**Supplementary Table 11:** Position, composite likelihood-ratio statistics (CLR) and strength of selection ($\alpha$, $2N_es$, and $s$) for the highest CLR and the smallest $\alpha$ value on each background scaffold ($\alpha_{min}$) for *H. erato*. Data are from SweepFinder2[3,4] runs with background site frequency spectrum estimated from background and colour pattern scaffolds.

| Population | Scaffold | Position | CLR | $\alpha$ | $2N_es$ | $s$ | Position ($\alpha_{min}$) | CLR ($\alpha_{min}$) | $\alpha_{min}$ | $2N_es$ ($\alpha_{min}$) | $s$ ($\alpha_{min}$) |
|---|---|---|---|---|---|---|---|---|---|---|---|
| *H. e. amalfreda* | Herato0411 | 4991530 | 16 | 1371.79 | 1020 | 0 | 4300335 | 2 | 69.4 | 20157 | 0.003 |
| *H. e. cyrbiaN* | Herato0411 | 4206446 | 13 | 1604.88 | 396 | 0 | 4308951 | 0 | 188.29 | 3371 | 0.001 |
| *H. e. demophoon* | Herato0411 | 4990346 | 8 | 5043.62 | 278 | 0 | 4251454 | 3 | 444 | 3162 | 0 |
| *H. e. emma* | Herato0411 | 4938642 | 13 | 2186.33 | 692 | 0 | 4305120 | 0 | 186.18 | 8124 | 0.001 |
| *H. e. erato* | Herato0411 | 4346608 | 11 | 191.65 | 7007 | 0.001 | 4303307 | 1 | 180.59 | 7437 | 0.001 |
| *H. e. etylus* | Herato0411 | 4938611 | 14 | 1015.4 | 1345 | 0 | 4305191 | 3 | 81.76 | 16705 | 0.003 |
| *H. e. favorinus* | Herato0411 | 4985282 | 8 | 1409.71 | 1073 | 0 | 4862378 | 1 | 600.3 | 2520 | 0 |
| *H. e. hydaraFG* | Herato0411 | 4326307 | 14 | 2276.13 | 590 | 0 | 4302656 | 2 | 66.65 | 20150 | 0.003 |
| *H. e. hydaraP* | Herato0411 | 4271057 | 14 | 1313.24 | 1069 | 0 | 4307208 | 2 | 162.05 | 8662 | 0.001 |
| *H.e. lativitta* | Herato0411 | 4276250 | 12 | 5194.78 | 270 | 0 | 4306401 | 1 | 183.19 | 7661 | 0.001 |
| *H. e. notabilis* | Herato0411 | 4202989 | 11 | 8102.94 | 169 | 0 | 4311093 | 0 | 402.78 | 3391 | 0.001 |
| *H. e. venus* | Herato0411 | 4892869 | 11 | 1942.13 | 465 | 0 | 4766762 | 5 | 240.52 | 3757 | 0.001 |
| *H. e. amalfreda* | Herato0601 | 1398524 | 24 | 1608.56 | 694 | 0 | 1736845 | 18 | 302.21 | 3696 | 0.001 |
| *H. e. cyrbiaN* | Herato0601 | 1370053 | 39 | 178.88 | 2834 | 0.001 | 1366752 | 28 | 158.91 | 3190 | 0.001 |
| *H. e. demophoon* | Herato0601 | 1517230 | 21 | 688.49 | 1628 | 0 | 1517330 | 20 | 668.58 | 1677 | 0 |
| *H. e. emma* | Herato0601 | 1398482 | 18 | 3321.79 | 364 | 0 | 1354881 | 15 | 334.98 | 3606 | 0.001 |
| *H. e. erato* | Herato0601 | 1313021 | 19 | 1029.58 | 1042 | 0 | 1354924 | 6 | 413.26 | 2595 | 0 |
| *H. e. etylus* | Herato0601 | 1764630 | 28 | 289.3 | 3770 | 0.001 | 1763880 | 17 | 259.31 | 4206 | 0.001 |
| *H. e. favorinus* | Herato0601 | 1398537 | 25 | 1625.82 | 743 | 0 | 1355285 | 2 | 367.87 | 3283 | 0 |
| *H. e. hydaraFG* | Herato0601 | 1650955 | 46 | 472.99 | 2267 | 0 | 1650005 | 32 | 357.85 | 2997 | 0 |
| *H. e. hydaraP* | Herato0601 | 1398435 | 23 | 1204.59 | 931 | 0 | 1763450 | 9 | 575.82 | 1947 | 0 |
| *H.e. lativitta* | Herato0601 | 1651545 | 30 | 1450.78 | 772 | 0 | 831854 | 1 | 364.85 | 3072 | 0 |
| *H. e. notabilis* | Herato0601 | 1495484 | 16 | 2463.33 | 443 | 0 | 1353925 | 2 | 852.72 | 1279 | 0 |
| *H. e. venus* | Herato0601 | 1370059 | 49 | 117.58 | 6137 | 0.001 | 1370159 | 48 | 117.29 | 6152 | 0.001 |
| *H. e. amalfreda* | Herato0821 | 2893797 | 62 | 208.41 | 6326 | 0.001 | 2896597 | 34 | 139.75 | 9434 | 0.001 |
| *H. e. cyrbiaN* | Herato0821 | 2736672 | 28 | 220.18 | 2717 | 0.001 | 2736722 | 28 | 220.07 | 2718 | 0.001 |
| *H. e. demophoon* | Herato0821 | 2900543 | 22 | 804.48 | 1644 | 0 | 2972095 | 4 | 390.28 | 3390 | 0.001 |
| *H. e. emma* | Herato0821 | 2893838 | 43 | 514.73 | 2769 | 0 | 2889038 | 2 | 236.65 | 6023 | 0.001 |
| *H. e. erato* | Herato0821 | 2887040 | 28 | 567.55 | 2230 | 0 | 2896540 | 8 | 230.79 | 5484 | 0.001 |
| *H. e. etylus* | Herato0821 | 2893790 | 9 | 2444.68 | 527 | 0 | 2971794 | 2 | 336.69 | 3823 | 0.001 |
| *H. e. favorinus* | Herato0821 | 2885842 | 19 | 589.22 | 2419 | 0 | 2888292 | 6 | 287.11 | 4965 | 0.001 |
| *H. e. hydaraFG* | Herato0821 | 2015021 | 14 | 1921.28 | 659 | 0 | 2499056 | 0 | 634.99 | 1993 | 0 |
| *H. e. hydaraP* | Herato0821 | 2738537 | 20 | 508.65 | 2601 | 0 | 2736737 | 11 | 384.27 | 3443 | 0.001 |
| *H.e. lativitta* | Herato0821 | 2896842 | 24 | 207.54 | 6373 | 0.001 | 2896592 | 23 | 205.36 | 6440 | 0.001 |
| *H. e. notabilis* | Herato0821 | 2115614 | 7 | 3396.92 | 379 | 0 | 2888345 | 1 | 734.68 | 1752 | 0 |
| *H. e. venus* | Herato0821 | 2735284 | 23 | 289.64 | 2940 | 0.001 | 2735434 | 22 | 287.56 | 2961 | 0.001 |

| Population | Scaffold | Position | CLR | $\alpha$ | $2N_es$ | $s$ | Position ($\alpha_{min}$) | CLR ($\alpha_{min}$) | $\alpha_{min}$ | $2N_es$ ($\alpha_{min}$) | $s$ ($\alpha_{min}$) |
|---|---|---|---|---|---|---|---|---|---|---|---|
| *H. e. amalfreda* | Herato1901 | 3171719 | 35 | 466.22 | 2159 | 0 | 2648039 | 2 | 171.54 | 5867 | 0.001 |
| *H. e. cyrbiaN* | Herato1901 | 2846944 | 38 | 231.72 | 1971 | 0.001 | 2647282 | 9 | 86.68 | 5268 | 0.002 |
| *H. e. demophoon* | Herato1901 | 3397207 | 126 | 42.94 | 23520 | 0.004 | 3395757 | 119 | 41.88 | 24111 | 0.004 |
| *H. e. emma* | Herato1901 | 3171790 | 56 | 338.86 | 3211 | 0 | 3437758 | 3 | 98.34 | 11064 | 0.002 |
| *H. e. erato* | Herato1901 | 3171726 | 42 | 539.69 | 1790 | 0 | 2647507 | 2 | 176.06 | 5488 | 0.001 |
| *H. e. etylus* | Herato1901 | 3171722 | 65 | 268.26 | 3663 | 0.001 | 3366479 | 30 | 132.22 | 7431 | 0.001 |
| *H. e. favorinus* | Herato1901 | 3172741 | 33 | 942.19 | 1155 | 0 | 3366954 | 16 | 259.96 | 4186 | 0.001 |
| *H. e. hydaraFG* | Herato1901 | 3171813 | 39 | 486.51 | 1986 | 0 | 3366818 | 3 | 153.76 | 6283 | 0.001 |
| *H. e. hydaraP* | Herato1901 | 3396656 | 155 | 31.3 | 32265 | 0.005 | 3396656 | 155 | 31.3 | 32265 | 0.005 |
| *H.e. lativitta* | Herato1901 | 3171698 | 45 | 359.63 | 2807 | 0 | 3366706 | 15 | 153.9 | 6560 | 0.001 |
| *H. e. notabilis* | Herato1901 | 3171700 | 41 | 364.14 | 2698 | 0 | 3366257 | 12 | 302.85 | 3245 | 0.001 |
| *H. e. venus* | Herato1901 | 3337404 | 62 | 143.11 | 4542 | 0.001 | 2646034 | 16 | 72.63 | 8950 | 0.002 |
